# Convergent state-control of endogenous opioid analgesia

**DOI:** 10.1101/2025.01.03.631111

**Authors:** Blake A. Kimmey, Lindsay Ejoh, Lily Shangloo, Jessica A. Wojick, Samar Nasser Chehimi, Nora M. McCall, Corinna S. Oswell, Malaika Mahmood, Lite Yang, Vijay K. Samineni, Charu Ramakrishnan, Karl Deisseroth, Richard C. Crist, Benjamin C. Reiner, Lin Tian, Gregory Corder

## Abstract

Pain is a dynamic and nonlinear experience shaped by injury and contextual factors, including expectations of future pain or relief^1^. While µ opioid receptors are central to the analgesic effects of opioid drugs, the endogenous opioid neurocircuitry underlying pain and placebo analgesia remains poorly understood. The ventrolateral column of the posterior periaqueductal gray is a critical hub for nociception and endogenous analgesia mediated by opioid signaling^2^. However, significant gaps remain in understanding the cell-type identities, the sub-second neural dynamics involved in pain modulation, the role of endogenous peptide neuromodulators, and the contextual factors influencing these processes. Using spatial mapping with single-nuclei RNA sequencing of pain-active neurons projecting to distinct long-range brain targets, alongside cell type-specific and activity-dependent genetic tools for *in vivo* optical recordings and modulation of neural activity and opioid peptide release, we identified a functional dichotomy in the ventrolateral periaqueductal gray. Neurons expressing µ opioid receptors encode active nociceptive states, whereas enkephalin-releasing neurons drive pain relief during recovery from injury, in response to learned fear predictions, and during placebo analgesia. Finally, by leveraging the functional effects of placebo analgesia, we used direct optogenetic activation of vlPAG enkephalin neurons to drive opioid peptide release, resulting in a robust reduction in pain. These findings show that diverse need states converge on a shared midbrain circuit that releases endogenous opioids with high spatiotemporal precision to suppress nociceptive activity and promote analgesia.

## INTRODUCTION

The aversive experience of pain entails a large, interconnected brain network, with diverse functions in sensory, affective, motivational, cognitive and homeostatic processes that collectively generate behavioral repertoires to limit and avoid harm^1,3^. Thus, pain is shaped not only by the immediate noxious event, but also the context in which it is experienced and the learned expectations surrounding how and when nociceptive processes will be engaged. The midbrain periaqueductal gray (PAG) has emerged as a central hub of the neural pain network that facilitates acute nociceptive signal processing and pain prediction error. The PAG is divided into functional domains, or columns, surrounding the cerebral aqueduct^4,5^ that are locally connected^6^, but differentially engaged to promote survival and recovery in the face threat, fear, and pain, among others^7^.

In the context of injury, the lateral (lPAG) and ventrolateral columns (vlPAG) receive direct ascending nociceptive signals from the spinal cord^8^ and descending cortical and subcortical inputs^9–11^ that facilitate ongoing pain modulation^12^. The convergence of pain information in the vlPAG is further modified by µ opioid receptors (MORs)^13^ engaged by exogenous opioid drugs and endogenous MOR agonists, primarily met- and leu-enkephalin peptides^14–18^. For example, both direct microinfusion of MOR agonists^19–21^ and enkephalinase inhibitor into posterior vlPAG^22^ produced robust antinociception that could be blocked with MOR antagonist. This suggests that endogenous enkephalins released in vlPAG inhibit MOR-expressing neurons and thereby activate the canonical descending disinhibition pathway to rostral ventromedial medulla (RVM) to produce analgesia via indirect control of the dorsal horn^12^. Foundational work supported excitatory, analgesic connectivity between PAG and RVM as microinfusion of glutamate into the vlPAG increased RVM cellular activity and reduced nociceptive reflexive paw withdrawal^23^. Likewise, broad chemogenetic^24^ or optogenetic^25^ stimulation of vlPAG neurons revealed that activation of vlPAG glutamatergic neurons is generally antinociceptive, while GABAergic neurons activation is pronociceptive^24,26^. However, MOR-expressing vlPAG cells are a mixed population with both ascending and descending projection populations that express glutamatergic and GABAergic markers^27,28^, indicating that MOR activation via endogenous enkephalin release in vlPAG may produce analgesia through a complex interplay of cell-types and functional connectivity that has not been captured in previous investigations, requiring new tools and approaches to uncover.

Here, we combine transcriptomic, molecular, and *in vivo* imaging-based approaches to reveal the nuanced and dynamic attributes of opioidergic circuitry in the vlPAG. For the first time, we show that ventrolateral PAG MOR+ neurons undergo experience-related modulation that contrasts with the fall and rise of released enkephalin using novel MOR promoter-driven fluorescent biosensors. Critically, we find that the interplay of MOR-expressing neural activity and enkephalin signaling is mutable according to learned expectations surrounding pain experience and uncovered the vlPAG enkephalinergic population as a possible driver of these effects.

## RESULTS

### Transcriptomic and projection profiling of nociceptive opioidergic cell types in the periaqueductal gray

The PAG is a critical site of nociceptive and antinociceptive signal processing and opioid-mediated analgesia. However, we still lack important information regarding the spatial and transcriptomic profile of the cells that may underlie those pain-related behavioral functions. We first investigated the anterior-posterior distribution of nociceptive PAG cells to determine coordinates of maximal noxious stimulus-related activation. To do this, we used the targeted recombination in active populations (*i*.*e*., TRAP2) transgenic mouse line^29^ crossed to the Cre-dependent reporter mouse line Ai9 for nociceptive or home-cage neural tagging (i.e. *nox*TRAP or *hc*TRAP, respectively) with the red fluorescent protein tdTomato. *Nox*TRAP is achieved via repeated noxious 55 °C water drops applied to the left hindpaw to trigger the immediate early gene (IEG) promoter for *Fos* to induce expression of 4-hydroxytamoxifen-inducible Cre recombinase (**Fig. 1a**). Cells expressing tdTomato were found throughout all PAG columns and across the entire anterior-posterior domain that was assessed (**Fig. 1b**). Significant differences between the *hc*TRAP (n=3 females) and *nox*TRAP (n=3; 1 female, 2 males) groups was found at coordinates −4.60 mm and −4.72 mm relative to Bregma when comparing total PAG counts (**Extended Data Fig. 1a-b**). Analysis of the ventrolateral column (vlPAG) revealed significant differences between *nox*TRAP mice and *hc*TRAP controls at posterior coordinates (**Extended Data Fig. 1c**); significance stars denote vlPAG comparisons (**Fig. 1b**). Based on these data, we identified the posterior vlPAG, beginning at −4.60 mm relative to Bregma, as a highly nociceptive region when compared to no-stimulus *hc*TRAP control mice.

**Figure 1.**
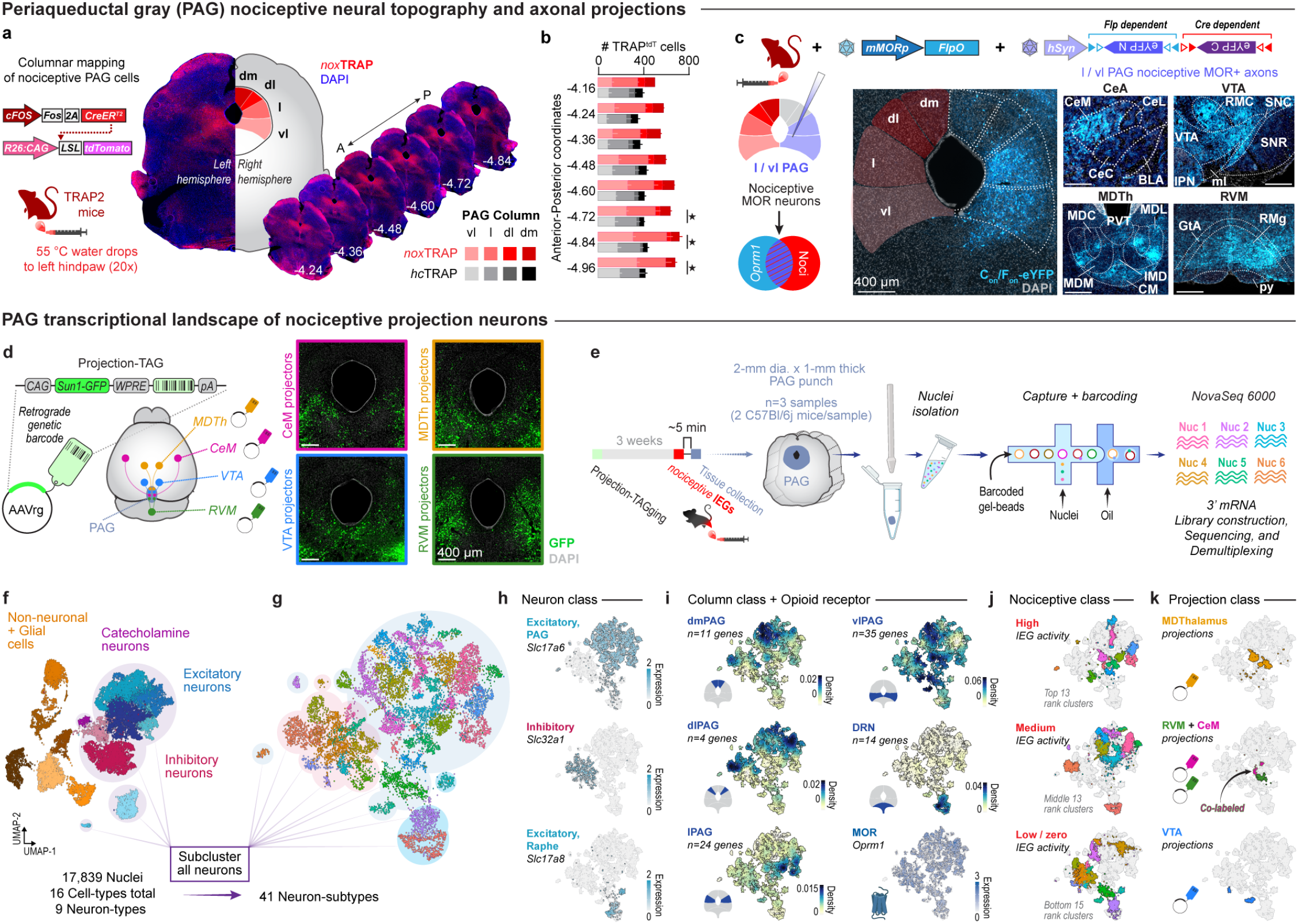
Noxious stimulation recruits transcriptionally-defined opioidergic efferent circuits arising in the ventrolateral PAG. **a**. Didactic (left) of noxious stimulus-induced targeted recombination in active populations (*nox*TRAP) procedure performed with TRAP2:Ai9 mice to permanently label nociceptive PAG cells with tdTomato fluorophore. The *nox*TRAP was achieved by repeated application of noxious 55° C water to the left hindpaw and administration of 4-hydroxy tamoxifen (4-OHT); unstimulated control mice received 4-OHT in the home cage (*hc*TRAP). Representative 4x images of anterior to posterior periaqueductal gray (PAG) sections demonstrating robust *nox*TRAP-related tdTomato labeling across columns of the PAG (right). **b**. Quantification of tdTomato-positive cells (TRAP^tdT^) within each PAG column between Control and 55° C water-stimulated groups; statistically significant between-groups comparisons are shown for the vl column. **c**. Methods for achieving intersectional labeling of nociceptive lPAG and vlPAG cells for output mapping (top). TRAP2 mice received a combination of AAVs expressing FlpO under control of the mMOR promoter and Cre/Flp-dependent eYFP under the hSyn promoter prior to the *nox*TRAP procedure (left). Representative 4x images (bottom right) demonstrate eYFP-expressing cells in blue in lPAG and vlPAG and their axonal projections in the medial nucleus of the central amygdala (CeM), ventral tegmental area (VTA), mediodorsal thalamic nuclei (MDTh), and rostral ventromedial medulla (RVM). **d**. Experimental design for injecting uniquely-barcoded retrograde-AAVs (i.e., Projection-TAGs) into four key nociceptive regions innervated by PAG: CeM, MDTh, VTA, and RVM (left). The Projection-TAG viruses also express nuclear localized Sun1-GFP in PAG in four separate mice (right). **e**. Three weeks after Projection-TAGging, mice received 55° C water drops to evoke noxious stimulus-related IEG expression in PAG followed by rapid dissection and tissue collection of the PAG using a 2-mm inner diameter micro-punch, followed by rapid freezing. Then, samples were prepared for single nucleus RNA sequencing. **f**. Uniform Manifold Approximation and Projection (UMAP) of all nuclei (left; n=17,839) captured by PAG punches in 16 unique clusters, 9 of which were neuronal. g. UMAP of the neuronal sub-clusters (right; n=41) separated into excitatory (blue shading) and inhibitory (red shading) classes. **h**. Feature plots of select genes representing PAG excitatory cell types (top; *Vglut2*), PAG inhibitory cell types (middle; *Vgat*), and dorsal raphe excitatory cell types (bottom; *Vglut3*). **i**. Density feature plots of gene groupings representative of each PAG column, dorsal raphe, and a feature plot of the µ opioid receptor (*Oprm1*). **j**. IEG expression was determined using a modular activity score and mapped to sub-clusters with high (top), medium (middle), and low or zero scores (bottom). **k**. Feature plots of neuronal sub-clusters containing Projection-TAGs.

Recent evidence demonstrates that MOR-expressing cells in vlPAG project broadly to hindbrain and forebrain structures, including the RVM, thalamus, and hypothalamus^28^. However, there are no studies to date that have examined the projection targets of the acutely nociceptive MOR-expressing PAG population. Therefore, we used our recently described synthetic mouse mu-opioid receptor promoter (*mMORp*) viral vectors^30^ to gain genetic access of the lPAG and vlPAG MOR+ population by cloning the FlpO recombinase sequence into the *mMORp* vector for AAV-mediated delivery of FlpO directly into l/vlPAG neurons (AAV-*mMORp*-Flp). By combining this viral approach with the TRAP2 mouse, we achieved combinatorial transduction of a Cre-dependent/Flp-dependent eYFP-expressing AAV (AAV8-*hSyn*-C_ON_/F_ON_-eYFP) in nociceptive MOR-expressing neurons. This INTRSECT^31^ approach revealed diffuse axonal projections from l/vlPAG nociceptive, MOR+ neurons, with relatively denser GFP+ fibers occurring in key ascending and descending nociceptive regions in a representative male mouse, including the medial nucleus of the central amygdala CeM, mediodorsal thalamus (MDTh), ventral tegmental area (VTA), RVM, among other regions with GFP expression (**Fig. 1c**; **Extended Data Fig. 1d**). These results suggest that the acutely nociceptive MOR-expressing cells in the l/vlPAG send extensive brain-wide projections that can influence the perception of pain and engagement in affective-motivational behaviors.

Next, we sought to elucidate the unique genetic profiles of nociceptive PAG projection neurons relative to all PAG and dorsal raphe nucleus (DRN) neurons. Several efforts have been made to characterize behaviorally-relevant PAG cell types at the RNA level, using highly multiplexed fluorescent in situ hybridization (i.e. MERFISH) and RNA sequencing-based approaches^32–36^. Here, we identify the PAG nociceptive cell types that co-express opioid gene markers while ascertaining the identity of several of the ascending and descending projection populations that emerged in our output tracing. We combined two techniques to: 1) obtain the transcriptional profile of all nociceptive PAG cell types and their overlap with opioid gene markers using single-nucleus RNA sequencing (snRNAseq; **Extended Data Fig. 2a-d**); and 2) map the output architecture of PAG cells to four key downstream structures, the CeM^37–39^, MDTh^40,41^, VTA^42–45^, and RVM^12,46^, that are implicated in neural pain processing using a novel retrograde adeno-associated viral approach to express exogenous barcode sequence elements and nuclear localized-GFP in the PAG in a projection-specific manner^47^ (**Fig. 1d**; **Extended Data Fig. 2e-h**). Next, 1-mm punches of the PAG were collected (n=3 biological replicates with 2 male mice combined for each replicate) from mice that were exposed to noxious 55° C water drops. The noxious water stimulation occurred 5-min prior to tissue collection to evoke IEG expression for “activity scoring” and IEG density mapping^48–50^. PAG samples were processed using the 10X Genomics 3’ gene expression assay (**Fig. 1e**). All PAG samples were collected and processed on the same day to avoid batch effects. After quality-control assessment, we analyzed 17,839 single nuclei and identified 16 distinct cell type clusters across all expected major cell classes (e.g. neurons, microglia, astrocytes, etc.; **Fig. 1f**).

Based on known cell class marker genes, we separated the clusters into 6 neuronal glutamatergic, 2 neuronal GABAergic, 1 neuronal catecholaminergic, and 7 non-neuronal clusters (**Extended Data Fig. 3a-b**). The neuronal clusters were further subclustered into 41 neuronal subtypes that were further examined according to top differentiating genes, opioid receptor and peptide genes, presence of Projection-TAGs, and IEG activity (**Fig. 1g**; **Extended Data Fig. 3c-g**). Broadly, excitatory PAG cell types were identified by the expression of *Slc17a6* (*Vglut2*), while inhibitory PAG cell types were marked by *Slc32a1* (*Vgat*); in the dorsal raphe, excitatory cell types were distinguished by the expression of *Slc17a8* (*Vglut3*) (**Fig. 1h**). We further parsed gene expression patterns by plotting gene groupings representative of each PAG column and the dorsal raphe with density mapping (**Fig. 1i**; **Extended Data Fig. 4**^51^), including a focused analysis of *Oprm1*, the gene for MOR, which showed a diffuse localization pattern (**Fig. 1i**; **Extended Data Fig. 5**^51^). All Projection-TAGs overlapped with vlPAG subclusters and excitatory neuronal subtypes, with one population of VTA-projecting neurons expressing the inhibitory marker *Gad2*. Subclusters containing Projection-TAGs for CeM, MDTh, and RVM exhibited elevated IEG modular activity scores (**Fig. 1j-k**; **Extended Data Fig. 3g**), suggesting that these neural populations are particularly activated by noxious stimulation. Correspondingly, these Projection-TAG subclusters expressed several genes that have been implicated in pain processes in the PAG, including *Cnr1* (CB1 receptor)^52^, *Tacr1* (NK1 receptor)^53^, and *Oprm1* (MOR)^19^ (**Extended Data Fig. 6**^54^). Interestingly, while all Projection-TAG-expressing nuclei expressed some level of *Oprm1*, the barcode corresponding to the MDTh projection population showed the highest degree of overlap with *Oprm1*-expressing neuronal subtypes. Although some genetic commonalities emerged among the Projection-TAG populations, only one neuronal subcluster possessed dual barcode expression representing CeM and RVM projectors. This demonstrates that at least one population of l/vlPAG neurons collateralizes to fore- and hindbrain structures, complementing a prior finding that showed collateralization from single vlPAG neurons to nucleus accumbens and RVM^55^. Collectively, our molecular and projection phenotyping of the PAG provides the first comprehensive map of the genetic, functional, and efferent organization of the PAG and its surrounding regions at the level of single nuclei and reveals multiple projection populations interacting with affective-motivational pain-encoding regions that are under endogenous opioid control.

#### Nociceptive landscape and functional dynamics of MOR-expressing PAG cells

Although we have long known that MOR is expressed in PAG^19,56^, the distribution of MOR-expressing cells in PAG and their functional activation by noxious stimulation remains relatively unexplored. Using fluorescent in situ hybridization (FISH), we determined the columnar organization of *Oprm1* mRNA in the highly nociceptive posterior PAG (i.e. approximately −4.60 to −4.84 mm relative to Bregma) and its overlap with the IEG *Fos* mRNA in either home cage-unstimulated controls (n=3 male mice) or noxious 55 °C water-stimulated mice (n=4 male mice; **Fig. 2a**). Across all mice, vlPAG contained the most *Oprm1*-positive cells relative to the other columns (**Fig. 2b, left**). However, while greater than 50% of all cells within a column expressed *Oprm1*, dorsomedial and dorsolateral PAG contained larger fractions of *Oprm1*-positive cells than lateral and ventrolateral PAG (**Fig. 2b, right**). When comparing the number of *Oprm1* transcripts in PAG and across each column, noxious 55 °C water drops significantly increased the proportion of cells with high (10-14) or very high (15+) numbers of *Oprm1* transcripts while decreasing low (1-4) expressing cells when compared to home cage controls (**Extended Data Fig. 7a-c**). Noxious 55 °C water drop application to the hindpaw substantially elevated total *Fos* transcript levels in the PAG. This effect was most prominent in both the lateral and ventrolateral columns (**Fig. 2c, left**). The proportion of neurons expressing very high (15+) numbers of *Fos* transcript was significantly elevated after noxious water drop application when compared to controls, particularly in vlPAG (**Extended Data Fig. 7d-f**). Columnar overlap of *Oprm1* and *Fos* was near or greater than 50% in both groups, however, in each column this overlap was increased in the noxious-stimulated group relative to controls (**Fig. 2c, right**). These results demonstrate that *Oprm1*-expressing cells across the PAG, primarily in the lPAG and vlPAG, are activated by noxious stimulus exposure beyond background levels of *Fos* expression.

**Figure 2.**
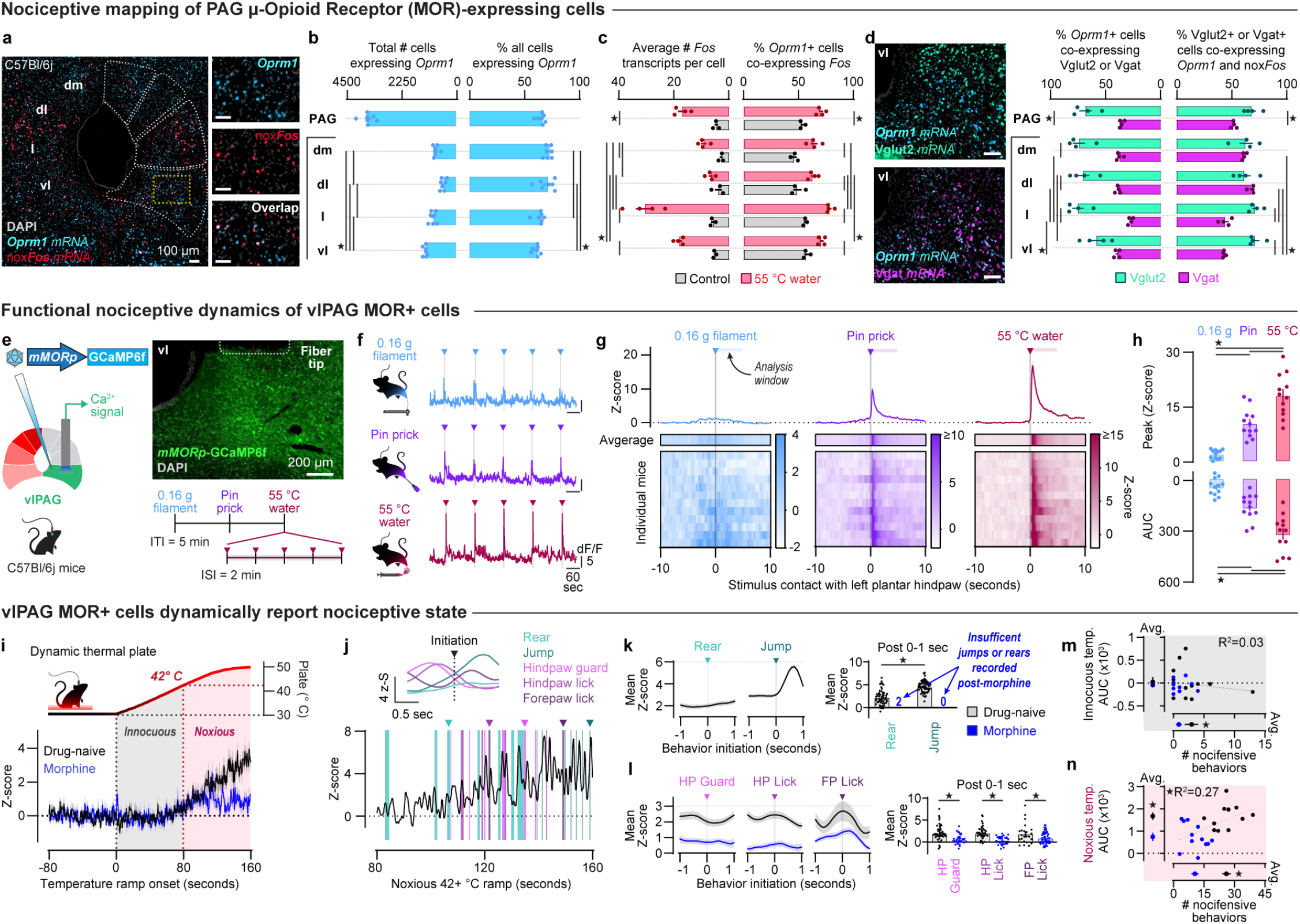
The vlPAG MOR-expressing population represents pain and affective-motivational behavioral states. **a**. Representative 20x fluorescence *in situ* hybridization images demonstrating mRNA transcripts for *Fos* (red) and *Oprm1* (light blue) in vlPAG at approximately A/P −4.72 relative to Bregma. **b**. Quantification of total number of cells expressing *Oprm1* transcript (left) and percent of all cells detected expressing *Oprm1* (right) in the PAG and between columns. **c**. Full PAG and columnar comparisons of *Fos* transcripts per cell (left) and percent of *Oprm1*+ cells co-expressing *Fos* (right) between Control and 55° C water-stimulated mice. **d**. Representative 20x fluorescence *in situ* hybridization images demonstrating mRNA transcripts for *Oprm1* (light blue), *Vglut2* (green; left, top) and *Vgat* (magenta; left, bottom) in vlPAG at approximately A/P −4.72 relative to Bregma. Quantification of *Oprm1*+ cells co-expressing *Vglut2* or *Vgat* (middle) and overlap of *Vglut2* or *Vgat* with both *Oprm1* and nox*Fos* (right) with respect to the PAG and individual columns. **e**. Fiber photometry recordings were performed in the right vlPAG of mice expressing GCaMP6f under the *mMOR* promoter. A battery of 3 distinct stimuli were delivered to the left hindpaw; each stimulus was presented 5 times with a 2-miniuted inter-stimulus interval. **f**. Representative traces from a single mouse demonstrating innocuous and noxious stimulus-evoked transients. **g**. Group average traces and individual mouse heatmaps showing evoked responses to each stimulus modality (i.e., 0.16 g von Frey filament, 25-gauge needle tip, 55° C water drop). **h**. Comparison of peak Z-scores (top) and area under the curve (AUC; bottom) achieved with each stimulus type during the first 5 seconds after stimulus delivery. **i**. The dynamic thermal assay presents mice with a changing thermal surface (i.e., 30° C to 50° C, at 10°C/min) to assess changes in nociception and nociceptive activity of vlPAG MOR+ cells. Averaged traces in the drug-naïve and morphine-treated (10 mg/kg, i.p.) conditions are shown. **j**. A representative mouse displayed a range of affective-motivational self-attentive and escape-related behaviors that corresponded with unique changes in the bulk fluorescent signal during the noxious phase of the thermal ramp (bottom). Example behaviors that were mapped onto the calcium trace are denoted with arrowheads and plotted (top). **k, l**. Comparison of group averaged mean Z-score during the 1 second immediately following behavior onset for escape-related (**k**) and self-attentive behaviors (**l**). **m**. Correlation and separate comparisons of innocuous GCaMP6f signal AUC and nocifensive behaviors. **n**. Correlation and separate comparisons of noxious GCaMP6f signal AUC and nocifensive behaviors.

*Oprm1*-expressing neurons in the PAG are a mixed population, largely expressing glutamatergic or GABAergic markers^27,28^, with the RVM-projecting population being almost entirely glutamatergic^28^. However, it remains unclear what proportions of these subpopulations of *Oprm1*-positive glutamatergic and GABAergic cells may be responsive to noxious stimuli. Therefore, we examined co-expression of *Vglut2-* and *Vgat*-expressing *Oprm1-*positive cells with noxious 55 °C water stimulation-induced *Fos*. Like previous studies examining *Oprm1* overlap with glutamatergic and GABAergic mRNA markers, we found that across the entire PAG there was greater overlap of *Vglut2* with *Oprm1*+ cells when compared with those cells co-expressing *Vgat* (**Fig. 2d, left** and **middle**). There were also significant differences within and between individual columns when comparing percent overlap of *Vglut2* or *Vgat* with *Oprm1*, with lPAG possessing comparatively fewer *Vgat*+ neurons co-expressing *Oprm1* than the other columns (**Fig. 2d, middle**). Finally, we assessed percent overlap of co-expressed *Oprm1* and noxious stimulation-induced *Fos* in either *Vglut2*+ or *Vgat*+ cells in PAG. For the entire PAG, we found significantly greater overlap of co-expressed *Oprm1* and *Fos* with *Vglut2* than *Vgat* (**Fig. 2d, right**). At the level of individual columns, vl and lPAG contained substantially greater proportions of *Vglut2*+ cells co-expressing *Oprm1* and noxious *Fos* mRNA than *Vgat*/*Oprm1*/*Fos* overlapping cells when compared to the other columns (**Fig. 2d, right**). With these results, we show that there is substantially greater *Oprm1* and *Vglut2* overlap across the entire PAG. Moreover, when *Oprm1* cells co-express noxious stimulation-induced *Fos*, those cells are more likely to be glutamatergic within the posterior lPAG and vlPAG.

While static measures of neural activation are useful for identifying subregions and cell types engaged by various stimuli, these methods lack the real-time activity dynamics afforded by *in vivo* imaging modalities. Therefore, we sought to provide insight into the functional recruitment of MOR-expressing PAG neurons with peripheral presentation of increasingly noxious stimuli. We focused our investigation on the posterior vlPAG (−4.60 to −4.72 mm relative to Bregma) due to its longstanding role in opioid-mediated analgesia^19,57^ and the high level of noxious stimulus-induced activation we observed across our molecular phenotyping experiments. To do this, we used a *mMOR* promoter AAV construct^30^ expressing the genetically-encoded calcium indicator GCaMP6f to achieve population-level fiber photometry recordings in the vlPAG (**Fig. 2e, left** and **top right**). As an initial assay of nociceptive function of vlPAG MOR+ cells, we delivered three batteries of distinct sensory stimuli including light touch, noxious mechanical stimulation, and noxious thermal heat (*i*.*e*., 0.16 g von Frey filament, 25G needle pin prick, and 55 °C water). Each trial of a given stimulus consisted of five presentations of the stimulus to the left hindpaw with an inter-stimulus interval of two minutes and an inter-trial interval of five minutes between different stimulus types (**Fig. 2e, bottom right**). Representative trial recordings from one mouse show the relative magnitude of MOR+ population calcium responses to each stimulus; population recruitment scaled with stimulus intensity (**Fig. 2f**). This effect was consistent across all mice tested (n=12; 5 males, 7 females) when averaged across the five presentations (**Fig. 2g**). Quantitative comparison of the peak Z-score across stimulus types revealed significantly greater activation of MOR+ vlPAG cells with hot water than both pin prick and 0.16 g von Frey filament (**Fig. 2h, top**). Similarly, when assessing total fluorescence across the first five seconds after stimulus application, 55 °C water produced the largest bulk fluorescence (*i*.*e*., area under the curve, AUC) increase compared to pin prick and 0.16 g von Frey filament (**Fig. 2h, bottom**). Analysis of stimulus-evoked responses across the five presentations revealed no significant changes in magnitude of response over a given trial regardless of stimulus type and we found no difference when comparing the bulk fluorescent change induced by each stimulus type between sexes (**Extended Data Fig. 8a-g**). These data reveal that vlPAG MOR+ cells, at least transiently, report noxious experience.

Nociceptive activation of vlPAG MOR-expressing cells is likely nuanced and related to individual behaviors and the circuits that subserve those behaviors. To examine nociceptive vlPAG MOR+ population engagement beyond acute all-or-none measures, we employed an inescapable dynamic thermal plate assay in which the temperature of the plate ramped from neutral 30° C to noxious 50° C at a rate of approximately 10° C per minute. With this task, we observed behavioral and neural population state changes across innocuous and noxious temperature ranges (i.e. < or >42° C). Fiber photometry recordings showed that the MOR+ vlPAG population in mice naïve to the test and any drug treatment (n=12; 5 males, 7 females) substantially increased in bulk calcium-related fluorescence as the temperature of the plate changes from innocuous to noxious (**Fig. 2i, black trace**). In contrast, when the same mice were pretreated with an analgesic dose of morphine (i.e. 10 mg/kg, i.p.), there was a marked blunting of the noxious-related fluorescent signal (**Fig. 2i, blue trace**).

Analysis with the Behavioral Observation Research Interactive Software (BORIS)^58^ was performed to assess morphine’s analgesic efficacy in the dynamic thermal plate assay and to align fiber photometry signals to individual affective-motivational nocifensive behaviors^1^ performed by the mice. These nocifensive behaviors included those within a self-directed attentive domain: hindpaw guarding, hindpaw licking, and forepaw licking; and an externally-directed escape domain: rearing and jumping. The fiber photometry recording from one naïve male mouse during the noxious phase of the dynamic thermal plate assay is shown with representative peri-event time histograms (PETHs) displayed for each nocifensive behavior type analyzed (**Fig. 2j, bottom**). With self-directed attentive behaviors, we generally observed a decrease in GCaMP signal that followed onset of the behavior, whereas onset of escape behaviors produced an increase in GCaMP signal (**Fig. 2j, top**). Group-averaged PETHs revealed the time course of MOR+ population activity relative to onset of each behavior. During the first second after behavior initiation, we observed a significantly greater mean Z-score for jumping relative to rearing in the naïve condition (**Fig. 2k**); insufficient instances of rearing and jumping were observed to make statistical comparisons between the naïve and morphine-treated conditions for escape behaviors. With respect to self-attentive nocifensive behaviors, we found significant differences between groups for hindpaw guarding, hindpaw licking, and forepaw licking, whereby morphine treatment blunted each behavior-related change in signal (**Fig. 2l**).

We further analyzed the relationship between the fiber photometry signal recorded from the vlPAG MOR+ population and the nocifensive behaviors mice performed under either naïve or morphine-treated conditions. During the innocuous temperature phase of the dynamic thermal plate assay, we observed no significant relationship between the bulk fluorescence signal and nocifensive behaviors performed; however, we did observe a modest, yet significant reduction in nocifensive behaviors following morphine treatment when compared to the naïve condition (**Fig. 2m**). These behaviors primarily consisted of rearing and forepaw licking. Contrastingly, we saw a significant positive correlation between noxious temperature-evoked bulk fluorescence and nocifensive behaviors displayed with significantly higher bulk fluorescence and nocifensive behaviors performed in the drug naïve condition relative to the morphine-treated condition (**Fig. 2n**). Additionally, analysis of the fiber photometry bulk fluorescence and nocifensive behaviors across sexes revealed no significant differences at innocuous or noxious temperatures for both the naïve and morphine-treated conditions (**Extended Data Fig. 8h-l**). With respect to the effect of morphine on basal neural activity, we found that morphine significantly reduced spontaneous calcium-related transients (*i*.*e*., not evoked by nociceptive stimulus) in a sex-independent manner, indicating an overall reduction in excitability of the MOR-expressing vlPAG population caused by morphine administration (**Extended Data Fig. 8m-q**). In total, our data demonstrate that the vlPAG MOR+ cells are a largely glutamatergic population that is dynamically engaged in nociceptive behaviors and sensitive to agonist availability.

#### Pain state-dependent regulation of endogenous opioid release in vlPAG

Opioid analgesics mimic endogenous opioid peptide function in nociceptive neural circuits by engaging MOR signaling, resulting in antinociception and pain relief when administered directly into sites such as the vlPAG^19^. In so doing, exogenous opioid drugs mask the endogenous opioid peptide signaling dynamics that typically modulate MOR+ cells. The vlPAG contains a high density of enkephalinergic fibers^18,59^ that terminate on both GABAergic and glutamatergic neurons^15^. Local vlPAG neurons represent a discrete source of enkephalinergic inputs, yet the forebrain afferents releasing enkephalins in the vlPAG have not been fully characterized. Moreover, direct evidence of functional release of enkephalins in vlPAG from any input is scant^60–62^ and the sources of this innervation and temporal dynamics of enkephalin release under acutely noxious stimulation, protracted or chronic pain states remains unresolved. To begin to address these gaps in knowledge, we first mapped *Penk* expression within the columns of the posterior PAG and its overlap with *Fos* mRNA in home-cage controls (n=3 male mice) and noxious 55 °C water-stimulated mice (n=4 male mice) (**Fig. 3a**). Total number of cells expressing *Penk* mRNA within each column of the PAG differed significantly, with vlPAG possessing the highest number of *Penk*+ cells (**Fig. 3b, left**), with no overall difference in proportions of cells expressing low, moderate, high, or very high levels of *Penk* transcripts (**Extended Data Fig. 9a-c**). Following noxious water exposure, we found significantly more *Fos*-expressing *Penk*+ cells than in home cage controls when analyzing across the entire PAG; similarly, each PAG column within noxious-stimulated mice showed significantly greater co-expression of *Fos* in *Penk*+ cells than in home cage controls (**Fig. 3b, right**). At the protein level (n=6 mice; 3 male, 3 females; **Fig. 3c, left**), the distribution of met-enkephalin positive fibers corresponded with the distribution of *Penk*+ cells in PAG. We found that vlPAG possessed the greatest enkephalinergic innervation among all columns (**Fig. 3c, right**), beginning at approximately −4.60 mm relative to Bregma (**Extended Data Fig. 9d**). These results suggest a clear and substantial enkephalinergic innervation of vlPAG that may arise from within vlPAG itself, but likely includes multiple sources such as the central amygdala^60^ and parabrachial nucleus^28^, among others (**Extended Data Fig. 9e-f**).

**Figure 3.**
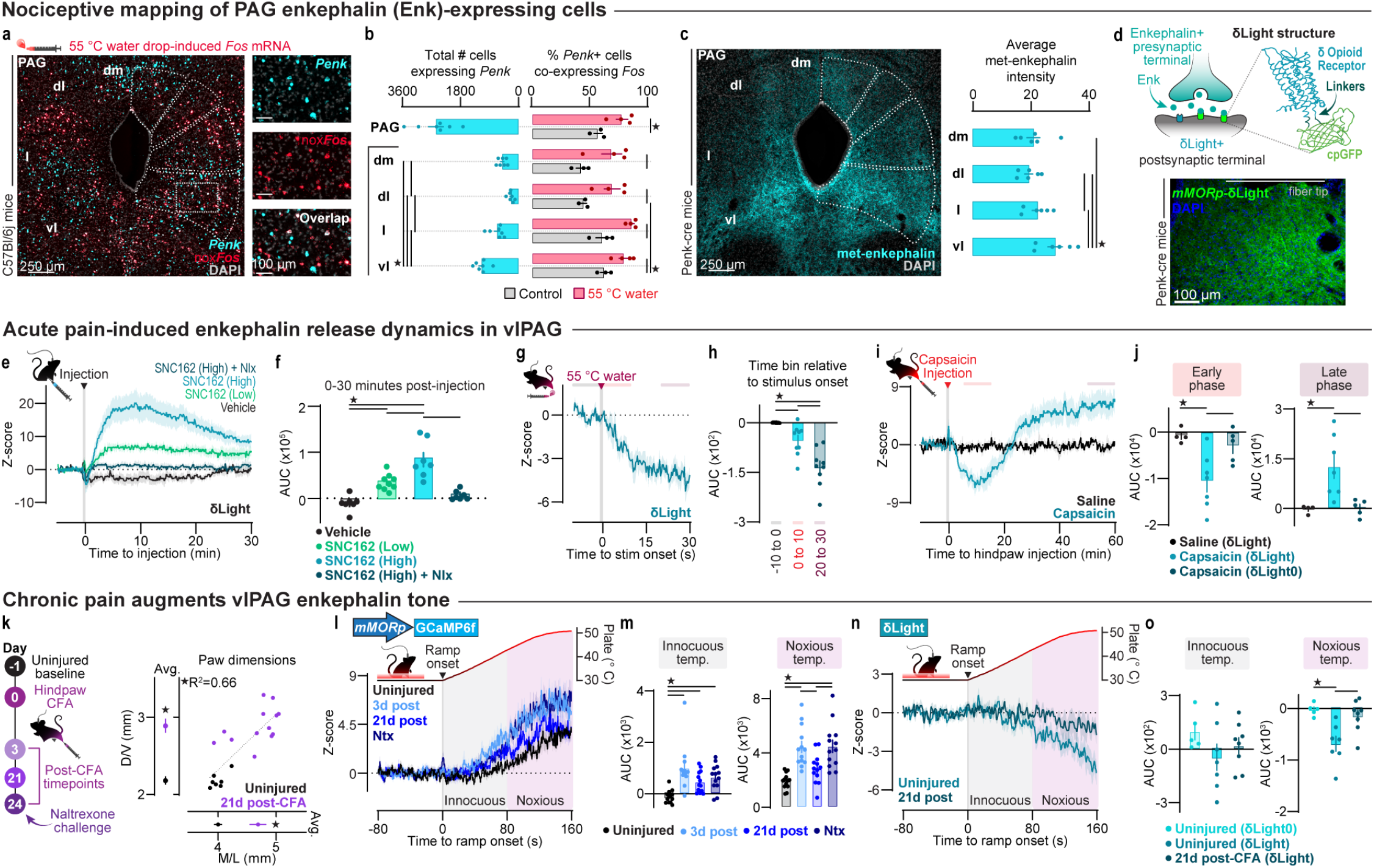
Rapid shifts in vlPAG enkephalin release define transitions from acute to protracted pain states. **a**. Representative 20x fluorescence *in situ* hybridization images demonstrating mRNA transcripts for *Penk* (teal) and *Fos* (red) in vlPAG at approximately A/P −4.72 relative to Bregma. **b**. Quantification of total number of cells expressing *Penk* (left) and comparison of the percent of *Penk*+ cells co-expressing *Fos* between Control and 55° C water-stimulated mice (right) in the PAG and individual columns. **c**. Met-enkephalin immunoreactivity in PAG (left) and comparison on fluorescence intensity across PAG columns (right). **d**. Didactic for δLight function *in vivo* (top) and representative 20x image displaying *mMORp*-δLight expression (bottom). **e**. Averaged traces demonstrating agonist-induced changes in δLight fluorescence. **f**. Comparison of bulk fluorescence changes in δLight arising from vehicle, δ opioid receptor agonist SNC 162 (2.5 or 5.0 mg/kg, i.p.), or co-administration of SNC 162 (5.0 mg/kg, i.p.) with the antagonist naloxone (4.0 mg/kg, i.p.). **g**. Averaged δLight response to noxious 55° C water application to the left hindpaw. **h**. Comparison of δLight signal change across timepoints relative to onset of the 55° C water application. **i**. Averaged δLight responses to either saline or capsaicin (10 µg) to the left hindpaw. **j**. Comparison of δLight bulk fluorescence during the early (left; 5-15 min) and late (right; 50-60 min) timepoints relative to saline or capsaicin administration in mice expressing either δLight or the control sensor δLight0. **k**. Timeline for fiber photometry recordings before and after induction of bilateral hindpaw inflammatory pain with complete Freund’s adjuvant (CFA; left). Correlation and separate comparisons of CFA-inflamed and uninjured average paw depth (D/V) and width (M/L) at the time of perfusion approximately 28 days after hindpaw injections (right). **l**. Averaged *mMORp*-GCaMP6f recordings captured at multiple timepoints after CFA administration during the dynamic thermal plate assay. **m**. Comparison of bulk calcium-related fluorescence at innocuous (left) and noxious temperatures (right) of the thermal ramp. **n**. Averaged δLight traces recorded in the dynamic thermal plate assay at pre- and 3 weeks post-CFA timepoints. **o**. Comparison of bulk δLight fluorescence at innocuous (left) and noxious temperatures (right) of the thermal ramp.

Existing attempts to measure enkephalin peptide levels directly in PAG have relied on the slow-timescale modality of microdialysis that collects samples on the order of tens of minutes^60–62^. These types of measurements occlude sub-minute and sub-second alterations in enkephalin release dynamics that likely occur. Therefore, we employed a newly developed fluorescence-based enkephalin biosensor called δLight^63^. This sensor can be targeted to cell types of interest, such as MOR+ cells with *mMORp* AAVs, and enables post-synaptic detection of enkephalin release and synaptic enkephalin tone in real-time, as binding of enkephalin peptide to the sensor combined with blue light excitation elicits green fluorescence that can be detected with standard fiber photometry equipment (**Fig. 3d**; **Extended Data 10a-c**). We first sought to demonstrate the ability of δLight expressed in vlPAG to detect and report agonist binding *in vivo* in awake, behaving mice using the exogenous δ opioid receptor agonist SNC 162. Across treatment conditions (n=16 mice, 11 males and 5 females), δLight produced fluorescence increases that scaled with dose of SNC 162 administered (**Fig. 3e**). SNC 162 at either the low (2.5 mg/kg, i.p.) or high dose (5.0 mg/kg, i.p.) produced significant signal increases as measured by area under the curve (**Fig. 3f**); the high dose elicited a significantly larger fluorescent response than the low dose. The effect of the high dose of SNC 162 was suppressed by co-administration of the antagonist naloxone (4.0 mg/kg, i.p.). An agonist binding-deficient version of the sensor, δLight0 (n-5, 3 males, 2 females), was used as a control for δLight and produced no substantial change in fluorescence in response to SNC 162 (**Extended Data Fig. 10d-f**). Additionally, we tested the ability of multiple δLight-expressing AAV constructs to fluoresce in response to the δ opioid receptor agonist SNC 162. When we compared *hSyn* and *mMOR* promoter AAVs, we found no difference in bulk fluorescence increase resulting from SNC 162 (5.0 mg/kg, i.p.; **Extended Data Fig. 10g-i**) or decrease resulting from naloxone (4.0 mg/kg, i.p.) given 45 min after SNC 162 (**Extended Data Fig. 10j-k**); therefore, we combined data for mice injected with these viruses across experiments unless explicitly stated otherwise. Overall, these data demonstrate the functionality and dose-dependent responsiveness of δLight to agonist *in vivo*.

We then sought to observe changes in enkephalin tone or release across pain states, from acute nociception to protracted and chronic pain. How enkephalin release is altered by acute nociceptive stimulus application in vlPAG has not been previously examined or reported. In contrast to the rapid 55 °C water drop-induced rise in MOR-expressing cell activation, we observed noxious hot water-related reduction in enkephalin tone (n=9 male mice; **Fig. 3g**) that persisted for at least 30 seconds after the initial stimulus application occurred (**Fig. 3h**) and remained consistent across all five stimulus applications (**Extended Data Fig. 11a**). This result indicates that acute nociception, on the order of seconds, attenuates enkephalin tone. However, we sought to examine whether pain states that extend over minutes or longer could result in different release dynamics. To investigate protracted pain-induced enkephalin release, we used the intraplantar capsaicin test^64^ in which the transient receptor potential vanilloid 1 (TRPV-1) receptor agonist capsaicin^65^ (10 μg) is injected into the hindpaw to induce a pain state that would persist while recording δLight or δLight0 signal over a longer time scale. Using this approach, we observed a capsaicin-induced (n=7, 4 males, 3 females) biphasic change in enkephalin release dynamics (**Fig. 3i**), with an early phase (i.e. 5-15 min post-injection) characterized by diminished enkephalin tone (**Fig. 3j, left**) and a late phase (i.e. 50-60 min post-injection) with heightened enkephalin release (**Fig. 3j, right**) that was not observed in saline-treated mice (n=4, 2 males, 2 females) or in δLight0-expressing mice (n=5, 3 males, 2 females; **Extended Data Fig. 11b**). These results suggest that there is ongoing nociceptive control of enkephalin release in vlPAG at proximal and later timepoints relative to onset of a nociceptive or pain state.

How endogenous opioids are regulated or released in the PAG during chronic pain is relatively unexplored. One clear example of this, however, occurs with Complete Freund’s Adjuvant (CFA)-induced inflammatory pain^66^. Across the progression of inflammation, mice treated with CFA demonstrate allodynia^67,68^ and thermal hypersensitivity^69,70^ that improves after the first week. By the third week post-injury, MORs in the spinal cord undergo plasticity that renders the receptors constitutively active such that CFA-treated mice display uninjured-like mechanical sensitivity that can be reverted to the injured state with naloxone administration; this phenomenon has been termed latent sensitization^67^. This switch in MOR function involves endogenous opioid upregulation. For these reasons, CFA-induced inflammatory pain is a tractable model for recording changes in vlPAG MOR-expressing cell activity and enkephalin tone at long time scales. Here, we performed fiber photometry recordings with either *mMORp*-GCaMP6f pre-CFA, and at timepoints 3-, and 21-days post-CFA intraplantar administration to both hindpaws, followed by naltrexone challenge on day 24, or δLight prior to CFA administration and 21 days post-CFA (**Fig. 3k, left**). When measured at 3 weeks after CFA treatment, GCaMP6f-expressing mice (n=13, 6 males and 7 females) displayed significantly larger paw thickness and width than untreated, age-matched mice (n=7, 4 males and 3 females). Thus, larger paw size corresponded with CFA treatment (**Fig. 3k, right**), verifying the effectiveness of CFA-induced inflammation and confirming that the inflammation persists despite possible changes in nociception.

To examine changes in vlPAG MOR+ population activity at timepoints before and after CFA inflammatory pain induction, we used the dynamic thermal place assay to compare bulk fluorescence changes across innocuous and noxious temperature ranges of the test. With this procedure we observed marked CFA-related elevation of both innocuous and noxious temperature-induced calcium activity in vlPAG MOR+ cells at 3 days post-CFA compared to the uninjured state (**Fig. 3l**). During the innocuous temperature range, CFA treatment resulted in significantly greater bulk fluorescence at days 3, 21, and 24 day (i.e. Ntx, naltrexone challenge) post-CFA when compared to the uninjured condition (**Fig. 3m, left**). At noxious temperatures, however, we found that only at the 3 days post-CFA and naltrexone challenge timepoints was vlPAG MOR+ population calcium signal significantly elevated compared to the uninjured condition, while the 21 day post-CFA recording mirrored the uninjured condition (**Fig. 3m, right**) In contrast, injection of intraplantar hindpaw saline (n=13, 6 males, 7 females) did not reproduce the effect of CFA. We observed only a modest increase in the activity of the vlPAG MOR+ population at noxious temperatures across days that was unaltered by naltrexone challenge (**Extended Data Fig. 12a-d**).

Our recordings with *mMORp*-GCaMP6f, suggested that by 3 weeks after induction of CFA inflammatory pain, there is an upregulation of endogenous MOR agonist in vlPAG that suppresses the thermal hypersensitivity-related activity of MOR-expressing cells, a signature of latent sensitization. Consistent with this hypothesis, increased enkephalin tone has been observed beginning within several hours after CFA administration in posterior vlPAG and extending for at least two weeks thereafter^66^. Here, we extend those prior findings using δLight. Using the dynamic thermal plate assay, we observed noxious temperature-induced reduction in enkephalin tone in uninjured δLight-expressing mice (n=7, 4 males, 3 females **Fig. 3n, light teal trace**) that was comparable between males and females (**Extended Data Fig. 11c**). This noxious thermal heat-induced reduction in enkephalin tone was not observed at 3 weeks post-CFA (n=8, 4 males, 4 females; **Fig. 3n, dark teal trace**). In the innocuous temperature range, there were no differences in bulk fluorescence between δLight uninjured, δLight 3 weeks post-CFA, and δLight0 uninjured groups (**Fig. 3o, left**). At noxious temperatures, however, there was a significant reduction in δLight signal only in the uninjured state when compared to uninjured δLight0-expressing mice, that was not evident at the 3 week post-CFA timepoint (**Fig. 3o, right**), indicating that the nociceptive suppression of enkephalin tone in vlPAG is attenuated several weeks after induction of an inflammatory pain state. This result could relate to an overall increase in synaptic enkephalin, a reduction in enkephalinase activity, or a combination of these processes. These data demonstrate that enkephalinergic innervation of vlPAG is robust and differentially engaged by pain state, whereby acute pain rapidly attenuates enkephalin tone and persistence of pain recruits and increases enkephalin.

#### Cognitive-state modulation of vlPAG opioid circuitry

The vlPAG not only passively relays peripheral nociceptive inputs or descending signals from cortical and sub-cortical areas, but also integrates these signals to update learning about ongoing pain and the contexts in which it is experienced^71,72^. This suggests that cognitive factors related to pain expectation can be harnessed to influence vlPAG opioidergic circuitry and pain experience. The integrative role of vlPAG in pain and negative affect more generally has been studied in the context of fear conditioning paradigms^73–75^ and placebo analgesia^76–78^. However, it remains unclear how genetically-defined opioidergic populations (*i*.*e*., *Oprm1* and *Penk*), long-implicated in pain expectation, participate in vlPAG-mediated pain modulation. To explore this possibility, we first used a trace fear conditioning paradigm to determine how enkephalin tone or release on MOR-expressing vlPAG neurons, as measured with δLight, is regulated by cues (*i*.*e*., conditioned stimulus, CS; tones) and contexts that signal impending noxious stimulation (i.e. unconditioned stimulus, US; shock). Our trace fear conditioning protocol consisted of two recording days, the first being the acquisition phase during which CS-US pairings occurred over 10 trials separated by a variable interval of 50-70 seconds, with each trial consisting of a train of 25 four-kilohertz tones preceding a 20-second period of quiescence (*i*.*e*., trace), followed by a 2-second 1 milliamp shock (i.e., Day 1; **Fig. 4a**). The second recording day captured extinction of the CS-US association, during which each of the 10 trials occurred similarly to the acquisition day without presentation of the shock and by replacing the contextual cues of the conditioning chamber (i.e., Day 2; **Fig. 4a**). On Day 1, acquisition, we observed a consistent pre-shock rise in enkephalin release, that peaked during the trace period, followed by a pronounced and rapid reduction in enkephalin tone that began during the shock (**Fig. 4b, left**). This pattern of enkephalin release dynamics can be seen across most mice tested (n=14, 7 males, 7 females) (**Fig. 4b, right**). Interestingly, the pre-shock rise in enkephalin signal that we detected appeared in both early (1-5) and late (6-10) acquisition trials (**Extended Data Fig. 13a-e**), suggesting that the CS itself, may have an intrinsically valuable or motivational property that the vlPAG encodes through enkephalin release^79^. In contrast, on Day 2, extinction, we observed little to no modulation of enkephalin signal across the tone, trace, and shock periods of all trials (**Fig. 4c, left**), that was largely consistent across all mice tested (**Fig. 4c, right**) and did not substantially differ between early and late trials (**Extended Data Fig. 13a-e**). When comparing the acquisition and extinction days, we found significant differences between days during the early and late tone presentations (*i*.*e*., 1-5 and 21-25), trace, and post-shock periods that were recorded (**Fig. 4f**). When comparing across sexes, we did not detect substantial differences between males in females during either the acquisition or extinction day, except during the pre-shock window of acquisition (**Extended Data Fig. 13f-i**). These data suggest that there may be pain-expectant enkephalin release in vlPAG that occurs prior to onset of a noxious stimulus that may act to blunt or otherwise attenuate the perceived pain of that stimulus.

**Figure 4.**
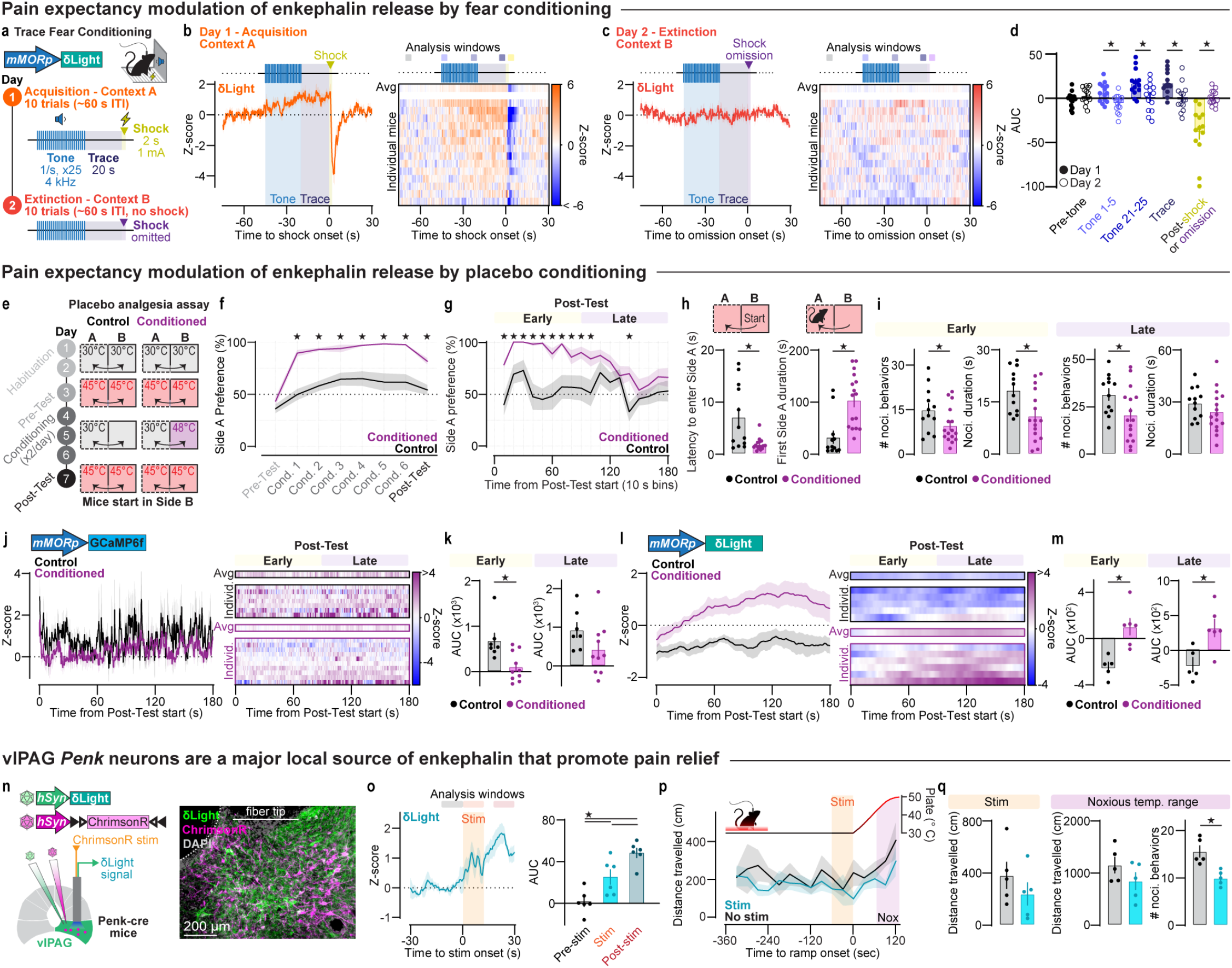
Opposing pain expectancies converge on vlPAG opioidergic neurocircuitry that promotes pain relief. **a**. The trace fear conditioning procedure involved two consecutive recording days. On the first, acquisition day, foot shock was preceded by a series of tones and a 20-second quiescent period. On the second, extinction day, a distinct context was used in which the tone and trace periods were not followed by shock. **b**. Group-averaged *mMORp*-δLight signal (left) and individual animal heatmaps (right) during the acquisition day of the trace fear conditioning protocol. **c**. Group-averaged *mMORp*-δLight signal (left) and individual animal heatmaps (right) during the extinction day of the trace fear conditioning protocol. **d**. Comparison of pre-tone, tone (i.e., tones 1-5 and 21-25), pre-shock, and post-shock 5-second windows between acquisition and extinction days. **e**. Timeline and experimental didactic for the placebo analgesia conditioning (PAC) assay. **f**. Comparison of preference for Side A across each session of the PAC assay between Control and Conditioned groups. **g**. Comparison of preference for Side A within the Post-Test of the PAC assay between Control and Conditioned groups. **h**. Comparisons of latency to enter Side A (left) and total duration of first visit to Side A (right) between Control and Conditioned groups. **i**. Nocifensive behavior counts and durations compared between Control and Conditioned groups during the early phase of the Post-Test (0-90 seconds; left column) and late phase of the Post-Test (90-180 seconds; right column). **j**. Averaged *mMORp*-GCaMP6f traces (left) and individual animal heatmaps (right) from Control and Conditioned groups during the Post-Test of the PAC assay. **k**. Comparisons of bulk GCaMP6f fluorescence between Control and Conditioned groups during the early Post-Test (left) and late Post-Test (right). **l**. Averaged *mMORp*-δLight traces (left) and individual animal heatmaps (right) from Control and Conditioned groups during the Post-Test of the PAC assay. **m**. Comparisons of bulk δLight fluorescence between Control and Conditioned groups during the early Post-Test (left) and late Post-Test (right). **n**. Experimental didactic for recording optically-evoked enkephalin release in vlPAG with δLight (left) and representative 20x image of δLight and Cre-dependent expression of the excitatory opsin ChrimsonR in vlPAG (right). **o**. Group-averaged δLight signal before, during, and after onset of 10 seconds of LED stimulation to activate ChrimsonR (left) and comparison of evoked enkephalin release at each timepoint (right). **p**. The dynamic thermal plate assay was used to assess the effect of unilateral optogenetic stimulation (10 Hz) of right vlPAG Penk cells on overall locomotor activity and nocifensive behaviors. Group-averaged distance travelled is plotted to compare movement across timepoints. **q**. Comparison of distance travelled during the stim period (left) between unstimulated and optogenetically-stimulated groups. Following optogenetic stimulation, the noxious temperature range of the thermal ramp was analyzed for comparison of distance travelled (left) and total nocifensive behaviors (right).

Pain expectancies arise both negative and positive learned factors^80^. In contrast to the cue-primed pain expectation that occurs in fear conditioning or nocebo paradigms, placebo analgesia is pain relief derived from expectations that an intervention or context has therapeutic value, despite being inert or neutral^81^. Pain relief seeking and expectation of an analgesic outcome, therefore, are forms of negative reinforcement that direct an animal toward removal of a source of noxious stimulation to achieve harm reduction and experience pain relief. Moreover, this pain relief expectation involves release of endogenous opioids in the brain, in areas such as the ACC and PAG^82^. Despite these important findings, we still lack basic information at the cellular and neural population levels of the PAG regarding how pain relief expectancies shape or modulate opioid circuit function. For this reason, we employed a model of placebo analgesia that provides a tractable, reproducible experimental platform for interrogating PAG opioid neurocircuitry while mice are actively engaged in a nociceptive task. Our model is based on a recently published report by Chen *et al*.^*83*^. In brief, mice learn that two contiguous hotplate chambers can be either noxious or neutral and mice freely choose between either context (**Extended Data Fig. 14**). During the Conditioning phase, Conditioned mice (n=16, 5 males, 11 females) are those that learn the contingency that pain will be experienced in one context (Side B) and pain relief can be achieved by escaping to and occupying the other, physically connected context (Side A), while Control mice (n=12, 4 males, 8 females) do not learn this contingency. When testing for a placebo analgesia-like effect, the side where pain relief is expected is made noxious (**Fig. 4e**).

In the Post-Test, we measured the efficacy of the placebo analgesia conditioning paradigm across multiple domains, including place preference, instrumental escape, and affective-motivational pain behaviors (i.e., hindpaw guarding, hindpaw licking, rearing, jumping, side escapes or center crosses, digging, and postural extension) to determine if conditioning caused a placebo analgesia-like phenotype. Mice in both the Control and Conditioned groups displayed comparable preference for Side A during Pre-Testing; however, Conditioned mice rapidly learned to escape from Side B during the first Conditioning session, showing significantly greater preference for Side A across all subsequent sessions including the Post-Test when compared to Control mice (**Fig. 4f**). The effect of conditioning that separated the Control and Conditioned groups was maintained during the Post-Test, in which Conditioned mice exhibited heightened preference for Side A, at least during the first 90 seconds of the test, when compared to Controls (**Fig. 4g**). Between sexes, we observed minimal differences in Side A preference among Control mice (n=7, 4 males, 3 females) across all sessions (**Extended Data Fig. 15a-b**); however, among Conditioned mice (n=10, 5 males, 5 females), we found that, during the Post-Test, female mice displayed higher and more sustained preference for Side A than male mice (**Extended Data Fig. 15c-d**). The effect of conditioning was evident from the outset of the Post-Test. Conditioned mice immediately escaped from Side B to occupy Side A, while Controls required significantly more time to transition from Side B to Side A (**Fig. 4h, left**). Conditioned mice also spent significantly more time on Side A during the first visit after escaping from Side B compared to Controls (**Fig. 4h, right**). These initial escape and preference metrics did not differ between males and females in the Control group (**Extended Data Fig. 15e**); however, male mice in the Conditioned group occupied Side A for significantly less time than females during the initial visit (**Extended Data Fig. 15f**). Congruent with enhanced preference for Side A, Conditioned mice engaged in less nocifensive behaviors than Controls during the Early phase of the Post-Test (**Fig. 4i, left**). This placebo analgesia-like effect waned during the Late phase of the Post-Test (**Fig. 4i, right**). Male and females in the Control group exhibited similar amounts of nocifensive behaviors across the Post-Test, while Conditioned males performed more nocifensive behaviors than females during the Late phase of the Post-Test (**Extended Data Fig. 15g-h**). These data collectively demonstrate that placebo analgesia-like behavior was achieved presenting the opportunity to measure endogenous opioid dynamics in Conditioned and Control mice using our *mMORp*-driven biosensors, GCaMP6f and δLight.

Consistent with the observation that placebo analgesia protocols can increase MOR binding in the PAG^77,82,84^, we observed substantial modulation of MOR-expressing neurons and release of enkephalin onto those neurons when testing for placebo analgesia-like behavior. We used fiber photometry to record the calcium-related activity of MOR-expressing vlPAG neurons during the Post-Test of the placebo analgesia assay. Across Conditioned mice, MOR neurons showed robustly blunted activity during the Early phase of the Post-Test that gradually increased as the test progressed (**Fig. 4j**). Area under the curve analysis revealed significantly less bulk calcium signal in vlPAG MOR neurons of Conditioned mice only during Early Post-Test when compared to Controls (**Fig. 4k**). Male and female mice showed comparable modulation of MOR-expressing vlPAG neurons during Early and Late Post-Test in both the Control and Conditioned groups (**Extended Data Fig. 15i-l**); however, male Conditioned mice were more likely to exhibit elevated Late Post-Test bulk calcium signal compared to female Conditioned mice (p=0.084). Based on our combined behavioral and *mMORp*-GCaMP6f imaging data, we reasoned that our conditioning protocol produced stronger placebo analgesia-like phenotypes in female mice and therefore a stronger recruitment of endogenous opioid signaling in vlPAG than that achieved in male mice. To further explore this effect, in Control (n=5 mice) and Conditioned (n=6 mice) females, we measured enkephalin release dynamics during the placebo analgesia conditioning assay Post-Test with δLight expressed in vlPAG. We observed sustained, increased enkephalin release across most Conditioned mice and a sustained decrease in enkephalin tone in Control mice (**Fig. 4l**). Comparison of δLight fluorescence during both Early and Late phase Post-Test revealed significantly greater bulk fluorescence in the Conditioned group when compared to Controls (**Fig. 4m**). Collectively, these data indicate that placebo analgesia conditioning produces a robust pain relief phenotype that covaries with endogenous opioid signaling in the vlPAG, resulting in diminished nociceptive MOR neuron activation and increased enkephalin release.

Expectations related to pain may alter enkephalin release in vlPAG through modulation of activity of the enkephalinergic inputs to this midbrain structure. These inputs arise both from efferent projections into vlPAG from areas like the CeM and PBN, as well as locally-synapsing PAG *Penk* neurons (**Extended Data Fig. 9e-f**). Because *Penk* and met-enkephalin protein are so highly expressed in vlPAG, we hypothesized that the source of pain-relieving enkephalin release in vlPAG is from within vlPAG itself. To investigate this possibility, we performed optogenetic stimulation of *Penk*+ neurons in vlPAG with the excitatory opsin ChrimsonR during fiber photometry recordings of enkephalin release with δLight (**Fig. 4n**). Using a 10-second, 10-Hz optogenetic stimulation protocol, we observed light-evoked enkephalin release during and after cessation of light pulses (**Fig. 4o, left**). Compared to the pre-stimulation window, stimulation resulted in significantly increased enkephalin release that was sustained and elevated for at least 20 seconds after the stimulation occurred (**Fig. 4o, right**). Stimulating *Penk* neurons for 15 minutes at 10 Hz^85^ produced a more pronounced increase in enkephalin release that was maintained for at least 30 minutes after stimulation ended (**Extended Data Fig. 16a-c**). This suggests that direct activation of vlPAG *Penk* neurons can elevate enkephalin tone in a manner that may be antinociceptive, far outlasting the immediate effect of stimulating the neural population. In the dynamic thermal plate assay, in addition to nocifensive behavior quantification, we tracked the movement of mice in the hotplate arena to assess potential locomotor effects of vlPAG *Penk* neuron stimulation, as activation of some genetically-defined (*e*.*g*., *Chx10*)^86–88^ or output-defined populations^42,89^ can produce immobility or freezing phenotypes (**Fig. 4p**). One minute of stimulation of vlPAG *Penk* neurons expressing excitatory opsin (n=5 mice, 3 male, 2 female; **Extended Data Fig. 16d**), at the neutral plate temperature of 30 °C, did not produce a significant reduction in locomotion when compared to unstimulated controls (n=5, 3 male, 2 female; **Fig. 4q, left**). However, in several stimulated mice we observed reduced locomotion that co-occurred with a tail rattle phenotype. During the noxious temperature range of the dynamic thermal plate ramp that immediately followed cessation of the optogenetic stimulation, we saw no difference in locomotion between stimulated and non-stimulated mice (**Fig. 4q, middle**). Critically, optogenetic stimulation of vlPAG *Penk* neurons significantly reduced noxious temperature-related nocifensive behavior when compared to non-stimulated mice (**Fig. 4q, right**). Thus, direct activation of vlPAG *Penk*+ cells is antinociceptive, suggesting that analgesia can be achieved via harnessing expectation-induced activation of this endogenous opioidergic neural population.

## DISCUSSION

The midbrain vlPAG is a key brain structure that participates in the experience of noxious events and the perception of pain. Peripheral, nociceptive signals from the spinal cord are received by the vlPAG and transmitted to other opioid-sensitive subcortical structures to orchestrate defensive, autonomic, and recuperative responses. These pro- and antinociceptive roles of vlPAG are strongly influenced by MOR agonists^19^. Despite the clear role of the vlPAG in ascending and descending pain modulation, our understanding of the endogenous opioid circuitry that may mediate this modulation has trailed basic discoveries of vlPAG functions. Moreover, the conditions and timescales under which MOR-expressing ventrolateral PAG neurons are modulated by external noxious stimuli and endogenous opioid peptide release remain largely obscure. Therefore, we sought to elucidate the molecular and functional aspects of vlPAG MOR+ neurons and their endogenous modulatory counterparts, enkephalin neurons, to discern population dynamics that may be harnessed for pain relief.

### Transcriptionally-defined opioidergic efferents of the vlPAG engage multiple pain responsive neural structures

The PAG is a functionally- and molecularly-heterogeneous structure^4,33^. Previous work revealed that, while multiple PAG columns may display FOS immunoreactivity in response to disparate noxious protocols (*e*.*g*., radiant heat or intraperitoneal acetic acid), the ventrolateral column is consistently activated^7^. Using the genetic capture method TRAP2^29^, we confirmed and extended those previous results, demonstrating clearly that vlPAG, in addition to lPAG, is strongly activated by noxious hot water exposure beyond the existing background activity level observed in control animals. Moreover, these nociceptive vlPAG and lPAG neurons express MOR and project to other affective-motivational brain structures, namely the medial nucleus of the CeM, MDTh, VTA, and RVM. Thus, nociceptive vlPAG neurons are integrated with brain-wide centers that respond to and integrate noxious information^12^. While these projections have been identified in the context of pain, the molecular profile of the projection populations has remained relatively unexplored.

Combining a retrograde-AAV barcoding strategy (i.e., Projection-TAGs^47^) with single nucleus RNA sequencing, here we provide new columnar, functional, and projection population transcriptomic analysis of the PAG. Within our tissue punches, we obtained nuclei both within and adjacent to the PAG columns, including the dorsal raphe nuclei, LDTg, and oculomotor nucleus, in addition to the cerebral aqueduct. To distinguish these subregions, we established *a priori* genetic markers that localize to either individual or multiple PAG columns and the DRN. This parcellation revealed overlap of Projection-TAGs labeling CeM, MDTh, VTA and RVM projectors with the vlPAG column class, congruent with our output tracing. These projection cell types were further differentiated according to IEG activity, with CeM, MDTh, and RVM projectors demonstrating the highest levels of noxious 55 °C water activation and overlap with known targets of active pain and analgesic drug research. Among the Projection-TAG-labeled nuclei, CeM, MDTh, and RVM projectors were exclusively glutamatergic; however, only MDTh projectors exhibited substantial *Oprm1* mRNA expression, despite all four projection populations possessing the promoter for *Oprm1*. Importantly, a lack of *Oprm1* mRNA, however, does not exclude the existence of the *Oprm1* gene as indicated by our nociceptive MOR neuron output tracing. Thus, from our nociceptive, projection, and transcriptional mapping of PAG, we conclude that posterior vlPAG neurons projecting to pain-relevant fore- and hindbrain regions encode and likely transmit acutely noxious stimulus information from the periphery in a manner that is modifiable by MOR agonists.

### Dynamic, pain state-dependent interplay of nociceptive MOR function and enkephalin release in vlPAG

The primacy of vlPAG function in nociception and its ability to produce pain relief through endogenous opioid signaling is well-known^46^. Electrical stimulation of the vlPAG elicits release of the endogenous MOR agonist met-enkephalin^14^ and results in pain relief in both animals and humans^90–93^ that is, at least partially, blocked by MOR antagonists^57,94^. Correspondingly, infusion of MOR agonists into the vlPAG also produces robust antinociception^19,22^, suggesting that endogenous opioid inhibition of vlPAG MOR-expressing neurons may facilitate pain relief by suppressing nociceptive activation of the MOR+ population^28^. Our data support this hypothesis. Here, we show that acute nociception corresponds with a coordinated population response in vlPAG MOR neurons, such that increasingly noxious stimulation elicits larger bulk calcium-related events. Nociceptive calcium-related events in vlPAG MOR neurons corresponded with nocifensive behaviors in the dynamic thermal plate assay. Critically, this nocifensive neural activity was suppressed by peripheral morphine administration and raising the possibility that changes in endogenous opioid release may promote antinociceptive or pain-relieving inhibition of MOR+ cell types in vlPAG.

Like MORs, we found that *Penk* and met-enkephalin protein are most highly expressed in vlPAG. The high density of enkephalinergic fibers in vlPAG^17,18,59,95^ terminate on both GABAergic and glutamatergic neurons^15^, the latter of which are the predominant MOR-expressing population from our data presented here and by others^27^. Evidence that the vlPAG is a recipient of enkephalin peptide presents the question of the source of this release. Enkephalin peptides are likely released from any of several candidate regions that we identified with our retrograde AAV tracing, including but not limited to the central amygdala, parabrachial nucleus, and vlPAG itself, which also sends collaterals to forebrain areas such as the central amygdala^96^ and nucleus accumbens^97^. To measure real-time enkephalin release dynamics, we used the recently-published biosensor δLight^63^. This imaging tool enabled detection of enkephalin at millisecond resolution in genetically-defined cell populations, which is unfeasible with existing microdialysis or voltammetric methods. δLight recordings revealed, for the first time, that acute nociceptive stimulation reduces enkephalin tone in vlPAG. Blunted enkephalin release resulting from noxious water stimulation to the hindpaw occurred in opposition to the immediate increase in MOR cell activation. However, the attenuating effect of acute nociception on enkephalin release was transient during protracted pain elicited by hindpaw capsaicin. We observed a reversal of nociceptive suppression of enkephalin release, such that prolonged pain boosted enkephalin release above baseline. This suggests that enkephalin release is regulated as a function of pain state.

We further explored the hypothesis that MOR neurons and enkephalin release in vlPAG operate in opposition in the context of the long-lasting pain state induced by inflammation. Complete Freund’s Adjuvant (CFA) administered to the hindpaw provokes allodynia^67,68^ and thermal hypersensitivity^69,70^, that improves within several weeks after initial injury, despite the persistence of inflammation in the affected limb. During this inflammatory pain state, endogenous opioid signaling is increased, such that administration of the MOR antagonist naloxone unmasks the inflammation-induced pain phenotype^67^. This phenomenon is termed latent sensitization and has received further investigation in the context of post-operative pain^98^. How CFA inflammatory pain impacts vlPAG opioidergic circuitry has not been thoroughly investigated, however. Previous microdialysis studies revealed that enkephalin release begins to rise within hours after induction of inflammation with CFA^61,66^. In contrast, recent evidence indicates that MOR-expressing vlPAG neurons become hyperexcitable in response to acute thermal radiant heat stimulation within three days immediately following inflammation onset^99^. We confirmed and extended this latter result in the dynamic thermal plate assay. By 3 days post-CFA we observed a substantial increase in noxious heat-related MOR neuron activity; however, by 3 weeks post-CFA, vlPAG MOR neurons displayed pre-injury-like activity in response to the noxious temperature range of the test. With administration of naltrexone, we found that inflammatory thermal hypersensitivity was reversed, consistent with a tonic, elevated enkephalin tone in vlPAG following CFA-induced inflammation. Thus, vlPAG MOR neurons undergo nociceptive activity-dependent regulation involving endogenous opioid release that does not occur with non-inflammatory hindpaw saline administration. With δLight, we demonstrated directly that nociceptive enkephalin tone is elevated and resistant to noxious thermal heat-induced suppression - by 3 weeks post-CFA, we no longer observed a reduction in enkephalin tone or release that occurred prior to injury. This change in nociceptive enkephalin dynamics indicates that increased enkephalin tone is either maintained or further increased multiple weeks after onset of inflammation. Furthermore, our results suggest that reductions in vlPAG enkephalin are permissive for the acute experience of pain, while upregulation of the peptide is an essential component of protracted pain and potential long-term mitigation of injury-related pain.

### Pain-related expectancies modulate vlPAG opioidergic neurocircuitry

Pain is an unpleasant, subjective, multidimensional experience that arises from the integration of nociceptive and contextual information with cognitive processes^100,101^. As such, learned associations between environments, cues, individuals, controllability, and even nonconscious conditioned stimuli are made with the experience of pain (i.e., nocebo) or its relief (i.e., placebo analgesia)^102–104^ that produce expectancies capable of influencing the perception of pain or lack thereof. The predictive value of these expectancies depends on conditioned factors (*e*.*g*., prior learning) and the reliability of the cues or contexts that precede pain or its relief^71^. At the neural level, the PAG is a critical locus of pain expectancies^80,105^ and, more generally, aversive prediction error encoding^73,106^, through its connectivity with affective-motivational brain areas (e.g., central amygdala^60,107^) and its facilitation of defensive responses to threats^7^. Endogenous opioid signaling at MORs in PAG modulate both pain expectancy^81^ and aversive learning^75^.

A prime example of prediction error occurs in fear learning models, wherein mice are presented with a conditioned stimulus (*e*.*g*., tone, CS) that is followed either immediately or after a trace period by an aversive, typically noxious unconditioned stimulus (*e*.*g*., foot shock, US). Over repeated pairings, this association can lead to the display of canonical defensive behaviors, such as freezing. Endogenous opioid signaling modulates the strength of this learning. For example, blockade of MORs with peripherally-administered naloxone can promote acquisition of fear learning in rats^79^ and humans^108^,while intra-vlPAG microinfusion of the selective MOR antagonist CTAP was found to prevent associative blocking of fear^109^. This effect is likely not purely pain-driven as acquisition of fear to aversive, non-noxious unconditioned stimuli is also enhanced by MOR antagonist treatment^110^. From these findings, it is hypothesized that endogenous opioid signaling in vlPAG increases in response to CS presentations as fear conditioning progresses in a manner that limits fear learning itself^75^ and possibly the experience of expected pain. However, real-time measurement of opioid peptide release during fear conditioning has not been done. Therefore, in a trace fear conditioning paradigm, we used δLight to detect changes in enkephalin release across ten fear conditioning trials and subsequent extinction of fear learning. Our findings agree with those prior studies, as we found a CS-evoked increase in enkephalin release onto vlPAG MOR neurons that persisted through the trace period during acquisition of fear learning. Onset of the US caused a rapid, robust, but transient decrease in enkephalin tone below baseline, consistent with the acutely nociceptive effect of the shock. The CS-mediated rise in enkephalin was not evident during extinction, suggesting that both contextual and sensory cues must be intact to engage vlPAG opioid peptide release through fear learning.

Prediction error also influences the outcome of analgesic interventions, exemplified in the clinical phenomenon of placebo analgesia. Placebo analgesia is pain relief derived from expectations that an intervention or context has intrinsic therapeutic value and involves release of endogenous opioids in the brain^77,82,111,112^, while some protocols identified non-opioid-related placebo analgesia mechanisms^113^. Research on placebo effects in preclinical animal models has trailed discovery in humans^102^ due to distinct constraints imposed by methods used and that non-verbal species lack language to express expectancies related to pain relief^114^. Most preclinical studies to date have used classical conditioning with opioids to elicit drug-context associations followed by assessments of reflexive antinociception, with mixed results^115–121^. While these models have demonstrated a role for endogenous opioids, or possibly altered MOR function, in the placebo analgesia-like phenotypes measured, the variability in those studies may be related, in part, to repeated opioid exposure, which can induce paradoxical hyperalgesia^122^. Recent efforts have moved away from the opioid conditioning paradigms, in favor of pain or drug-free analgesic contextual learning^123,124^ or volitional protocols, like that used in our studies based on a recently-published model ^83^. In this placebo analgesia conditioning assay, mice learn that two contiguous hotplate chambers can be either noxious or neutral and freely choose between either context. By measuring differences across multiple domains, including preference, instrumental escape, and affective-motivational pain behavior, we observed clear evidence for a placebo analgesia-like phenotype in the Conditioned group relative to the unconditioned Control group.

Critically, the emergence of the placebo analgesia-like phenotype during the early segment (i.e., 0-90 seconds) of the test session corresponded with both suppression of nociceptive vlPAG MOR neural activity and a rise in released enkephalin. These effects are consistent with reduced availability of MOR receptors seen in human PET imaging scans during the experience of placebo analgesia in humans^82^. Interestingly, we observed putative within-session extinction of the placebo analgesia-like phenotype corresponding with a rise in vlPAG MOR activity, indicative of positive prediction error signaling in vlPAG (*i*.*e*., actual pain experience exceeding expectation). Within these results we found evidence for sex differences in strength of placebo analgesia conditioning. In both preference for the conditioned chamber and nocifensive behavior, male mice exhibited a more rapid transition to extinction-like behavior compared to females. This contrasts with human studies, which have not reached a consensus on the impact of sex on placebo analgesia. Some reports indicate that this effect is more robust in males via differences in stress^125^, for example, while others indicate that placebo analgesia is likely achieved and optimized in different ways between males and females^126^. Taken in conjunction with our fear conditioning results, which demonstrated a sex difference in trace period enkephalin release during acquisition, it is possible that female mice attend to and capitalize on cues predicting an expected outcome more efficiently than male mice in the assays we used. Consequently, during the late Post-Test extinction period, enkephalin remains elevated in the vlPAG of female mice, suggesting that the duration of the placebo analgesia-like effect and concomitant rise in enkephalin may be extended or shortened according to the strength of learning that occurred during conditioning.

### Harnessing enkephalin release for on-demand pain relief

The source of pain- or expectancy-related endogenous opioid release in vlPAG remains unresolved. With our molecular phenotyping data characterizing *Penk* expression and *Penk*+ inputs to vlPAG, we reasoned that local *Penk*-expressing cells are the most abundant source of modulatory enkephalin within vlPAG. Optogenetic activation of the vlPAG *Penk* population produced an elevation in enkephalin release onto local vlPAG MOR neurons expressing δLight that outlasted the duration of light stimulation and scaled with stimulation duration. This prolonged increase in enkephalin was sufficient to suppress nocifensive behaviors typically evoked by noxious thermal heat exposure. Crucially, these data are congruent with the effect of electrical stimulation of PAG on enkephalin release^14^ and the antinociceptive effect of microinfused enkephalin in vlPAG^127^.

Collectively, this work demonstrates a functional dichotomy in the midbrain vlPAG MOR-expressing neurons that positively respond to pain and the neighboring *Penk* neurons that release enkephalin onto MOR neurons in a pain- and cognitive-state-dependent manner. The vlPAG receives direct spinal nociceptive inputs^128^ and consequently this innervation serves to increase MOR cell activity directly while either directly or indirectly blunting enkephalin tone. This suppression of enkephalin signal may serve to permit transmission of nociceptive signals to forebrain affective-motivational centers via projections to medial thalamic nuclei, for example, as nociception is protective. Progression of nociception into protracted or chronic pain states boosts enkephalin release in vlPAG in a manner that may facilitate pain relief and recovery. This tonic pain-related increase in enkephalin can be mimicked through modulation of expectancies surrounding pain or aversive experience and direct optogenetic activation of vlPAG *Penk*+ neurons. Operationally, we hypothesize that increased enkephalin release in vlPAG is antinociceptive while suppressed enkephalin release is pronociceptive.

Human^81,82,105^ and our new rodent evidence now converge to demonstrate that upregulation of endogenous opioid peptide signaling and suppression of vlPAG MOR neuron activity are key mechanisms of analgesia. These insights underscore the dynamic regulation of vlPAG MOR neurons by endogenous opioids in distinct pain states and in response to multiple cognitive expectancies, such as fear and placebo. By specifically enhancing enkephalin signaling within the vlPAG, it may be possible to achieve effective pain relief without the harmful side effects associated with prescribed opioid analgesics. Harnessing the vlPAG *Penk* neural population offers a promising avenue for future pain therapies, with the potential to alleviate dependence on prescribed opioids and deliver precise, safe, and effective pain management solutions.

## ACKNOWLEDGMENTS

This work was supported by: National Institute of Health (NIH) awards NIGMS DP2GM140923 (G.C.), NIDA R00DA043609 (G.C.), NIDA R01DA054374 (G.C.), NINDS R01NS130044 (G.C.), NIDA R01DA056599 (G.C.), NIDA F32DA055458 (B.A.K.), NINDS F99NS135765 (L.E.), NINDS F31NS125927 (J.A.W.), NIDA F32DA053099 (N.M.M),, NIDA R21DA055846 (B.C.R.), NIDA R01DA056829 (V.K.S), NIDDK R01DK128475 (V.K.S), NIDDK R01DK139386 (V.K.S), NINDS U01NS120820 (L.T.), and NINDS U01NS113295 (L.T.). This work was further supported by the Howard Hughes Medical Institute (K.D.), the Whitehall Foundation (G.C.), and the Rita Allen Foundation (G.C.). We thank the Penn University Laboratory Animal Resources (ULAR) veterinarians and husbandry staff for animal care services. We thank Stanford University “Cracking the Neural Code” Program and Gene Vector and Virus Core and the Neurophotonics Centre at Laval University for custom viruses and the Molecular Pathology and Imaging Core at the University of Pennsylvania for sequencing services. Finally, we thank members of the Corder Lab for additional experimental support and discussions.

## DATA AND MATERIALS AVAIALBILITY

All custom viruses and/or plasmids will be available for academic use from the Stanford Gene Vector and Virus Core (https://neuroscience.stanford.edu/research/neuroscience-community-labs/genevector-and-virus-core) and/or by contacting the lead authors. Upon manuscript peer-reviewed publication, all data will be unembargoed at Zenodo and GEO.

## CONTRIBUTIONS

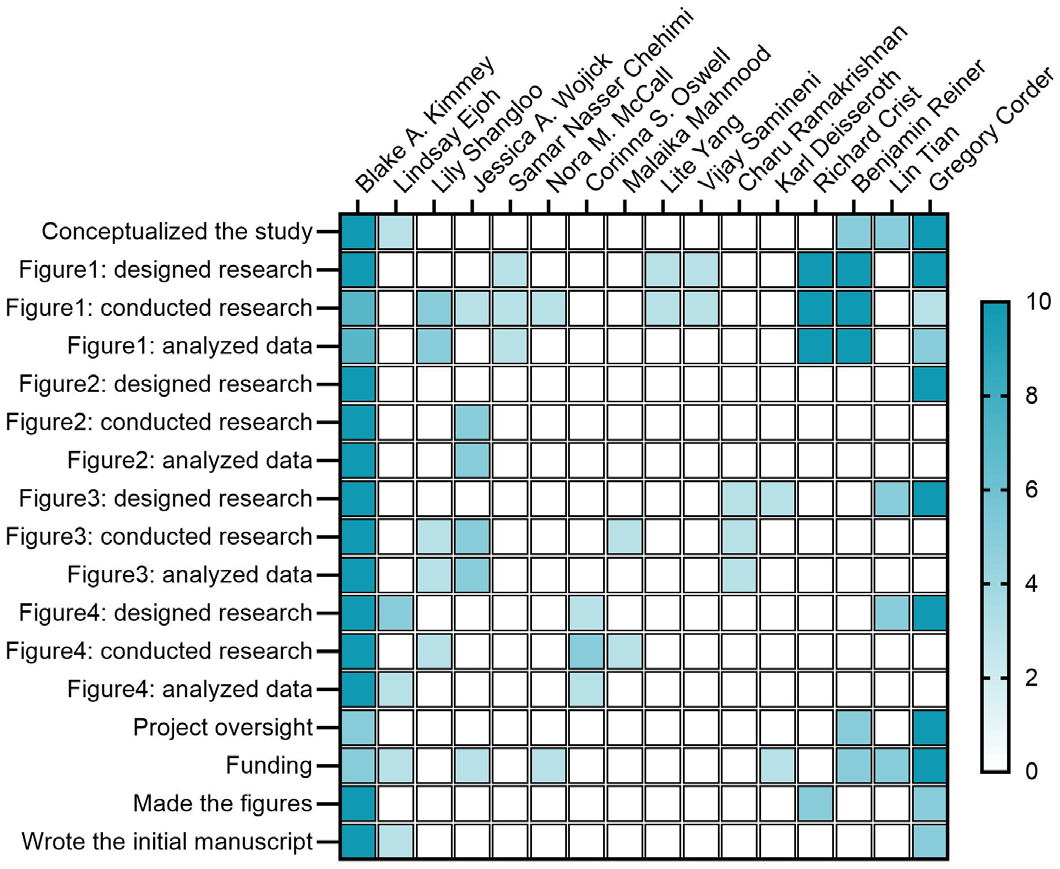

## METHODS

### Subjects

All experimental procedures were approved by the Institutional Animal Care and Use Committee of the University of Pennsylvania and performed in accordance with the US National Institutes of Health (NIH) guidelines. Male and female mice aged 2-6 months were obtained from Jackson Laboratory and housed 2-5 per cage while maintained on a 12-hour reverse light-dark cycle in a temperature and humidity-controlled environment. All behavioral experiments were performed during the dark cycle. Mice had ad-libitum food and water access throughout experiments. For anatomical experiments relying on FOS-mediated neuronal tagging, we used Fos-FOS-2A-iCreERT2 (TRAP2; Fos^tm2.1(icre/ERT2)Luo)Luo/J^, Strain #030323) or TRAP2 mice crossed with Ai9 mice (B6.Cg-Gt(ROSA)26Sor^tm9(CAG-tdTomato)Hze/J^, Strain #007909) reporter mice that express a tdTomato fluorophore in a Cre-dependent manner. In all other experiments, we used C57BL/6J wild type mice (Strain #000664) and Penk-IRES2-Cre mice (Strain #025112). Mice used in fiber photometry and behavioral experiments were randomly assigned to control and test groups following recovery from surgery.

### Viruses

All viral vectors were either purchased from Addgene.org, or custom designed and packaged by the authors as indicated. All AAVs were aliquoted and stored at −80°C until use and then stored at 4°C for a maximum of four days. The following AAVs were used:

- AAV1*-mMORp*-GCaMP6f (Deisseroth Lab, Stanford University; titer: 3.69×10^12^ vg/ml)
- AAV1-*mMORp*-δLight (Deisseroth University; titer: 7.70×10^11^ vg/ml) Lab, Stanford
- AAV1-*mMORp*-FlpO (Deisseroth Lab, Stanford University; titer: 1.28×10^12^ vg/ml)
- AAV8-*hSyn*-Con/Fon-eYFP (Addgene, 55650; titer: 2.40×10^12^ vg/ml)
- AAV1-*hSyn*-δLight (Tian Lab, Max Planck Florida Institute for Neuroscience; titer: 1.8×10^12^ vg/ml)
- AAV1-*hSyn*-DOR3-V154K-PRC-ER2 (δLight; Neurophotonics Core, Laval University; titer: 3.4×10^12^ vg/ml)
- AAV1-*hSyn*-δLight 0 (Neurophotonics Core, Laval University; titer: 6.10×10^12^ vg/ml)
- AAVrg-*CAG-*Projection-TAG1 (Samineni Lab, Washington University in St. Louis; titer: 10^12^ vg/ml)
- AAVrg-*CAG-*Projection-TAG2 (Samineni Lab, Washington University in St. Louis; titer: 10^12^ vg/ml)
- AAVrg-*CAG-*Projection-TAG3 (Samineni Lab, Washington University in St. Louis; titer: 10^12^ vg/ml)
- AAVrg-*CAG-*Projection-TAG5 (Samineni Lab, Washington University in St. Louis; titer: 10^12^ vg/ml)
- AAVrg-*CAG-*Projection-TAG6 (Samineni Lab, Washington University in St. Louis; titer: 10^12^ vg/ml)
- AAV5-*Syn*-FLEX-rc[ChrimsonR-tdTomato] (Addgene, 62723; titer: 2.00×10^12^ vg/ml)
- AAV8-*Ef1a*-DIO-ChRmine-oScarlet (Deisseroth Lab, Stanford University; titer: 2.43×10^12^ vg/ml)
- AAVrg-*EF1a*-Nuc-flox(mCherry)-EGFP (Addgene, 112677; titer: 2.5×10^13^ vg/ml)

The mouse μ-Opioid Receptor promoter (*mMORp*) is a 1.5 Kb segment selected and amplified from mouse genomic DNA using cgcacgcgtgagaacatatggttggacaaaattc and ggcac-cggtggaagggagggagcatgggctgtgag as the 5’ and 3’ end primers respectively. All *mMORp* plasmids were constructed on an AAV back-bone by inserting either the *mMOR* promoter ahead of the gene of interest (e.g., GCaMP6f) using M1uI and AgeI restriction sites. Every plasmid was sequence verified. Next, all AAVs were produced at the Stanford Neuroscience Gene Vector and Virus Core. Genomic titer was determined by quantitative PCR of the WPRE element. All viruses were tested in cultured neurons for fluorescence expression prior to use in vivo.

### Stereotaxic surgery

Adult mice (∼8 weeks of age) were anesthetized with isoflurane gas in oxygen (initial dose = 5%, maintenance dose = 1.5%), and fitted into WPI or Kopf stereotaxic frames for all surgical procedures. 10 μL Nanofil Hamilton syringes (WPI) with 33 G beveled needles were used to intracranially infuse AAVs into specified brain regions and place optic fibers into the vlPAG. The following coordinates were used, based on the Paxinos mouse brain atlas and experimenter practice, to target these regions of interest: vlPAG (from Bregma, AP: −4.60 mm, ML: ± 0.50 mm, DV:−3.00 mm), CeM (from Bregma, AP: −1.00 mm, ML: ± 2.40 mm, DV:−5.00 mm), MDTh (from Bregma, AP: −1.60 mm, ML: ± 0.65 mm, DV:−3.60 mm), VTA (from Bregma, AP: −3.20 mm, ML: ± 0.70 mm, DV:−5.00 mm), and RVM (from Bregma, AP: −5.85 mm, ML: ± 0.25 mm, DV:−6.10 mm). Mice were given a 3–8-week recovery period to allow ample time for viral diffusion and transduction to occur. For fiber photometry and optogenetics studies, following viral injection into the right vlPAG, we placed a fiberoptic implant (400 μm diameter, 0.60 NA, Doric Lenses) approximately 0.2-0.3 mm above the DV coordinate of the injection site for the vlPAG. After setting the fiber optic in position, MetaBond (Parkell) and Jet Set dental acrylic (Lang Dental) were applied to the skull of the mouse to rapidly and firmly fix the fiberoptic in place. Prior to cementing, the exposed skull bone was scored with a scalpel blade and two small screw (∼1.7 mm diameter, 1.6 mm length) were partially threaded into the skull to provide adhesion points for the MetaBond. The MetaBond reagent was applied over the skull surrounding the optic fiber ferrule. Once dried, the MetaBond was then covered with a layer of Jet Set acrylamide dyed with iron oxide to create a reinforced head cap, as well as to cover the exposed skin of the incision site. Mice were then given a minimum of 3 weeks to recover and allow for optimal viral transduction and wound healing prior to beginning *in vivo* recordings. For all surgical procedures in mice, meloxicam (5 mg/kg, Norbrook, 6451602670) was administered subcutaneously at the start of the surgery, and a single 0.25 mL injection of sterile saline was provided upon completion. All mice were monitored for up to three days following surgical procedures to ensure proper recovery and to provide additional daily subcutaneous meloxicam to reduce pain and inflammation.

### Capsaicin acute pain model

To induce a prolonged, but reversible acute pain state, a total mass of 10 µg of capsaicin (Tocris), dissolved in Kolliphor oil and 0.1 M phosphate-buffered saline (PBS) was administered to the plantar surface of the hindpaws via a 20 µl Hamilton syringe fitted with a 33 gauge needle. Injections occurred at least 2 hours after the onset of the dark phase. At the time of injection, mice were gently restrained to isolate the hindpaws. Any excessive bleeding or leakage of capsaicin was noted. Capsaicin was administered to all mice in a cage selected for the acute pain study (i.e., uninjected and injected mice were not co-housed). Similar procedures were followed for saline administration to the plantar surface of the hindpaws. Capsaicin administration occurred after a 10 minute habituation period in an acrylic plastic box (16.51 cm × 16.51 cm) during a continuous fiber photometry recording. Administration of capsaicin was logged in the fiber photometry software Synapse (Tucker-Davis Technologies) with an Arduino Uno trigger button connected to a digital input of the RZ10X photometry processor (TDT).

### Complete Freund’s Adjuvant (CFA) inflammatory pain model

To induce a persistent inflammatory pain state, a total volume of 10 µl CFA (Sigma-Aldrich) was administered to the plantar surface of the hindpaws via a 20 µl Hamilton syringe fitted with a 33 gauge needle. Injections occurred at least 2 hours after the onset of the dark phase. At the time of injection, mice were gently restrained to isolate the hindpaws. Any excessive bleeding or leakage of CFA was noted. CFA was administered to all mice in a cage selected for inflammatory pain studies (i.e., uninjured and injured mice were not co-housed). Mice were subsequently monitored for any health issues arising from CFA administration. After completion of CFA experiments, approximately 3 weeks after induction of CFA inflammation, paw thickness was measured with a digital dial caliper while mice were anesthetized.

### Targeted recombination in active populations (TRAP) protocol

#### Noxious TRAP (noxTRAP)

*Nox*TRAP induction was performed similarly to previously described^129^. We habituated mice to a testing room for two to three consecutive days. During these habituation days, no nociceptive stimuli were delivered (i.e., mice were naïve to pain experience before the TRAP procedure). We placed individual mice within red plastic cylinders (∼9 cm in diameter), with a red lid, on a raised perforated, flat metal platform (61 cm × 26 cm). The experimenter sat in the testing room for the thirty minutes of habituation; this was done to mitigate potential alterations to the animal’s stress and endogenous antinociception levels. To execute the TRAP procedure, mice were placed in their habituated cylinder for 30 min, and then a 55 °C water droplet was applied to the central-lateral plantar pad of the left hindpaw once every 30 s over 10 min. Following the water stimulations, the mice remained in the cylinder for an additional 60 min before injection of 4-hydroxytamoxifen (4-OHT, 40 mg/kg in vehicle; subcutaneous). After the injection, the mice remained in the cylinders for an additional 4 hours to match the temporal profile for Fos protein expression, at which time the mice were returned to the home cage.

#### Home cage TRAP (hcTRAP)

*Hc*TRAP induction was performed without habituation. At least 2 hours into the dark cycle, mice were gently removed from their home cages. Mice were then injected with 4-OHT (40 mg/kg in vehicle; subcutaneous) and returned to their home cages.

### Immunohistochemistry

Animals were anesthetized using FatalPlus (Vortech) and transcardially perfused with 0.1 M phosphate-buffered saline (PBS), followed by 10% neutral buffered formalin solution (NBF, Sigma, HT501128). Brains were quickly removed and post-fixed in 10% NBF for 24 hours at 4 °C, and then cryo-protected in a 30% sucrose solution made in 0.1 M PBS until sinking to the bottom of the storage tube (∼48 h). Brains were then frozen in Tissue Tek O.C.T. compound (Thermo Scientific), coronally sectioned on a cryostat (CM3050S, Leica Biosystems) at 30 μm or 50 μm and the sections were stored in 0.1 M PBS. Floating sections were permeabilized in a solution of 0.1 M PBS containing 0.3% Triton X-100 (PBS-T) for 30 min at room temperature and then blocked in a solution of 0.3% PBS-T and 5% normal donkey serum (NDS) for 2 hours before being incubated with primary antibodies (1°Abs included: chicken anti-GFP [1:1000, Abcam, ab13970], rabbit anti-DsRed [1:1000, Takara Bio, 632496], rabbit anti-met-enkephalin [1:1000, ImmunoStar, 20065]; prepared in a 0.3% PBS-T, 5% NDS solution for ∼16-20 h at room temperature. Following washing three times for 10 min each in PBS-T, secondary antibodies (2°Abs included: Alexa-Fluor 488 donkey anti-chicken [1:500, Jackson Immuno, 703-545-155], Alexa-Fluor 594 donkey anti-rabbit [1:500, Thermo Scientific, A21207] Alexa-Fluor 647 donkey anti-rabbit [1:500, Thermo Scientific, A31573], prepared in a 0.3% PBS-T, 5% NDS solution were applied for ∼2h at room temperature, after which the sections were washed again three times for 5 mins each in 0.1 M PBS, then again three times for 10 min in 0.1 M PBS, and then counterstained in a solution of 0.1 M PBS containing DAPI (1:5,000, Sigma, D9542). Fully stained sections were mounted onto Superfrost Plus microscope slides (Fisher Scientific) and allowed to dry and adhere to the slides before being coated with Fluoromount-G Mounting Medium (Invitrogen, 00-4958-02) and cover slipped.

### Fluorescence *in situ* hybridization

Animals were anesthetized using isoflurane gas in oxygen, and the brains were quickly removed and fresh frozen in O.C.T. using Super Friendly Freeze-It Spray (Thermo Fisher Scientific). Brains were stored at −80° C until cut on a cryostat to produce 16 μm coronal sections of the PAG. Sections were adhered to Superfrost Plus microscope slides, and immediately refrozen before being stored at −80° C. Following the manufacturer’s protocol for fresh frozen tissue for the V2 RNAscope manual assay (Advanced Cell Diagnostics), slides were fixed for 15 min in ice-cold 10% NBF and then dehydrated in a sequence of ethanol serial dilutions (50%, 70%, and 100%). Slides were briefly air-dried, and then a hydrophobic barrier was drawn around the tissue sections using a Pap Pen (Vector Labs). Slides were then incubated with hydrogen peroxide solution for 10 min, washed in distilled water, and then treated with the Protease IV solution for 30 min at room temperature in a humidified chamber. Following protease treatment, C1 and C2 cDNA probe mixtures specific for mouse tissue were prepared at a dilution of 50:1, respectively, using the following probes from Advanced Cell Diagnostics: *Oprm1* (C1, 315841), *Fos* (C4, 316921), *Penk* (C2, 318761), *Vglut2* (C3, 456751) and *Vgat* (C3, 319191). Sections were incubated with cDNA probes (2 h), and then underwent a series of signal amplification steps using FL v2 Amp 1 (30 min), FL v2 Amp 2 (30 min) and FL v2 Amp 3 (15 min). 2 min of washing in 1x RNAscope wash buffer was performed between each step, and all incubation steps with probes and amplification reagents were performed using a HybEZ oven (ACD Bio) at 40° C. Sections then underwent fluorophore staining via treatment with a serious of TSA Plus HRP solutions and TSA Vivid dyes 520, 570, or and 650 fluorescent dyes at a dilution of 1:3000. All HRP solutions (C1-C2) were applied for 15 min and TSA Vivid dyes for 30 min at 40° C, with an additional HRP blocker solution added between each iteration of this process (15 min at 40° C) and rinsing of sections between all steps with the wash buffer. Lastly, sections were stained for DAPI using the reagent provided by the Fluorescent Multiplex Kit. Following DAPI staining, sections were mounted, and cover slipped using Fluoromount-G mounting medium and left to dry overnight in a dark, cool place. Sections from all mice were collected in pairs, using one section for incubation with the cDNA probes and another for incubation with the 4-Plex Negative Control probe (ACD Bio, 321831) to serve as a negative control.

### Imaging and Quantification

All tissue was imaged on a Keyence BZ-X all-in-one fluorescent microscope at 48-bit resolution using the following objectives: PlanApo-λ x4, PlanApo-λ x20 and PlanApo-λ x40. All image processing prior to quantification was performed with the Keyence BZ-X analyzer software (version 1.4.0.1). Quantification of neurons expressing fluorophores was performed via manual counting of TIFF images in Photoshop (Adobe, 2021) using the Counter function or using HALO software (Indica Labs), which is a validated tool for automatic quantification of fluorescently-labeled neurons in brain tissue^48,49,130–132^. Counts were made using 20X magnified z-stack images of a designated regions of interest (ROI). We used HALO software for all fluorescent *in situ* hybridization (FISH) quantifications. One representative 16 µm slice containing the PAG (selected from −4.6 to −4.8 relative to Bregma) was quantified per mouse, using HALO Image Analysis software (Indica Labs). The borders for left and right vlPAG, lPAG, dlPAG, and dmPAG were hand-drawn as individual annotation layers, using the Allen Brain Reference Atlas and *The Mouse Brain in Stereotaxic Coordinates*, 3^rd^ edition (Franklin and Paxinos) as guides. Slices were visually inspected for damage, dust or other debris, in addition to bound probe, and those areas with substantial damage or contamination were manually excluded from their respective annotation layers. Colocalization of nuclei (DAPI) with *Oprm1, Fos*, and *Vglut2, Vgat*, or *Penk* mRNA puncta was automatically quantified using the FISH module (version 3.2.3) and traditional nuclear segmentation. Setting parameters were optimized by comparing performance across 6 slices, randomly selected across experimental groups, and confirming proper detection by visual inspection. Identical parameters were applied across all slices processed through FISH on the same day.

### Single nuclei RNA sequencing

#### Nuclei preparation

A single punch of the periaqueductal gray (PAG) measuring 2 mm in width and 1 mm in depth, was used to prepare the nuclei suspensions. Slices were taken at coordinates approximately A/P −4.00 to −5.00 mm relative to Bregma. Nuclei isolation was performed using the Minute™ single nucleus isolation kit designed for neuronal tissue/cells (Cat# BN-020, Invent Biotechnologies), similar to our prior description^133–135^. Briefly, the tissue was homogenized in 500 µL of buffer A using a pestle in a 1.5 mL LoBind Eppendorf tube and then allowed in incubate on wet ice for 5 minutes.. The homogenate was then transferred to a filter column within a collection tube and incubated at −20°C for 10 minutes. Following this, the tubes were centrifuged at 13,000 x g for 30 seconds, the filter was discarded, the pellet resuspend, and the samples were centrifuged at 600 x g for 5 minutes. Supernatants were removed and the pellet underwent one wash with 200 μL of PBS + 5% BSA and then was resuspended in 60 μL of PBS + 1% BSA. The concentration of nuclei in the final suspension was assessed by staining with Trypan Blue and counting using a hemacytometer.

#### Single-Nuclei Gene Expression Assay

The nuclei suspensions were used for the 10x Genomics 3’ gene expression assay (v4), conducted following the instructions provided. The maximum number of nuclei for each sample, defined by concentration, was loaded into the 10x Genomics Chromium microfluidics controller,. Subsequently, sequencing libraries were constructed following the manufacturer’s protocol for unique dual Illumina indexes, and libraries containing unique indexes were pooled together at equimolar concentrations of 1.75 nM and sequenced on the Illumina NovaSeq 6000, using 28 cycles for Read 1, 10 cycles for the i7 index, 10 cycles for the i5 index, and 150 cycles for Read 2.

#### Data analysis

##### Processing and clustering of single nuclei data

Sequencing reads were processed to generate fastq files using 10x Genomics Cellranger v8.0.1, and reads were aligned to an mm10 genome optimized for single cell sequencing ^136^. Similar to our previous work^137^, filtered matrices for each individual sample were converted to Seurat objects using Seurat 5.0.1, and filtered to retain only nuclei with >200 minimum features and <5% mitochondrial reads. Initial dimensionality reduction and clustering was performed to enable adjustment of count matrices for cell-free mRNA using SoupX, with the contamination fraction set to 0.35^138^. The adjusted matrices were then filtered to retain only UMIs with >200 and <8k features, and <60k UMIs. SCTransform was used to normalize and scale expression data while regressing out the percentage of mitochondrial transcripts and IntegrateData was used to integrate data sets from all samples. scDblFinder v1.12.0 was used to identify doublets and all doublets were subsequently removed, as well as any clusters with expression of mixed cell type markers. High resolution clustering was then performed and additional residual putative multiplet clusters were removed. The final integrated and cleaned data set was clustered at a resolution of 0.3 using the first 20 principal components (PCs) to identify major cell types. Neuronal nuclei were subset and subclustered separately at a resolution of 1 using 40 newly calculated PCs.

##### Modular activity scoring

Modular activity scores were calculated in neuronal subclusters using 25 putative immediate early genes (*Arc, Bdnf, Cdkn1a, Dnajb5, Egr1, Egr2, Egr4, Fos, Fosb, Fosl2, Homer1, Junb, Nefm, Npas4, Nr4a1, Nr4a2, Nr4a3, Nrn1, Ntrk2, Rheb, Sgsm1, Syt4, Vgf*) against a control feature score of 5. IEG module score was visualized by violin plot. Module scores were also generated for each neuronal subcluster using known column marker genes (**Extended Data Fig. 4**^33^) and a control feature score of 5. Column marker module scores were visualized using Nebulosa^139^ plots generated with scPubR^140^.

### Behavioral testing

On test days, mice were brought into procedure rooms ∼1 h before the start of any experiment to allow for acclimatization to the testing environment. Mice were provided food and water ad libitum during this period. For multi-day testing conducted in the same procedure rooms, animals were transferred into individual “home away from home” (HAFH) cages ∼1 h prior to the start of testing and were only returned to their home cages at the end of the test day. All testing and acclimatization was conducted under red light conditions (5-30 lux). Equipment used during testing was cleaned with a 70% ethanol solution before starting, and in between, each behavioral trial to mask odors and other scents and remove animal waste. Behavioral tests were conducted by experimenters blinded to condition.

#### Sensory stimulus applications

To evaluate evoked responses to innocuous mechanical hindpaw stimulation during fiber photometry recordings, we used a 0.16 g von Frey filament. The filament was applied perpendicular to the plantar hindpaw with sufficient force to cause a slight bending of the filament. The filament was applied to the left hindpaw for approximately 2 seconds. To evaluate evoked responses to noxious mechanical stimulation during fiber photometry recordings, we used a sharp 25G syringe needle (pin prick). The pin prick was applied as a sub-second poke to the left hindpaw. To evaluate evoked responses to noxious thermal heat stimulation, a drop (approximately 25-50 µl) of 55 °C water was delivered to the left hindpaw, Onset of stimulus applications was annotated in the fiber photometry software Synapse using Arduino Uno trigger button connected via digital input to the RZ10X photometry processor. On testing days, mice were connected to the photometry system, and following a 10 min habituation period, recording sessions began. Each stimulus type was delivered to the left hindpaw 5 times with a 2-minute inter-stimulus interval. Between different stimulus types (0.16 g von Frey filament, 25G needle pin prick, 55 °C water droplet), the LED was turned off for 5 minutes to minimize photobleaching and animal stress.

#### Dynamic thermal plate assay

The dynamic thermal plate assay was used in conjunction with fiber photometry recordings to assess changes in nocifensive behavioral repertoires and neural activity as animals transitioned from an innocuous plate temperature to a noxious plate temperature (i.e., ≥42 °C). To do this, a controllable thermal plate (Bioseb) was programmed with integrated computer software to hold at 30 °C until the experimenter initiated a preset temperature ramp protocol with a mechanical foot pedal attached to the thermal plate. An Arduino Uno trigger button was used to time-lock the onset of the ramp in Synapse. The ramp protocol was defined to change the thermal plate temperature from 30 °C to 50 °C at a rate of 10 °C per minute. When 50 °C was achieved, animals remained on the plate for an additional 30 seconds prior to removal and return to HAFH cages. On each test day, mice were habituated to the 30 °C thermal plate for 5 minutes prior to the onset of the ramp protocol. For optogenetics experiments, a 60-second stimulation period (10 Hz, 5 ms pulsewidth, 5 mW) followed the 5-minute habituation period, just prior to the ramp onset. Video recordings were obtained for every test with a USB webcam and OBS software for post-hoc analysis of affective-motivational nocifensive behaviors (i.e., escape and attending). The following behaviors constituted escape behaviors: rearing on the wall of the thermal plate chamber and jumping. The following behaviors constituted attending behaviors: licking either hindpaw, guarding either hindpaw, and licking either forepaw. The affective-motivational behaviors were manually scored from the video recordings using the Behavioral Observation Research Interactive Software^58^ (BORIS) to time-lock onset of individual behaviors and quantify both the number and duration of each behavior. Overhead videos were obtained for optogenetics experiments to assess locomotor activity using Ethovision tracking software (Noldus). When applicable, morphine was injected 30 minutes prior to placing the mice in the thermal plate apparatus and naltrexone was administered immediately prior to placing mice in the apparatus.

#### Trace fear conditioning (TFC) and extinction

Mice underwent a 2-day TFC and extinction paradigm based on a previously described protocol from our group^141^. Briefly, a Med Associates operant box was equipped with fear conditioning peripherals (i.e., shock grid floor and tone generator) and the sound attenuating chamber was modified to accommodate fiber photometry patch cables. Acquisition and extinction occurred in the same fear conditioning box, however, environmental cues were altered to provide distinct contexts. Context A was defined by the presence of the shock grid floor suspended over a metallic tray filled with clean cage bedding and cleaned with Clidox-S disinfectant. Context B was defined by a smooth white floor insert covering the shock grid and was cleaned with ethanol. Acquisition occurred in Context A and Extinction occurred in Context B. For conditioning on the acquisition day, a train of 25 tones (4 kHz, 75 dB, 200 ms) delivered at 1 Hz was used as the conditioned stimulus (CS). The presentation of the CS was followed by a period of quiescence for 20 seconds (trace). The trace period was then followed by the unconditioned stimulus (US), a footshock (1 mA, 2 s). On the extinction day, the footshock was omitted. During both acquisition and extinction days, 10 trials occurred that were separated by an inter-trial interval of 60 ± 10 seconds. Presentation of all stimuli was annotated in Synapse via digital inputs from the Med Associates box to the RZ10X photometry processor.

#### Placebo analgesia conditioning (PAC) assay

To elicit placebo analgesia-like behavior in mice, we used a modified version of an existing protocol^83^. The PAC paradigm involved a 7-day procedure using two contiguous, controllable thermal plates and an acrylic plastic chamber creating two interconnected compartments via a partially open central divider and distinct contextual cues in either compartment (i.e., vertical vs. horizontal black and white stripes). The divider remained open throughout testing to allow animals to move freely between the two chambers. Prior to placement in the apparatus, mice were placed in a holding chamber for minutes to acclimate to tethering to the patch cable. On days 1 and 2, animals underwent two habituation sessions, 5 minutes each, during which the thermal plates were set to 30 °C. All subsequent sessions were 3 minutes in duration. For pre-testing on day 3, both thermal plate temperatures were set to 45°C. On days 4-6, twice-daily conditioning sessions were performed. For animals assigned to the Control group, both the left (Side A) and right (Side B) compartments were set to 30 °C. For animals assigned to the Conditioned group, Side A was set to 30 °C and Side B was set to 48°C. Within each conditioning day, the conditioning sessions were separated by 4 hours. During post-testing on day 7, the thermal plates in both compartments were set to 45°C for all animals. Overhead tracking was collected using a machine vision camera (Basler) integrated with Ethovision for movement and preference quantification. Mice exhibiting >85% preference for either compartment during the Pre-Test were excluded from analysis. Two front-facing webcams were used to record affective-motivational nocifensive behaviors through OBS software during the Post-Test. The webcam videos were analyzed post-hoc using the BORIS event-logging software. The following behaviors constituted escape behaviors: rearing on the walls of the apparatus, jumping, crossing the center to exit/enter either compartment and digging along the bottom of the chamber wall. The following behaviors constituted attending behaviors: licking either hindpaw, guarding either hindpaw, and postural extension. Thermal plate temperatures were controlled with an Arduino Uno microcontroller. An Arduino Uno trigger button was used to log the start of the Post-Test in Synapse and provide visual syncing of video recordings to photometry recordings.

### *In vivo* fiber photometry

Optical recordings of GCaMP6f and δLight fluorescence were acquired using an RZ10x fiber photometry detection system (Tucker-Davis Technologies), consisting of a processor and integrate Synapse software (Tucker-Davis Technologies), and optical components (Doric Lenses and ThorLabs). Excitation wavelengths generated by LEDs (465 nm blue light and 405 nm violet light) were relayed through a filtered fluorescence minicube at spectral bandwidths of 460–495 and 405 nm to a pre-bleached, low auto-fluorescence, mono fiberoptic patch cord connected to the implant on top of each animal’s head. Power output for the primary 465 nm channel at the tip of the fiber optic implant was adjusted to ∼50 μW. Signal in both 465 and 405 nm channels was monitored continuously throughout all recordings, with the 405 nm signal used as the isosbestic control and for correcting motion artifacts introduced by movement of components in the light path. Wavelengths were modulated at lock-in amplification frequencies of 327 Hz (405 nm) and 531 Hz (465 nm). All signals were acquired at 1 kHz and lowpass filtered at 4 Hz. Following testing, all mice were perfused, and the tissue was assessed for proper viral targeting and transduction efficacy, as well as fiber optic placement via immunohistochemistry.

#### GCaMP6f analysis

Analysis of the GCaMP signal was performed with the open source, MATLAB-based photometry modular analysis tool pMAT^142^. The MATLAB polyfit function was used to correct for photobleaching and motion artifacts for the duration of each recording, using linear least squares fitting of the 405 nm signal to the 465 nm signal and subtraction of the fitted 405 nm signal from the 465 nm signal. GCaMP fluorescence was determined as a change in the 465 nm divided by the fitted 405 signal (ΔF/F). The resulting ΔF/F was down-sampled by a factor of 100 and the perievent time histogram (PETH) module in pMAT was used to analyze time-locked epochs of the recordings. Time-locked behavioral events were captured by an Arduino Uno transistor-transistor logic (TTL) input. Baseline Z-score calculations were calculated from the PETH ΔF/F values. Peak Z-score and area under the curve were determined from these PETH analyses across groups. Z-scored traces were plotted in Prism (GraphPad).

#### δLight analysis

For short-window (< 2 minutes) analyses of δLight signal, the pMAT PETH module was used, as described above. In all other experiments using δLight or δLight0, where the entire length of the recording was analyzed, nonlinear least squares was calculated using a double exponential fit of the pre-injection or pre-behavior onset baseline period (5-15 min) of both the 405 nm and 465 nm channels in MATLAB, similar to that described previously^63,143^. The fitted 405 nm signal was subtracted from the 465 nm signal to detrend slow photobleaching artifacts and correct for motion artifacts, followed by ΔF/F calculation and down-sampling by a factor of 300 to reduce high frequency noise. Baseline Z-scoring was performed on the detrended ΔF/F trace.

### *In vivo* optogenetic activation

Optogenetic activation of *Penk* neurons was chosen to be unilateral to limit structural damage of the PAG to one hemisphere and correspond with fiber photometry experiments. For fiber photometry recordings, the right vlPAG was infused with Cre-dependent ChrimsonR virus 1 week prior to infusion of δLight and fiber optic implantation (400 μm diameter, Doric) over the right vlPAG. For behavioral experiments, Cre-dependent ChRmine infusion was immediately followed by fiber optic implantation over the vlPAG in the same surgery. Optogenetic stimulation was delivered with an orange/red LED (630 nm) via a patch cable. Power output at the fiber optic tip was measured to be ∼ 5 mW.

### Drugs

All drugs were administered either via the intraperitoneal (i.p.) or subcutaneous (s.c.) route at 10 ml/kg, according to the bodyweight of each mouse. 4-hydroxytamoxifen (4-OHT; HelloBio, HB6040) was dissolved in 100% ethanol (for a 10-mg bottle: 250 μL of ethanol) on the morning of use. The solution was heated at 50°C for up to 30 min to encourage dissolving. The 4-OHT solution was further diluted in Kolliphor EL (Sigma Aldrich, C5135-500G; 500 μL for a 10-mg bottle) and finally 1X phosphate buffered saline (PBS; Calbiochem, 524650; 1.75 ml for a 10-mg bottle) when it was delivered s.c. at a dose of 40 mg/kg body weight. The following opioid receptor ligands were dissolved in 0.9% saline to achieve final working solutions: morphine sulfate (10 mg/ml stock, Covertus) was diluted to 1 mg/ml for an injected dose of 10 mg/kg, i.p.; naltrexone hydrochloride (HelloBio, HB2452) was dissolved to a working concentration of 0.1 mg/ml for an injected dose of 1 mg/kg, i.p.; naloxone hydrochloride (HelloBio, HB2451) was dissolved to a working concentration of 0.4 mg/ml for an injected dose of 4 mg/kg, i.p. The δ opioid receptor agonist SNC 162 (Tocris, 15-291-0) was first dissolved in 1 M hydrochloric acid to a stock of 10 mg/ml and then subsequently diluted to final working solutions of 0.5 mg/ml (5.0 mg/kg, i.p. dose) or 0.25 mg/ml (2.5 mg/kg, i.p. dose). The vehicle control for SNC 162 consisted of 1 M HCl diluted with saline to 2% of the working volume.

### Statistics and data presentation

Group sizes were based off of published literature for the type of manipulation and measured outcome published in the field and/or by the authors involved. Sample sizes for all studies are included in the main text and in the statistics summary table (**Extended Data Table 1**). In order to reduce our animal usage and to account for reported sex differences in pain in humans, male and female mice were used throughout experiments. When male and female mice were used, an analysis for sex differences in the data was performed. For many of the studies, multiple cohorts were used in the final group sizes. All behavior results were consistent and replicated across cohorts. Individual data points are included and indicate consistent trends across mice in each behavior study. Mice were randomly assigned into control or experimental groups to the best of the experimenter’s abilities, with counterbalancing for age and sex as needed. Once experimental/control groups were formed to comprise the studies cohort of mice, the cohort underwent all behavioral testing concurrently and experimenters were blinded to initial condition for analysis. Representative images of histology were selected to display viral spread, fiber optic placements, and endogenous fluorescence. For all imaging and behavioral studies, virus injected animals with either little or no evidence of viral transduction and/or incorrect viral targeting were excluded from any final analyses. In cases of unidentifiable fiber optic cannula tracts, subjects were retained if viral payload was correctly delivered to vlPAG (n=1 across all fiber photometry experiments; **Extended Data Fig. 17**). All data are presented as mean ±the standard error of the mean (SEM) for each group, and all statistical analyses were performed using Prism 9 & 10 software (GraphPad). Statistical tests used throughout this paper include paired and unpaired t-tests, Pearson’s correlations, and one- and two-way ANOVAs. When post-hoc testing was appropriate, following a significant ANOVA test, the Bonferroni correction was applied to all data. Data was represented using Prism and figures were finalized in Illustrator (Adobe). Significance was determined with a p-value of <0.05. All figures with star indicate comparisons that produced a statistical p-value less than 0.05. All significant p-values visualized in the main and supplemental figures are reported in **Extended Data Table 1**.

## EXTENDED DATA for

**This file includes:**

1. Extended Data Figures 1-17
2. Extended Data Table 1

**Extended Data Figure 1.**
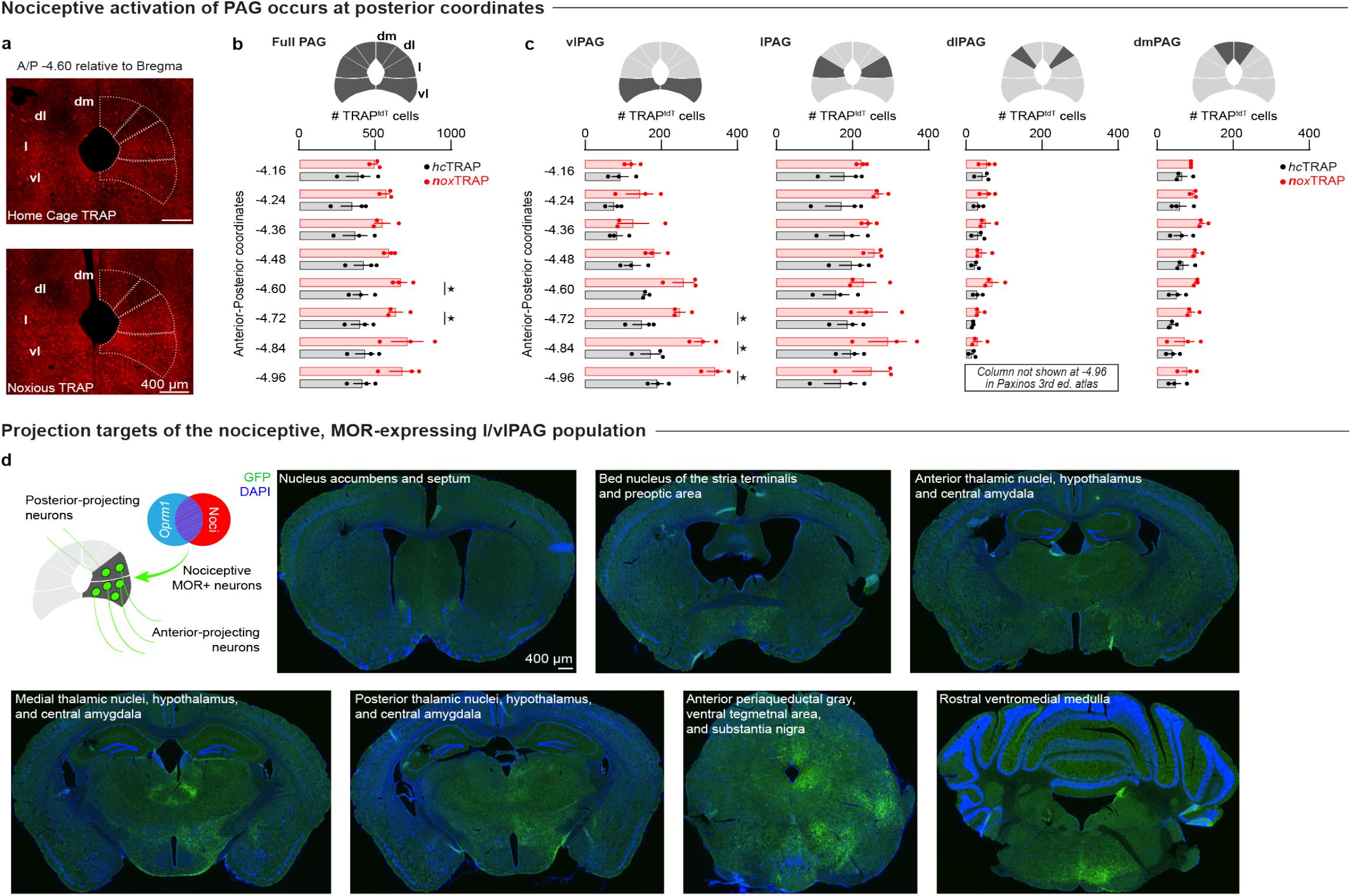
Posterior vlPAG neurons are activated by acutely noxious stimulation and projects to other affective-motivational brain areas. **a**. Representative 20x images of home cage TRAP (*hc*TRAP) and noxious 55 °C water-stimulated TRAP (*nox*TRAP) PAG slices at approximately A/P −4.60 relative to Bregma. **b**. Comparison of total TRAP-tdTomato (TRAP^tdT^) counts across all PAG columns combined between *hc*TRAP and *nox*TRAP groups. **c**. Comparison of total TRAP^tdT^ counts for vlPAG, lPAG, dlPAG, and dmPAG, respectively, between *hc*TRAP and *nox*TRAP groups. **d**. Representative 4x images of brain areas with GFP+ axonal projections from nociceptive, *Oprm1*+ vlPAG neurons related to Figure 1c.

**Extended Data Figure 2.**
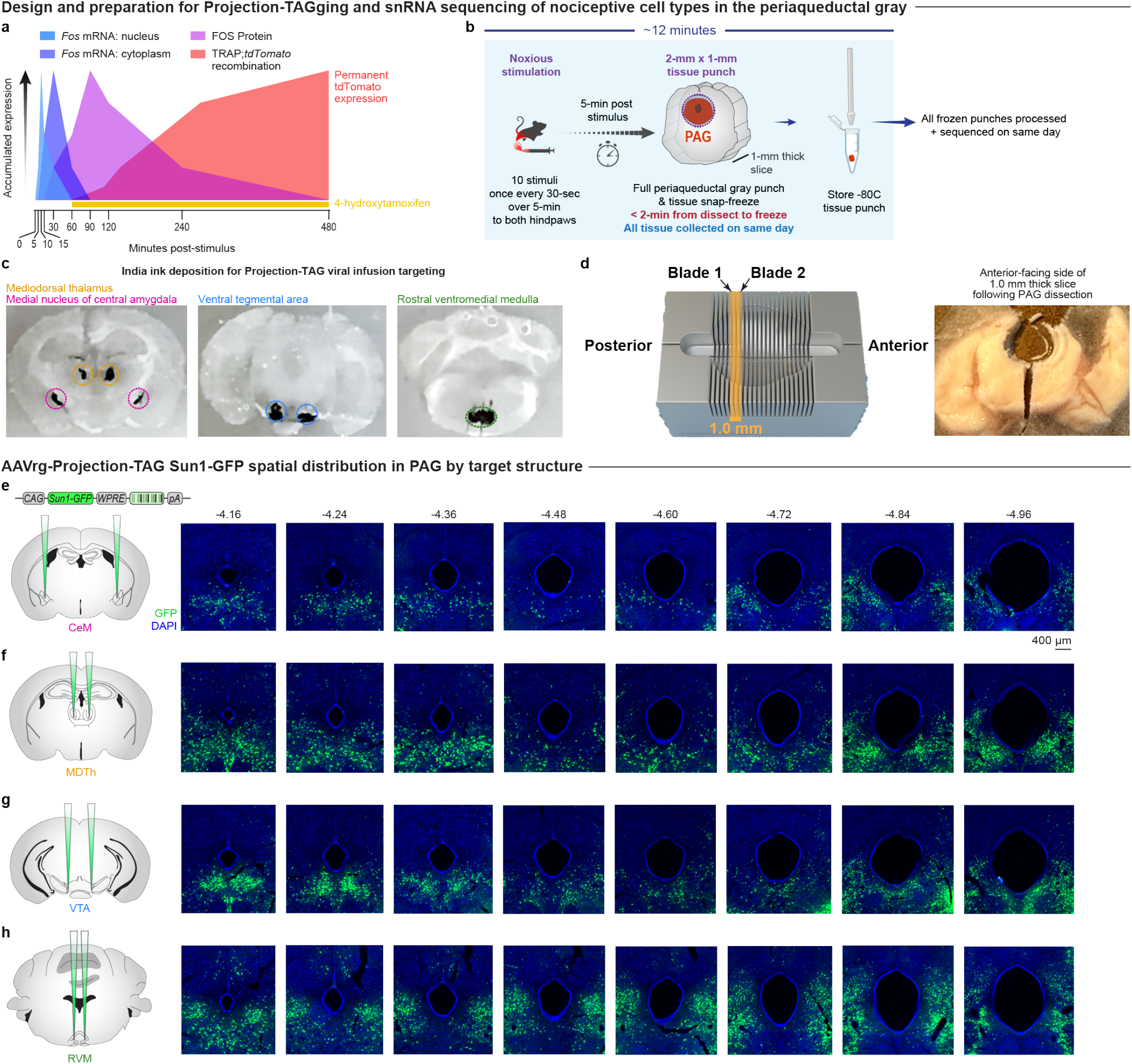
Details of tissue collection for single nucleus RNA sequencing and PAG Projection TAGging. **a**. Time of Fos mRNA and FOS protein accumulation in the nucleus and cytoplasm of the cell used for designing tissue collection timing across all experiments. **b**. Timeline and schematic of tissue collection for single nucleus RNA sequencing of PAG. **c**. Example mouse used for establishing viral injection coordinates for each Projection-TAG; locations of India ink deposition are circled according to the Projection-TAG that would be injected into each region. **d**. Didactic for approximate location of 1-mm thick PAG slice collection in a brain sectioning matrix with example image of a punch made of the PAG after slicing and flash freezing. **e-h**. Representative 4x images of PAG demonstrating Sun1-GFP signal expressed by the AAVrg-*CAG*-Projection-TAG injected into medial nucleus of the central amygdala (CeM, **e**), mediodorsal thalamus (MDTh, **f**), ventral tegmental area (VTA, **g**), rostral ventromedial medulla (RVM, **h**).

**Extended Data Figure 3.**
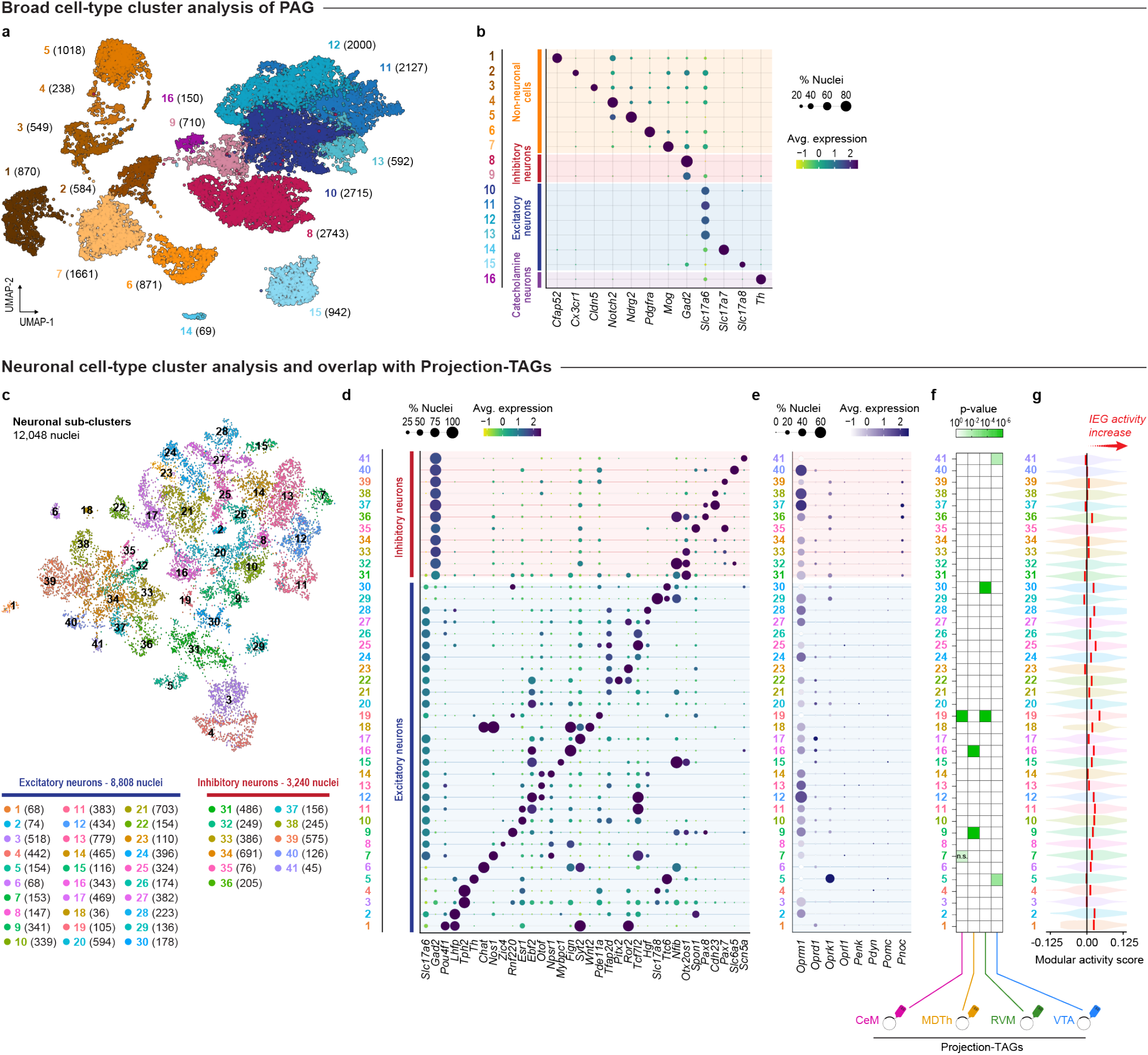
Single nucleus RNA sequencing broad cell-type and neuronal cluster analyses. **a**. Uniform Manifold Approximation and Projection (UMAP) of all nuclei (n=17,839) from the 3 noxious 55 °C water-stimulated samples (N=2 PAG punches per sample). **b**. Dotplot displaying major cell-type marker genes differentiating the broad cell-types. **c**. UMAP of neuronal nuclei (n=12,048) sub-clusters. **d**. Dotplot displaying a top gene differentiating each neuronal sub-cluster and categorization of each sub-cluster as excitatory or inhibitory neurons. **e**. Dotplot of opioid receptor and opioid peptide genes expressed in each neuronal sub-cluster. **f**. Heatmap of p-values demonstrating neuronal sub-clusters with significant levels of Projection-TAG barcode transcripts. **g**. Violin plot displaying the modular activity scores for immediate early gene expression across all neuronal sub-clusters.

**Extended Data Figure 4.**
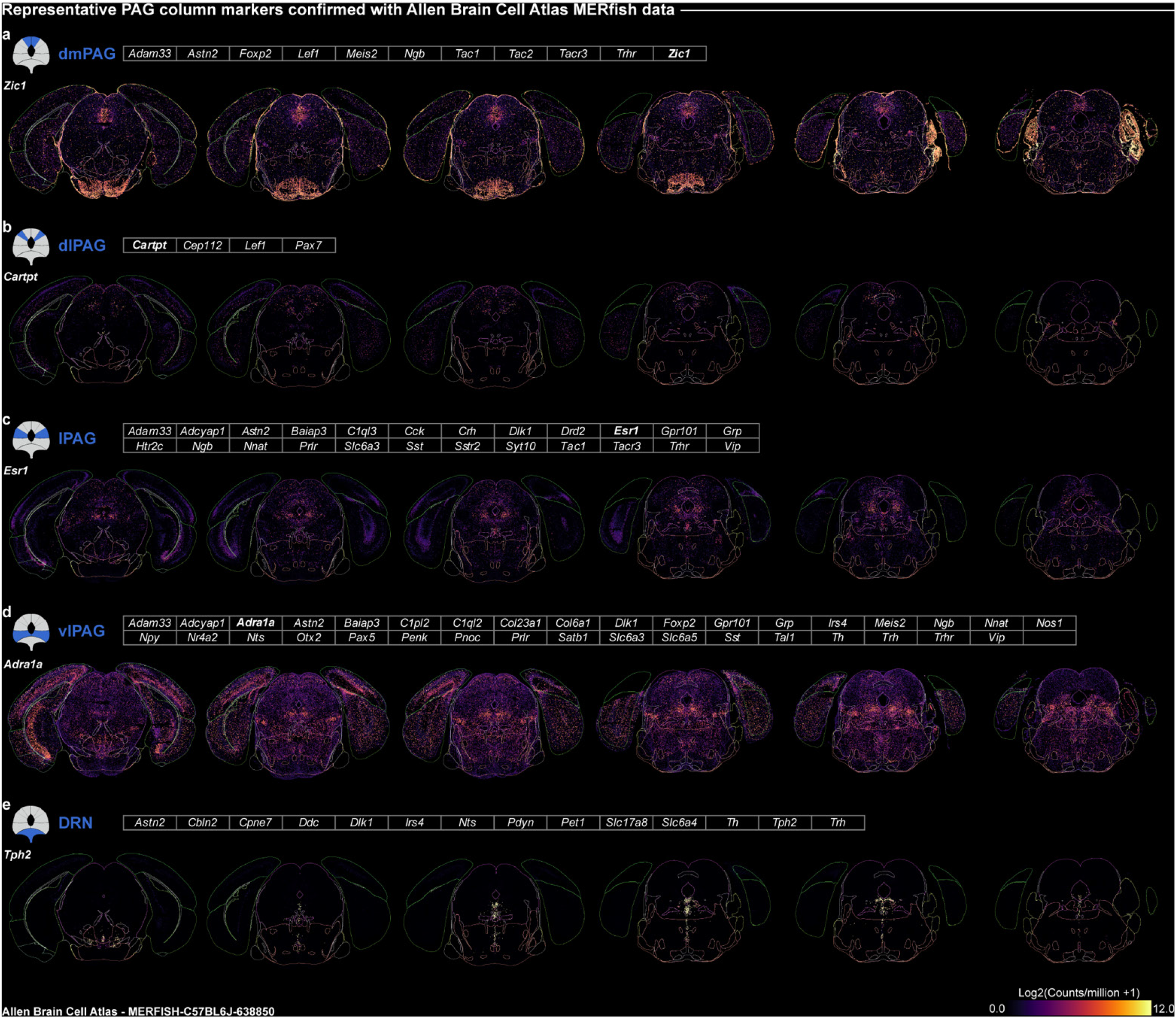
Modular stratification of PAG columns for single nucleus RNA sequencing analysis. **a-e**. Representative multiplexed error-robust fluorescence *in situ* hybridization (MERFISH) images from the Allen Brain Cell Atlas database (https://portal.brain-map.org/atlases-and-data/bkp/abc-atlas; MERFISH-C57BL6J-638850) demonstrating expression of select genes in dorsomedial (dm; **a**), dorsolateral (dl; **b**), lateral (l; **c**), and ventrolateral (vl; **d**) PAG columns in addition to the dorsal raphe nucleus (DRN; **e**). PAG column gene modules used to prepare density feature plots in Figure 1i are shown in table format.

**Extended Data Figure 5.**
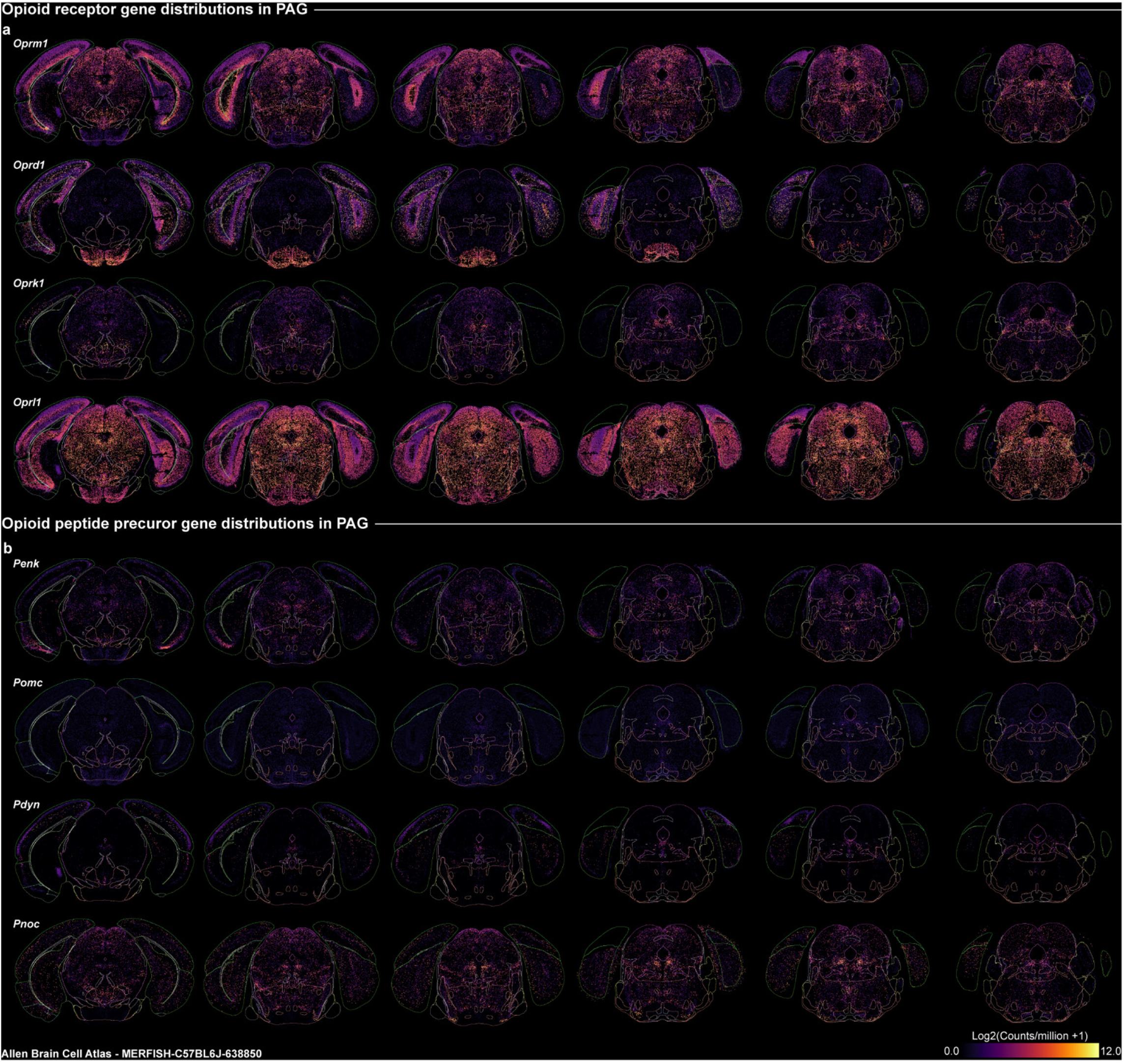
Opioid receptor and peptide gene expression in PAG. **a-b**. Representative MERFISH images from the Allen Brain Cell Atlas database (https://portal.brain-map.org/atlases-and-data/bkp/abc-atlas; MERFISH-C57BL6J-638850) demonstrating expression of opioid receptor genes (**a**) and opioid peptide precursor genes (**b**) across PAG columns and anterior-posterior coordinates.

**Extended Data Figure 6.**
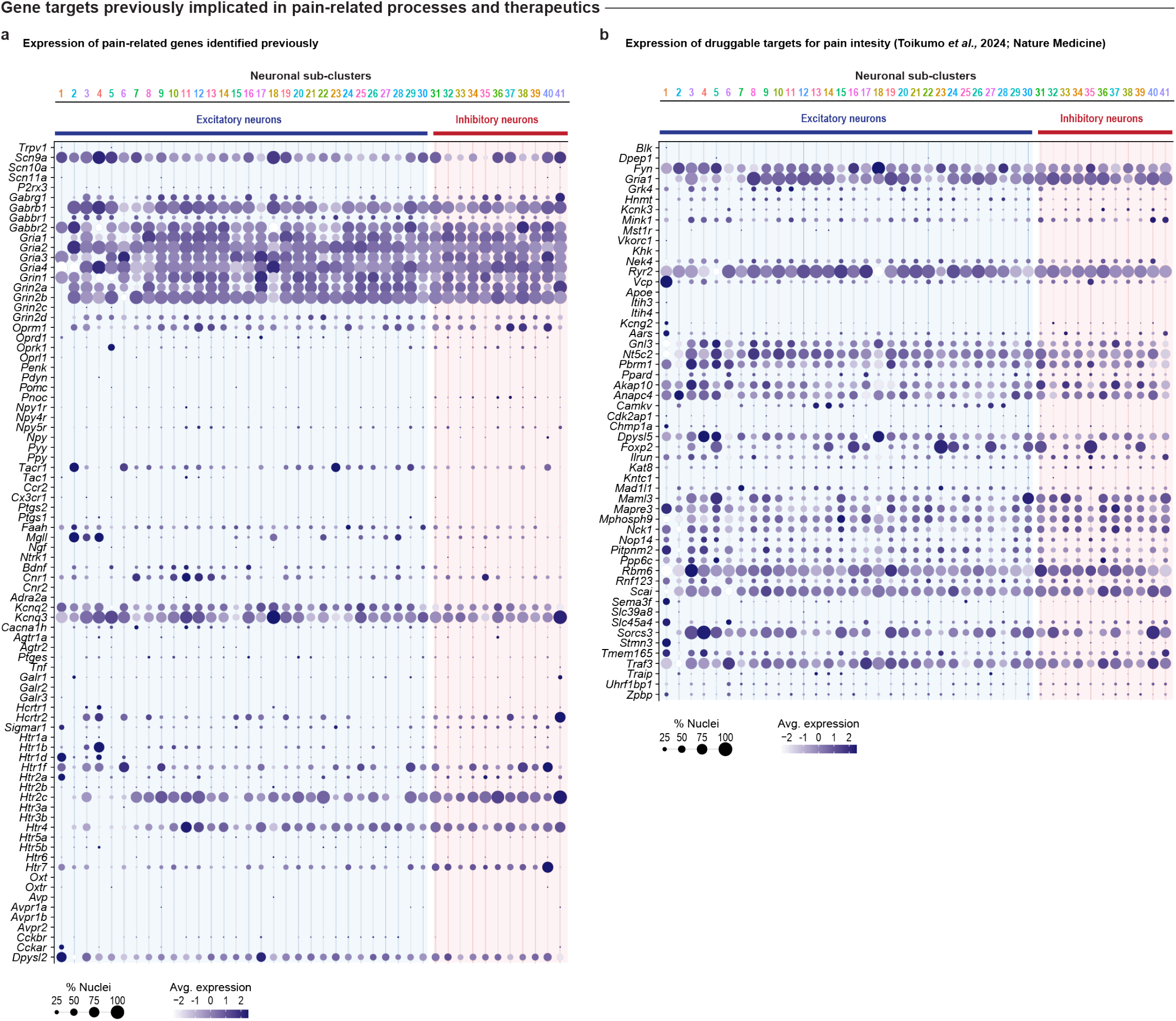
Expression of pain-related and therapeutic gene targets in PAG. **a**. Dotplot displaying top genes implicated in pain processes and their expression levels across all PAG neuronal sub-clusters. **b**. Dotplot displaying top genes identified as druggable targets for pain intensity by the Druggable Genome and Drug Gene Interaction Database and published as supplementary material in Toikumo et al., 2024, *Nature Medicine*, PMID: 38429522.

**Extended Data Figure 7.**
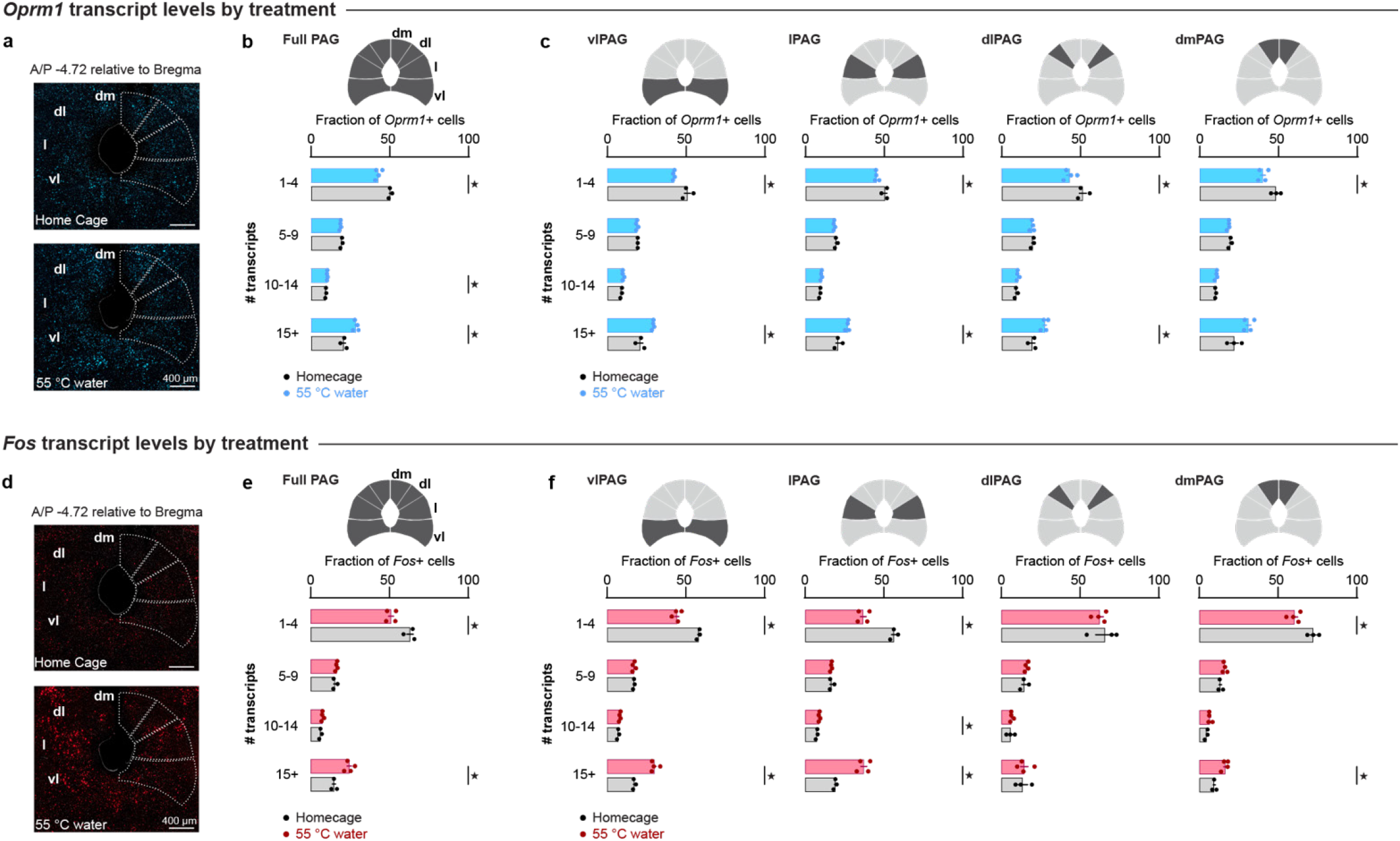
*Oprm1* and *Fos* transcript levels by PAG column and treatment group. **a**. Representative 20x fluorescence *in situ* hybridization images demonstrating mRNA transcripts for *Oprm1* in a home cage control mouse and a noxious 55 °C water-stimulated mouse at approximately A/P −4.72 relative to Bregma. **b**. Comparison of *Oprm1*+ nuclei expressing low (1-4), moderate (5-9), high (10-14), and very high (15+) levels of *Oprm1* transcript across all PAG columns between control and noxious 55 °C water-stimulated mice. **c**. Individual column comparisons of *Oprm1* transcript levels within *Oprm1*+ nuclei between control and noxious 55 °C water-stimulated mice. **d**. Representative 20x fluorescence *in situ* hybridization images demonstrating mRNA transcripts for *Fos* in a home cage control mouse and a noxious 55 °C water-stimulated mouse at approximately A/P −4.72 relative to Bregma. **e**. Comparison of *Fos* transcript levels within *Fos*+ nuclei between control and noxious 55 °C water-stimulated mice. **f**. Individual column comparisons of *Fos* transcript levels within *Fos*+ nuclei between control and noxious 55 °C water-stimulated mice.

**Extended Data Figure 8.**
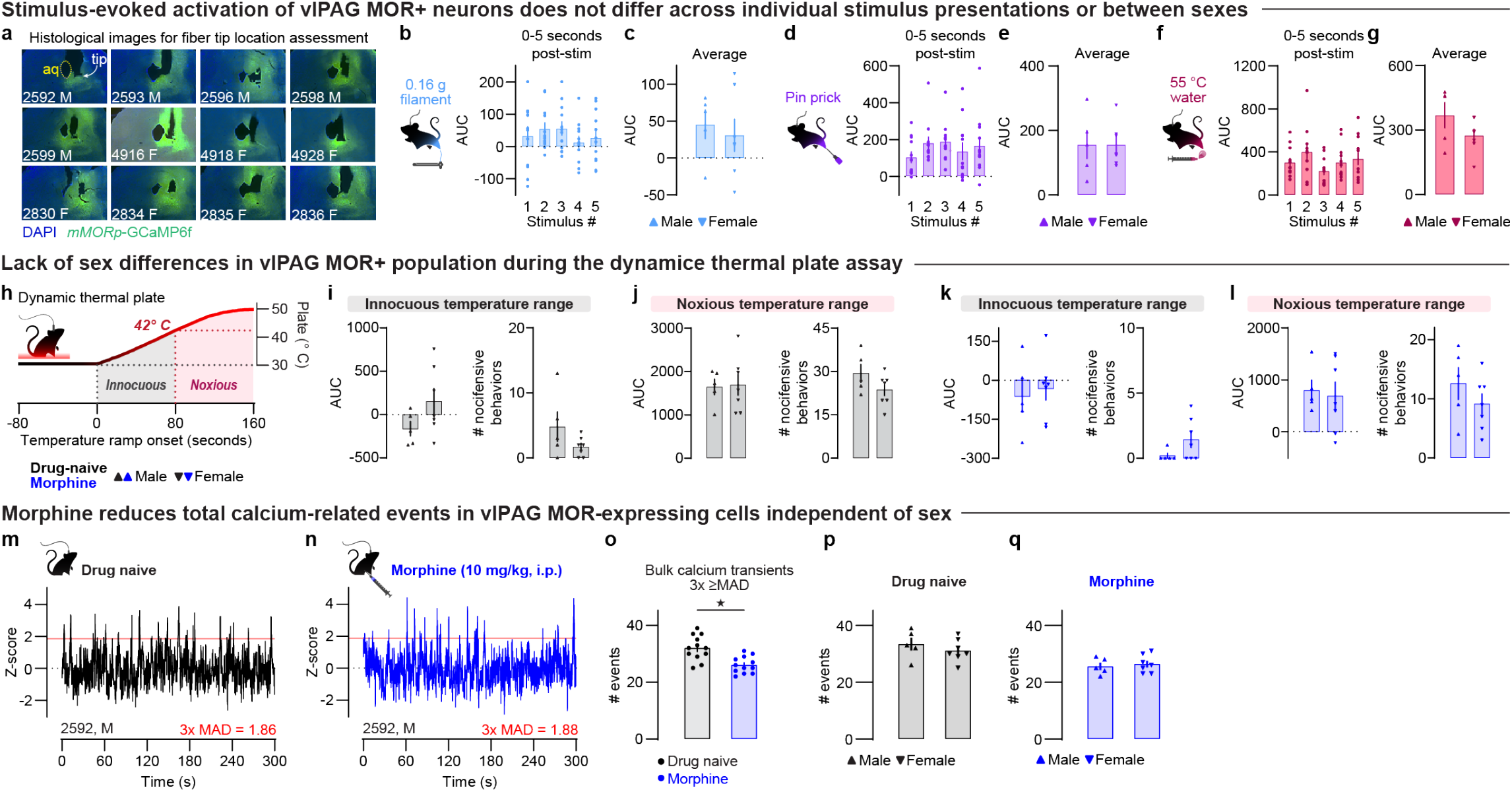
Nociceptive and opioid-related changes in vlPAG MOR+ population activity does not differ between sexes. **a**. Representative 4x images of fiber placements and viral *mMORp*-GCaMP6f expression in right vlPAG related to Figure 2; white arrow in M1 denotes fiber tip location and yellow dashed circle indicates the cerebral aqueduct (aq). **b**. MOR+ population responses to light touch to the left hindpaw with a 0.16g von Frey filament across stimulus applications. **c**. Comparison of light touch-evoked MOR+ population response by sex. **d**. MOR+ population responses to noxious pin prick to the left hindpaw with a 25 gauge needle across stimulus applications. **e**. Comparison of noxious pin prick-evoked MOR+ population response by sex. **f**. MOR+ population responses to noxious 55 °C water to the left hindpaw across stimulus applications. **g**. Comparison of noxious 55 °C water-evoked MOR+ population response by sex. **h**. The dynamic thermal plate assay assesses changes in MOR+ population calcium-related responses across a range of temperatures, both innocuous (<42 °C) and noxious (≥42 °C), and treatment conditions (e.g. drug naïve vs. morphine-treated). **i**. Comparison of drug-naive bulk fluorescence (area under the curve, AUC; left) and total nocifensive behavior count (right) by sex at innocuous temperatures. **j**. Comparison of drug-naïve AUC (left) and total nocifensive behavior count (right) by sex at noxious temperatures. **k**. Comparison of morphine-treated AUC (10 mg/kg, i.p.; left) and total nocifensive behavior count (right) by sex at innocuous temperatures. **l**. Comparison of morphine-treated AUC (left) and total nocifensive behavior count (right) by sex at noxious temperatures. **m**. Detection of spontaneous calcium-related events in vlPAG was determined by calculating the median absolute deviation (MAD) of the recording baseline and using 3x MAD as the cutoff for event detection above noise. An example drug-naïve trace is shown. **n**. Calcium event transient detection in the same mouse treated with morphine (10 mg/kg, i.p.). **o**. Comparison of drug-naïve and morphine-treated events. **p**. Comparison of events detected in the drug-naïve condition by sex. **q**. Comparison of events detected in the morphine-treated condition by sex.

**Extended Data Figure 9.**
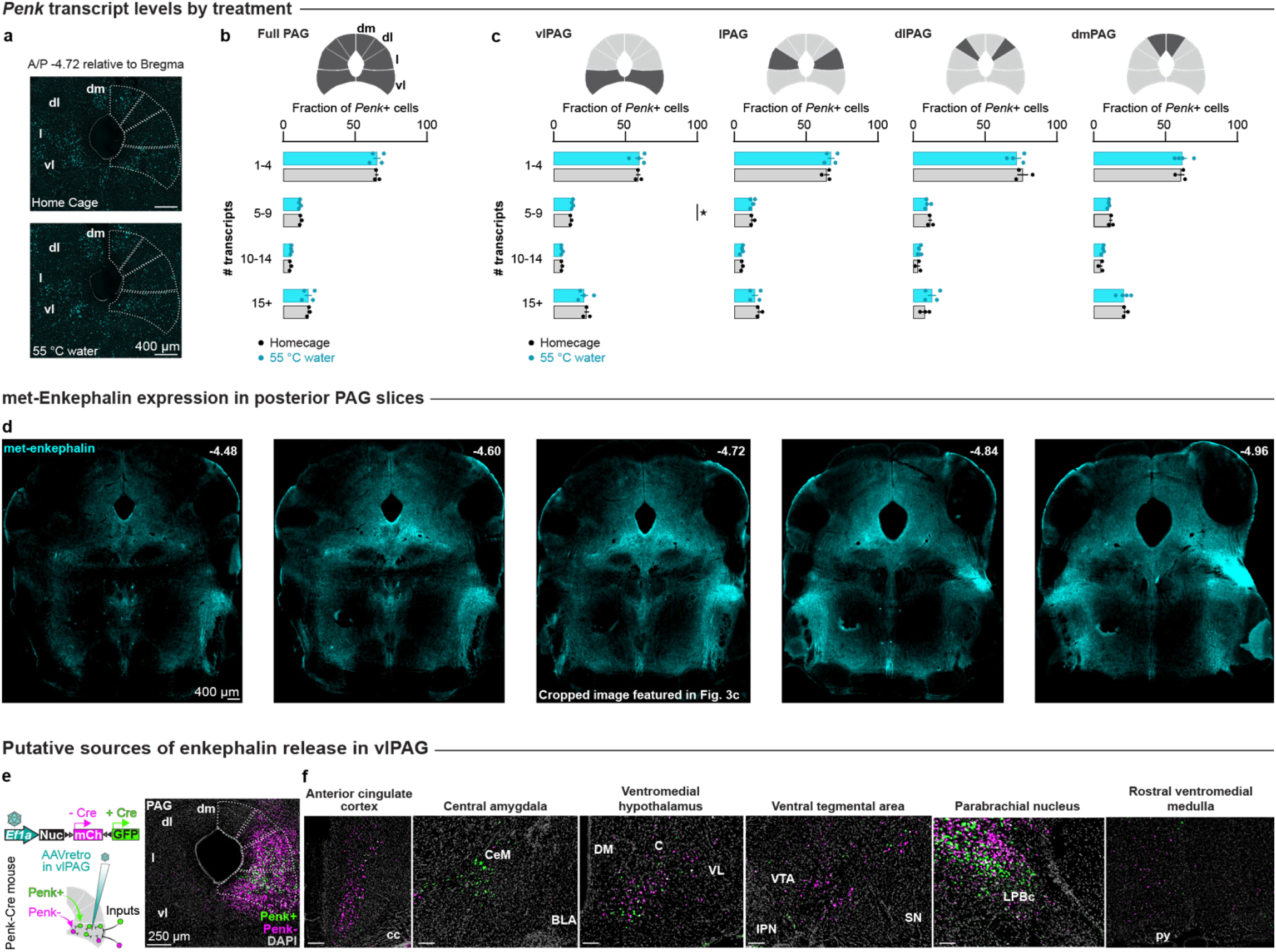
Enkephalinergic innervation of the PAG. **a**. Representative 20x fluorescence *in situ* hybridization images demonstrating mRNA transcripts for *Penk* in a home cage control mouse and a noxious 55 °C water-stimulated mouse at approximately A/P −4.72 relative to Bregma. **b**. Comparison of *Penk*+ nuclei expressing low (1-4), moderate (5-9), high (10-14), and very high (15+) levels of *Penk* transcript across all PAG columns between control and noxious 55 °C water-stimulated mice. **c**. Individual column comparisons of *Penk* transcript levels within *Penk*+ nuclei between control and noxious 55 °C water-stimulated mice. **d**. Representative 4x images of met-enkephalin immunostaining in posterior PAG sections related to Fig. 3c. **e**. Retrograde-AAV mapping approach for identifying enkephalinergic inputs to vlPAG (left) and representative 20x viral expression in vlPAG (right). **f**. Detection of enkephalinergic (GFP-labeled; green) and non-enkephalinergic (mCherry-labeled; magenta) cells that project to vlPAG in the anterior cingulate cortex (ACC), medial nucleus of the central amygdala (CeM), ventromedial nuclei of the hypothalamus (VMH), ventral tegmental area (VTA), parabrachial nucleus (PBN), and rostral ventromedial medulla (RVM).

**Extended Data Figure 10.**
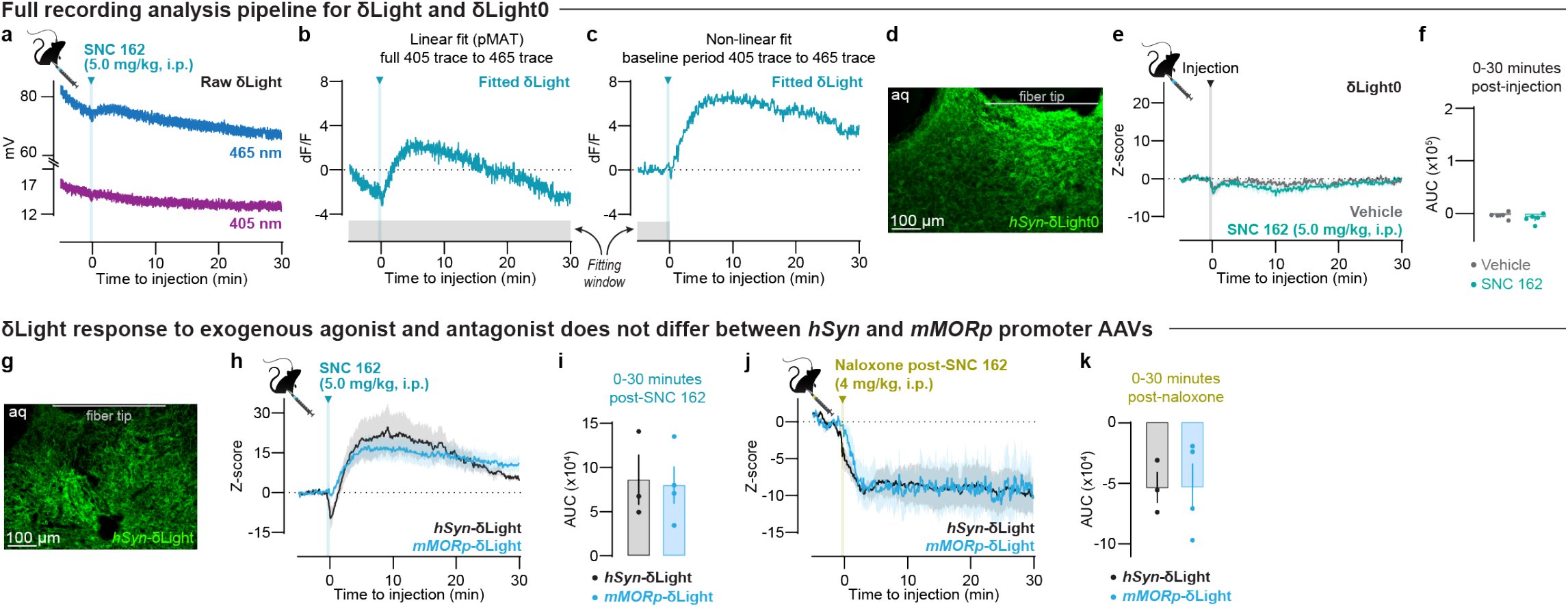
Details of δLight and δLight0 recording analysis and response to exogenous ligands. **a**. Representative raw 465 nm (signal) and 405 nm (isosbestic) traces from a mouse expressing δLight in vlPAG that received an injection of the δ opioid receptor agonist SNC 162 (5.0 mg/kg, i.p.). **b**. Linear fit calculation using the entire 405 nm and 465 nm traces in the Photometry Modular Analysis Tool (pMAT), followed by subtraction of the fitted isosbestic from the excitation signal, revealed a persistent downward trend in the resultant ΔF/F trace. **c**. Non-linear, exponential fitting calculated using the pre-injection 405 nm and 465 nm traces, followed by subtraction of the fitted isosbestic from the excitation signal, mitigated the slow decay in δLight-related ΔF/F. **d**. Representative 20x fluorescence image demonstrating viral transduction of AAV1-hSyn-δLight0 and fiber optic tip location in vlPAG. **e**. Fluorescence responses of δLight0 to vehicle and SNC 162 (5.0 mg/kg, i.p.). **f**. Quantified δLight0 bulk fluorescence change (AUC) in response to either vehicle or the δ opioid receptor agonist SNC 162 (5.0 mg/kg, i.p.). **g**. Representative 20x image of AAV1-hSyn-δLight in vlPAG with fiber optic tip location noted. **h**. Average Z-scored responses to SNC 162 (5.0 mg/kg, i.p.) recorded from mice expressing δLight under the control of either the *hSyn* or *mMOR* promoter. **i**. Comparison of *hSyn*-δLight and *mMORp*-δLight fluorescence response to SNC 162. **j**. Average Z-scored responses to naloxone (4.0 mg/kg, i.p.) given 45 minutes after SNC 162 (5.0 mg/kg, i.p.) recorded from mice expressing δLight under the control of either the *hSyn* or *mMOR* promoter. **k**. Comparison of δLight bulk fluorescence response to naloxone by promoter.

**Extended Data Figure 11.**
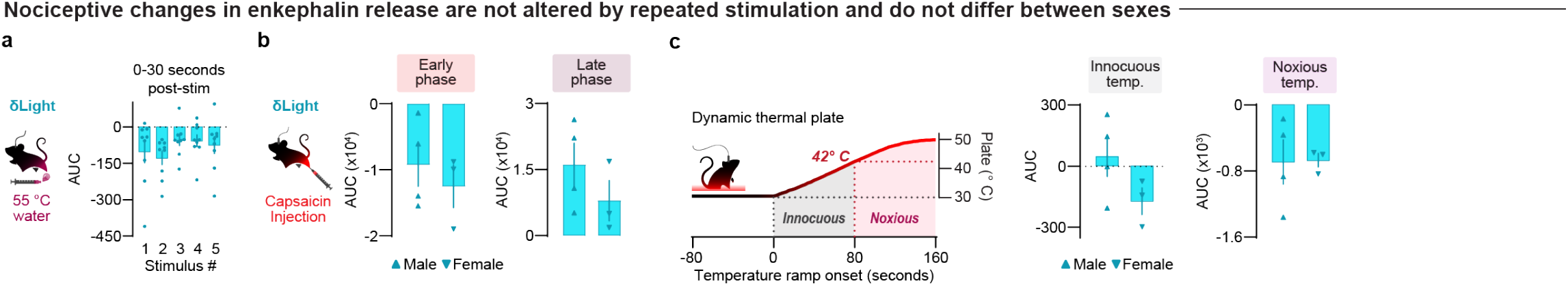
Nociceptive changes in vlPAG enkephalin are the same across stimulus presentations and sex. **a**. Comparison of noxious 55 °C water application to the hindpaw across each individual stimulus presentation. **b**. Comparison of δLight bulk fluorescence (AUC) in response to left plantar hindpaw capsaicin (10 µg) administration with respect to sex at early (5-15 min post-injection; left) and late (50-60 min post-injection; right) timepoints relative to injection onset. **c**. Innocuous (left) and noxious (right) temperature comparison of δLight bulk fluorescence by sex in the dynamic thermal plate assay.

**Extended Data Figure 12.**
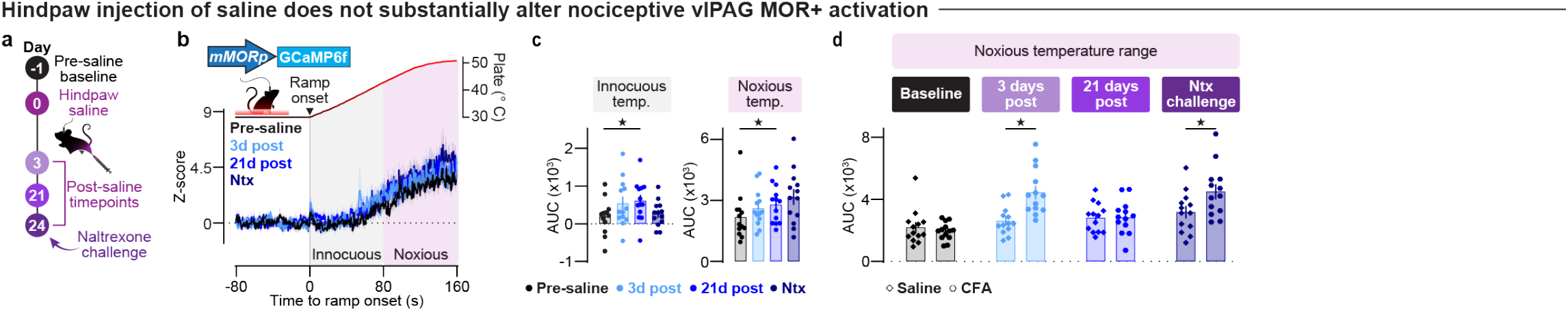
Hindpaw saline injection does not substantially alter noxious thermal heat-related nociceptive responses in vlPAG MOR-expressing cells. **a**. Timeline of dynamic thermal plate assay test days prior to and following hindpaw plantar injection of saline. **b**. Bulk calcium-related fluorescence in the vlPAG MOR+ population during the dynamic thermal plate assay at pre- and post-saline injection timepoints. **c**. Comparison of *mMORp*-GCaMP6f AUC at pre- and post-saline injection timepoints in the innocuous (left) and noxious (right) temperature ranges of the dynamic thermal plate assay. **d**. Noxious temperature range comparisons of Saline vs. CFA group AUC across each pre- and post-hindpaw injection timepoints.

**Extended Data Figure 13.**
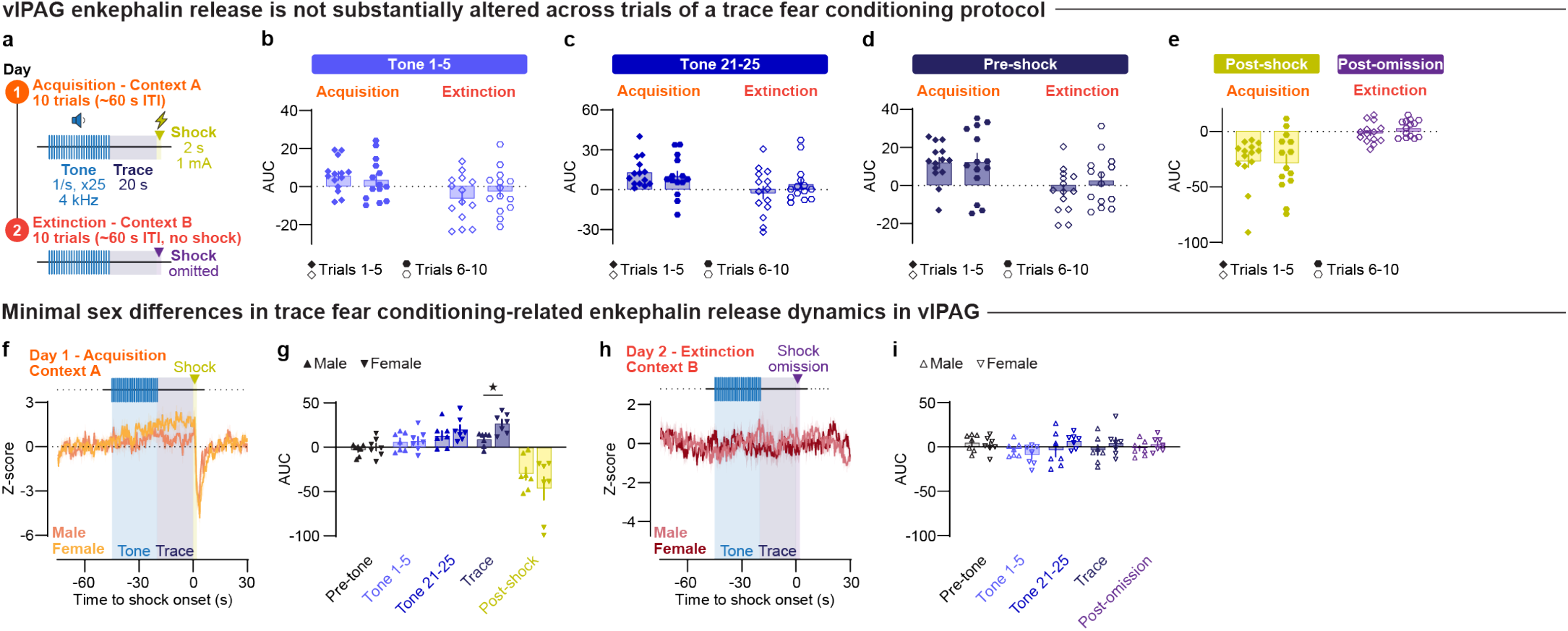
vlPAG enkephalin release does not substantially differ across fear conditioning trials or by sex. **a**. Didactic for the trace fear conditioning behavioral paradigm. **b**. Comparison of early (1-5) and late (6-10) trials during the initial tone presentations (Tone 1-5) within acquisition (left) and extinction (right) test days. **c**. Comparison of early and late trials during the final tone presentations (Tone 21-25) within acquisition (left) and extinction (right) test days. **d**. Comparison of the 5-second window prior to shock or omission during the trace period (Pre-shock) within acquisition (left) and extinction (right) test days, respectively. **e**. Comparison of the 5-second window immediately following the 2-second shock or equivalent period of shock omission (Post-shock and Post-omission) within acquisition (left) and extinction (right) test days, respectively. **f**. Average Z-scored *mMORp*-δLight fluorescence during trace fear conditioning acquisition in male and female mice. **g**. Comparison of δLight AUC at each trial phase period between sexes on the acquisition day. **h**. Average Z-scored *mMORp*-δLight fluorescence during trace fear conditioning extinction in male and female mice. **i**. Comparison of δLight AUC at each trial phase period between sexes on the extinction day.

**Extended Data Figure 14.**
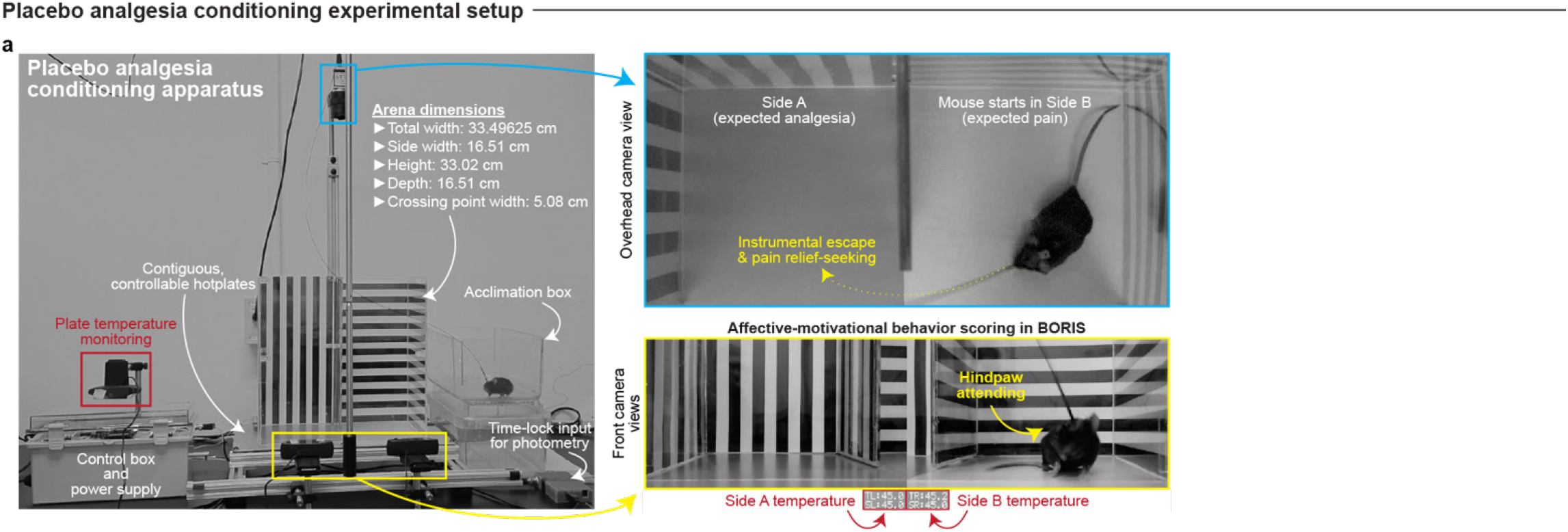
Experimental setup for placebo analgesia conditioning. **a**. Behavioral apparatus and equipment setup for placebo analgesia conditioning. The apparatus consists of two contiguous adjustable hotplate surfaces and a 2-chamber box with distinct visual cues. The hotplates are independently controlled via an Arduino microcontroller. Features of the setup are highlighted, showing overhead and front-facing camera views. Movement tracking was collected with the overhead camera and processed with Ethovision XT while nocifensive behaviors were quantified using the front-view camera in BORIS.

**Extended Data Figure 15.**
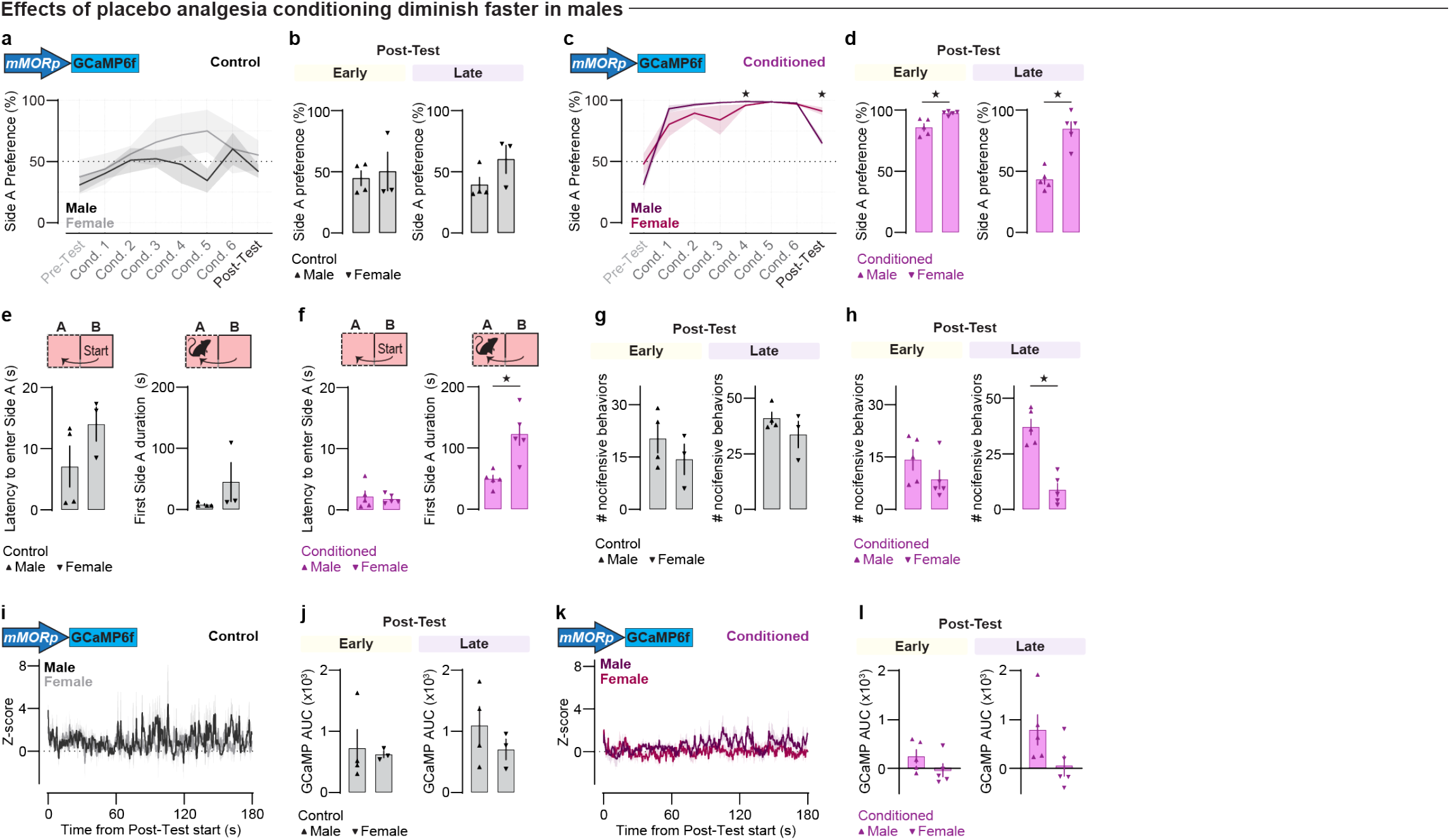
Placebo analgesia conditioning wanes in a sex-specific manner. **a**. Comparison of male and female Side A preference across test days in the *mMORp*-GCaMP6f Control group. **b**. Comparisons of Side A preference during early (left) and late (right) Post-Test phases in the Control group. **c**. Comparison of male and female Side A preference across test days in the *mMORp*-GCaMP6f Conditioned group. **d**. Comparisons of Side A preference during early (left) and late (right) Post-Test phases in the Conditioned group. **e**. Comparison of latency to first entry into Side A (left) and duration of first Side A visit (left) by sex in the Control group. **f**. Comparison of latency to first entry into Side A (left) and duration of first Side A visit (left) by sex in the Conditioned group. **g**. Comparison of total number of nocifensive behaviors observed during the early (left) and late (right) Post-Test phases by sex in the Control group. **h**. Comparison of total number of nocifensive behaviors observed during the early (left) and late (right) Post-Test phases by sex in the Conditioned group. **i**. Average Z-scored *mMORp*-GCaMP6f bulk fluorescence traces for male and female mice in the Control group during the Post-Test. **j**. Comparison of bulk GCaMP6f fluorescence during the early (left) and late (right) Post-Test by sex in the Control group. **k**. Average Z-scored *mMORp*-GCaMP6f bulk fluorescence traces for male and female mice in the Conditioned group during the Post-Test. **l**. Comparison of bulk GCaMP6f fluorescence during the early (left) and late (right) Post-Test by sex in the Conditioned group.

**Extended Data Figure 16.**
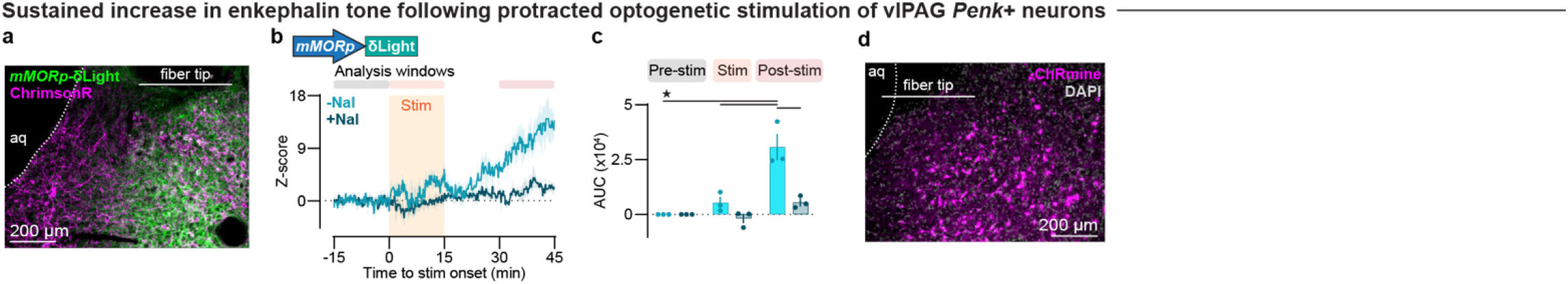
Optogenetic stimulation of *Penk*+ neurons in vlPAG promotes sustained enkephalin release. **a**. Representative 20x image of fiber optic tip location and viral expression of *mMORp*-δLight and Cre-dependent ChrimsonR in vlPAG of a Penk-Cre mouse. **b**. Average Z-scored *mMORp*-δLight traces surrounding 15 minutes of 10-Hz optogenetic stimulation in untreated and naloxone pre-treated (4.0 mg/kg, i.p.) conditions. **c**. Comparison of changes in δLight bulk fluorescence (AUC) across time windows prior to, during, and following optogenetic stimulation by drug treatment. **d**. Representative 20x image of fiber tip location and Cre-dependent ChRmine expression in a Penk-Cre mouse related to Figure 4r-s.

**Extended Data Figure 17.**
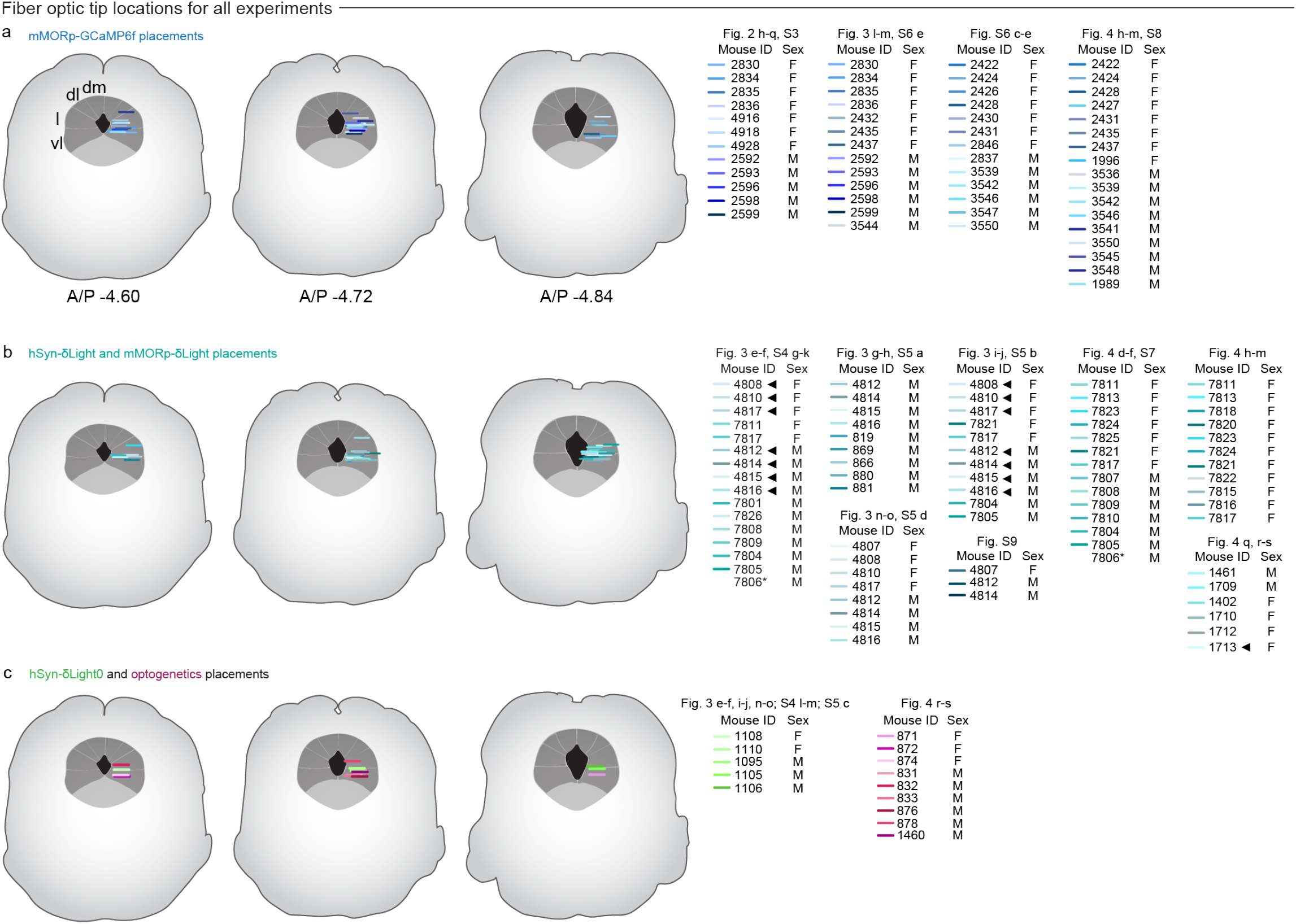
Fiber optic tip locations for all fiber photometry and optogenetics experiments. **a**. Fiber optic tip locations for mice expressing *mMORp*-GCaMP6f in vlPAG by animal number, sex, and relevant experimental figure. Animals used in multiple figure panels are shown once for clarity. **b**. Fiber optic tip locations for mice expressing either *hSyn*-δLight or *mMORp*-δLight in vlPAG. Mice denoted with an arrowhead indicate inclusion in all listed figures. The mouse denoted with a star indicates histological verification of fiber optic tip placement is unavailable. **c**. Fiber optic tip locations for mice expressing hSyn-δLight0 or red-shifted opsins (*i*.*e*., ChrimsonR or ChRmine) in vlPAG.

**Extended Data Table 1.**
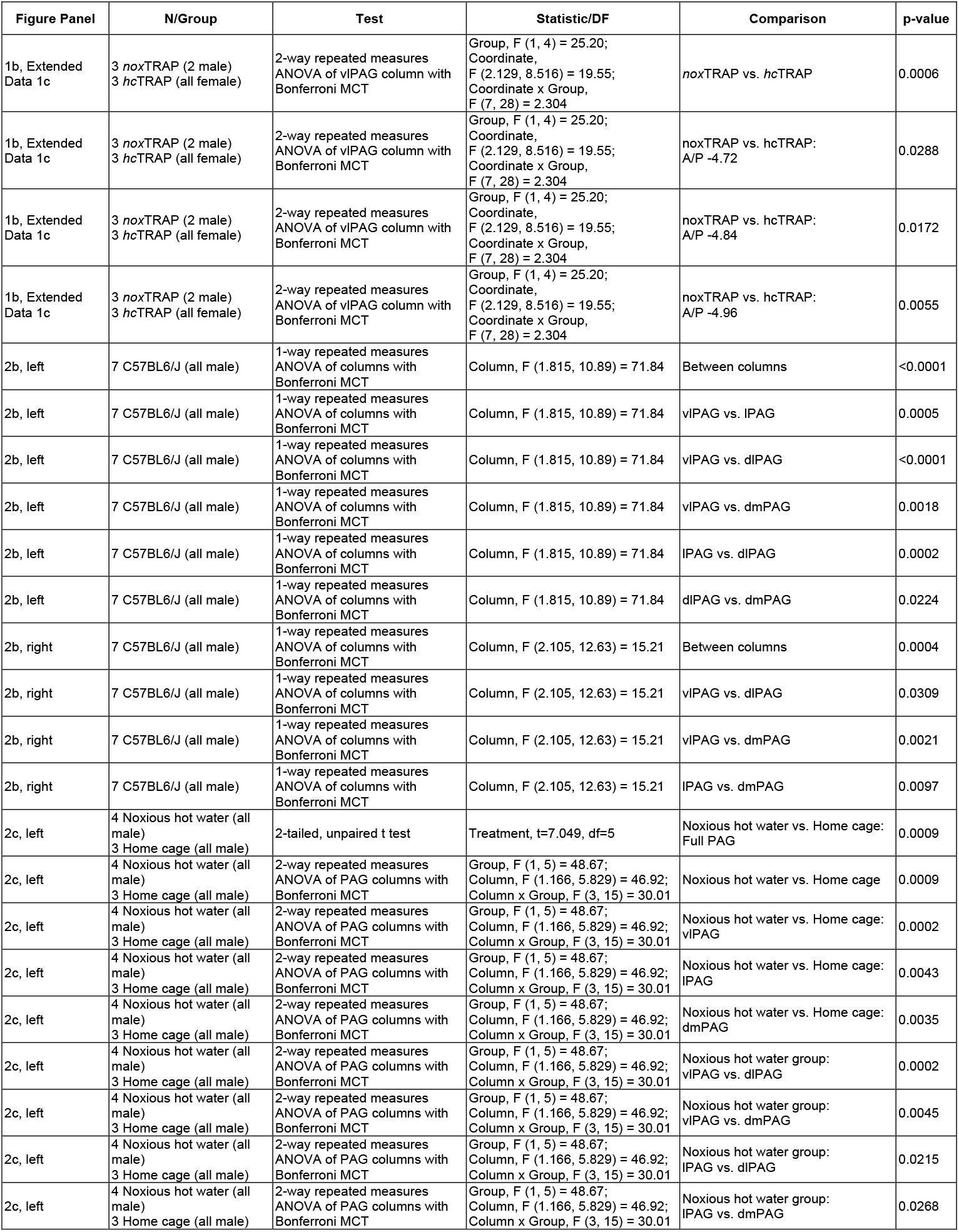
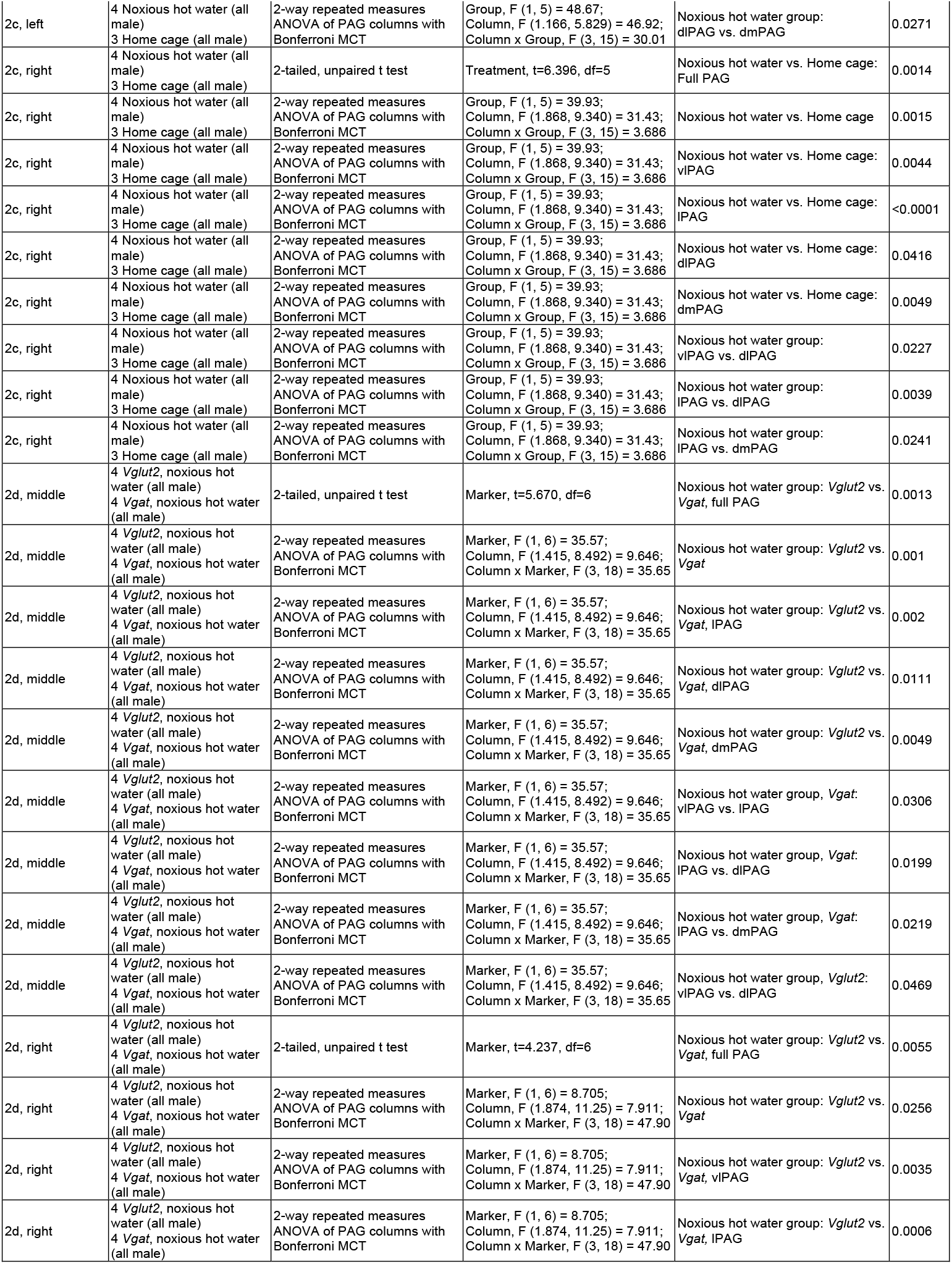
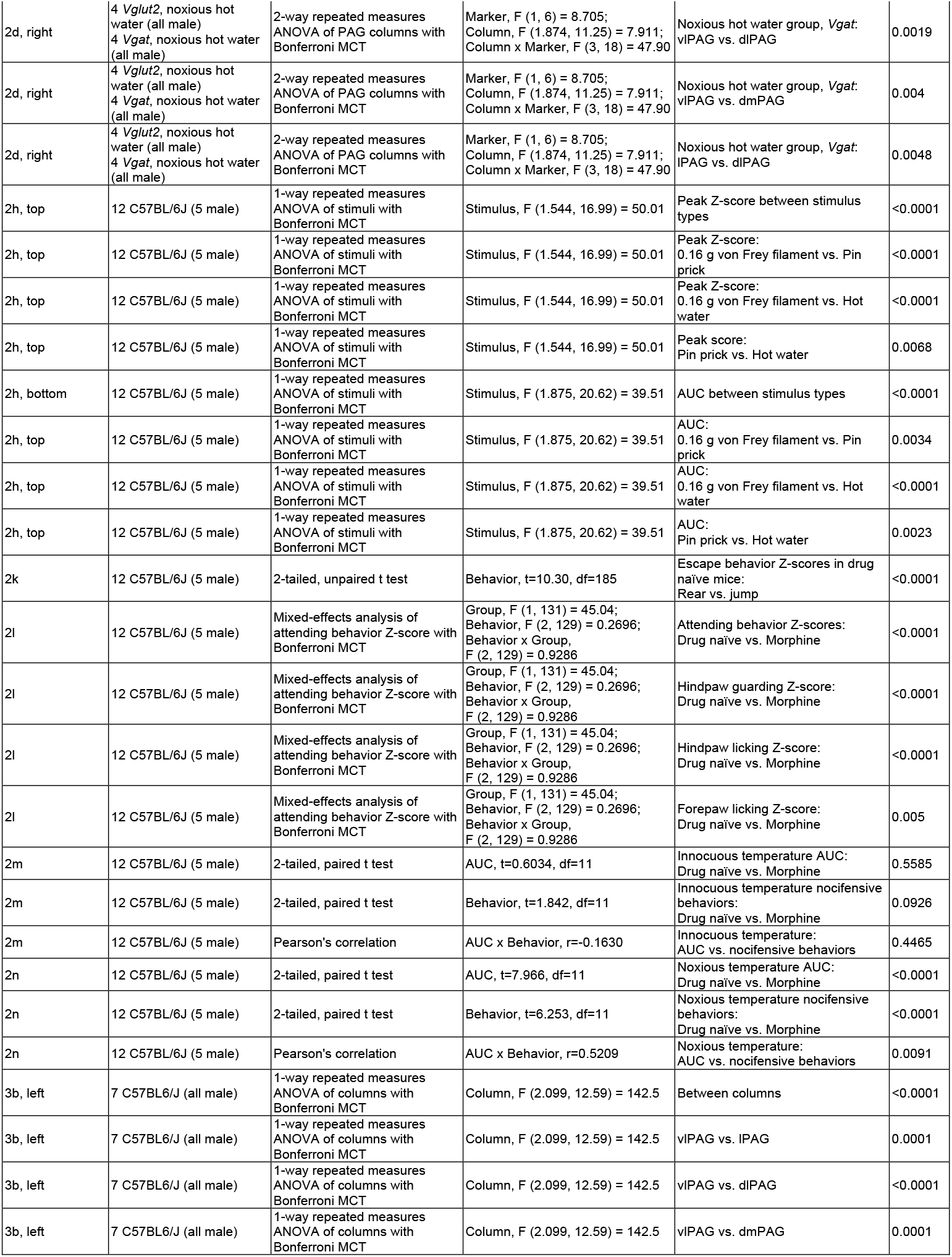
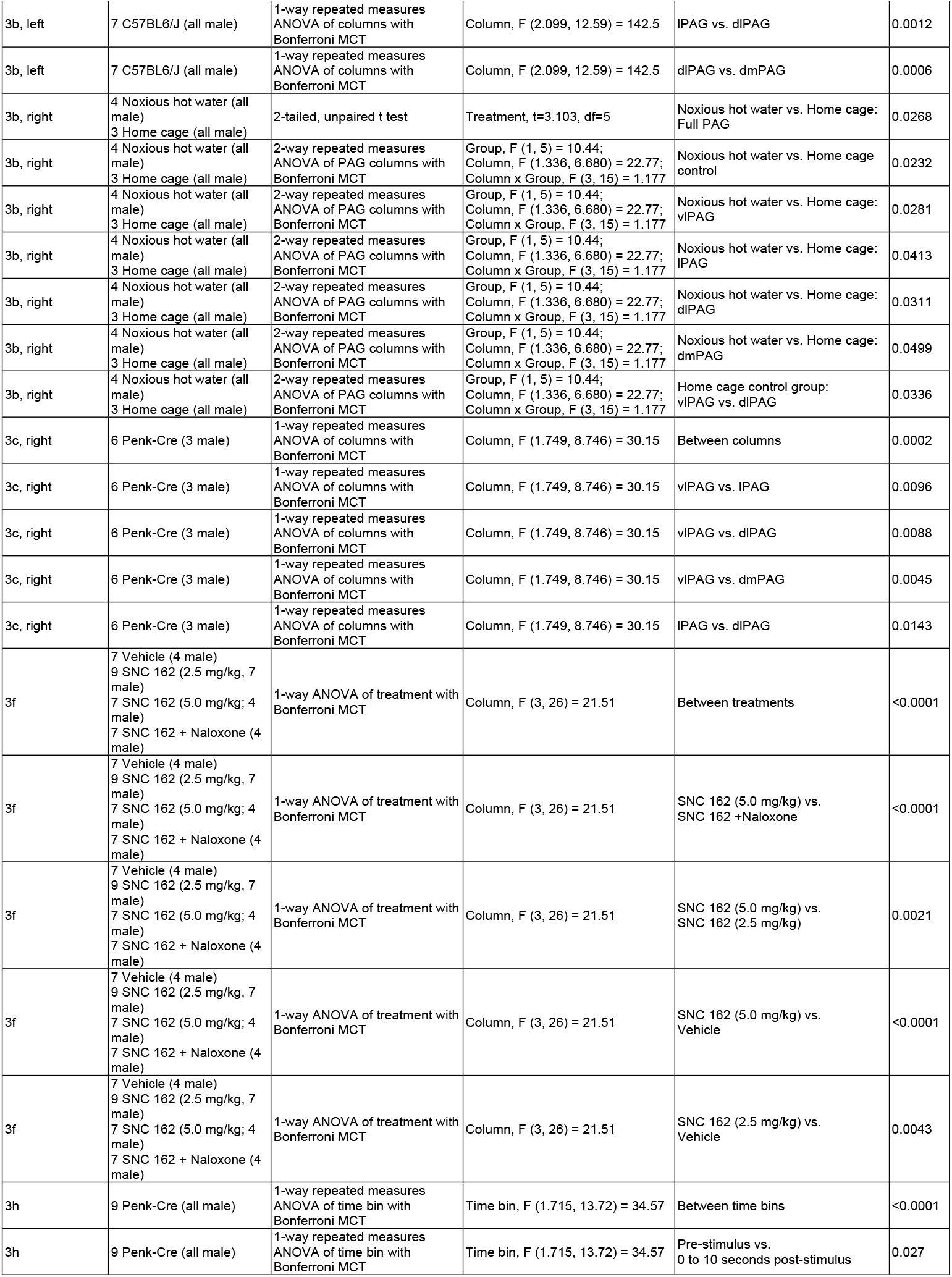
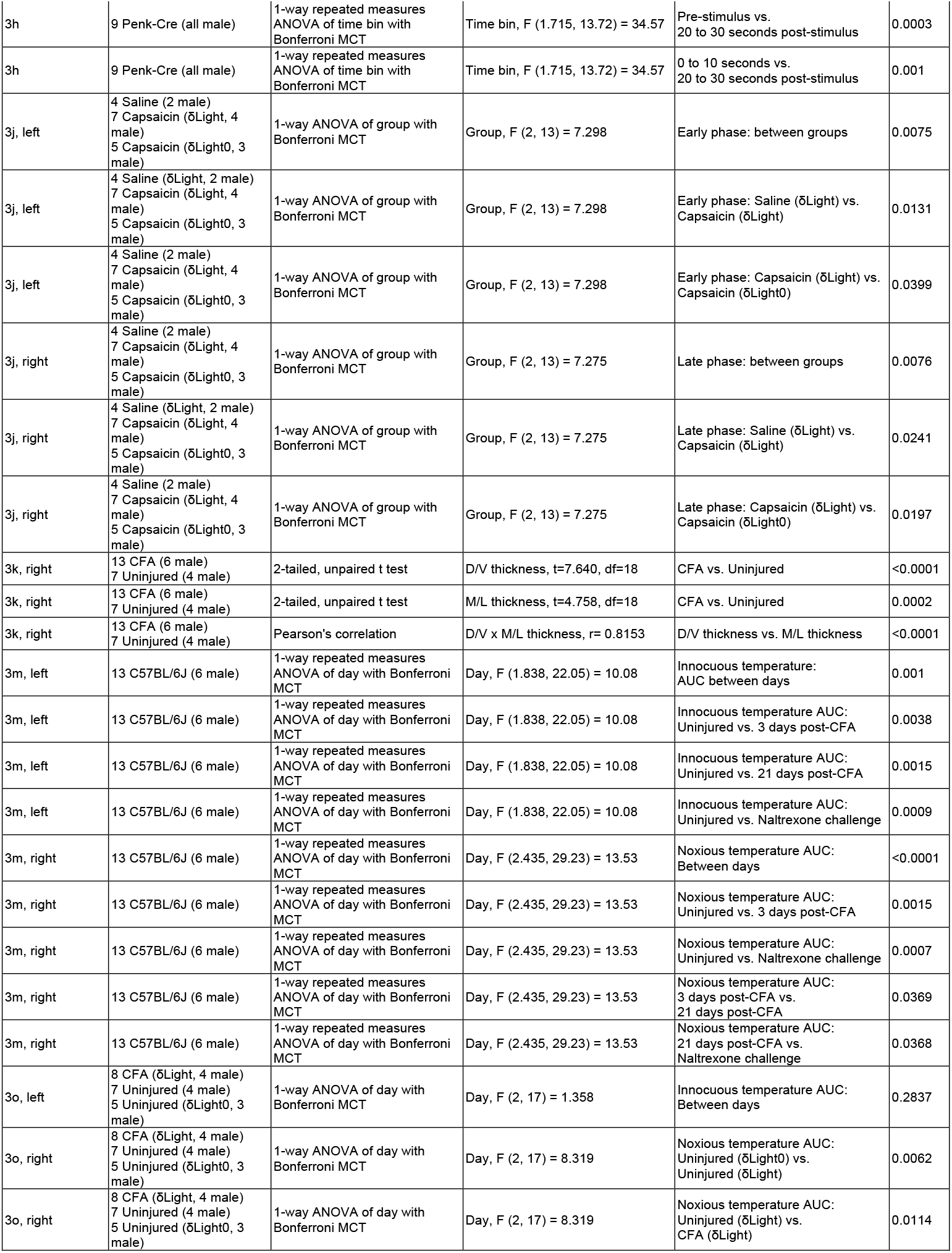
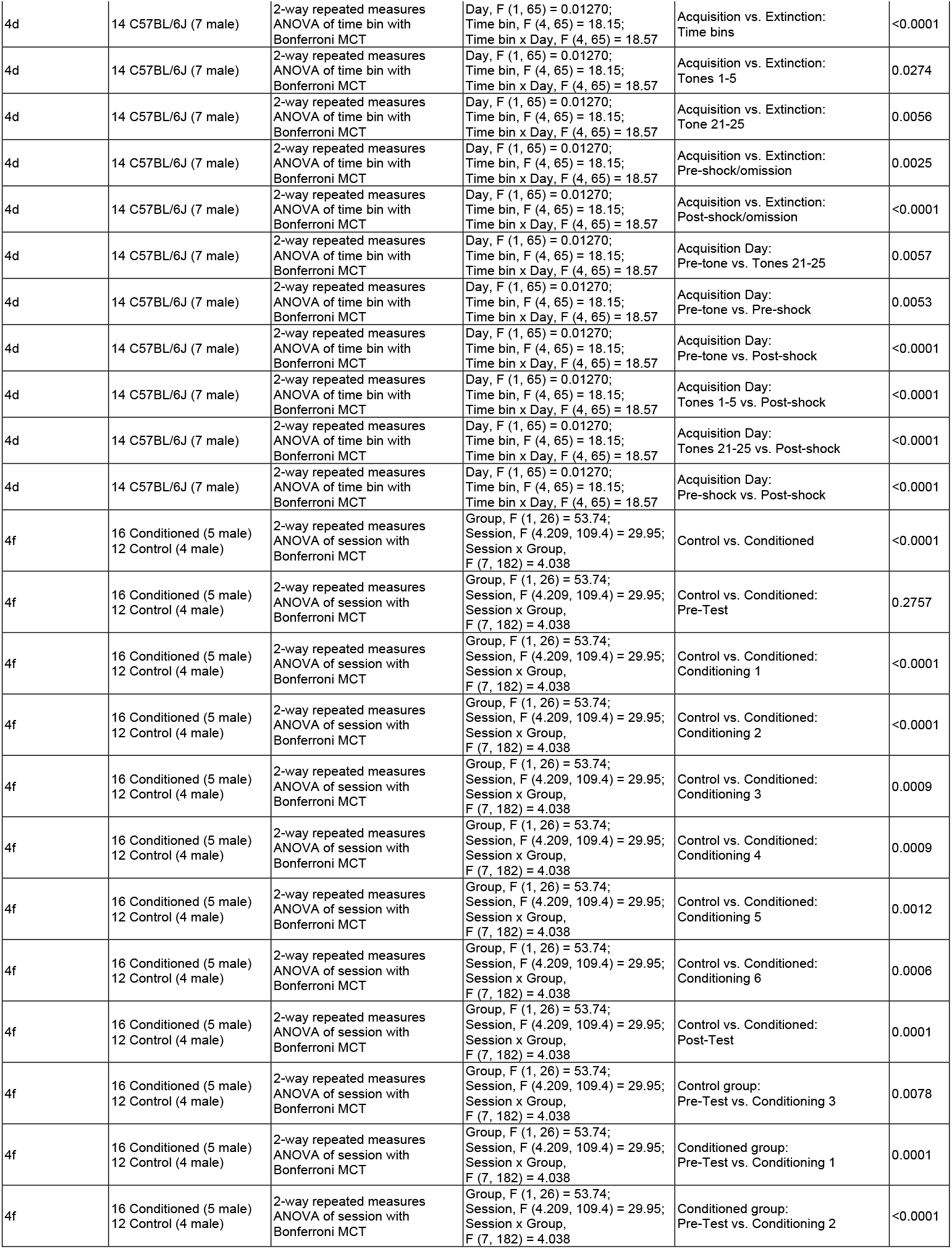
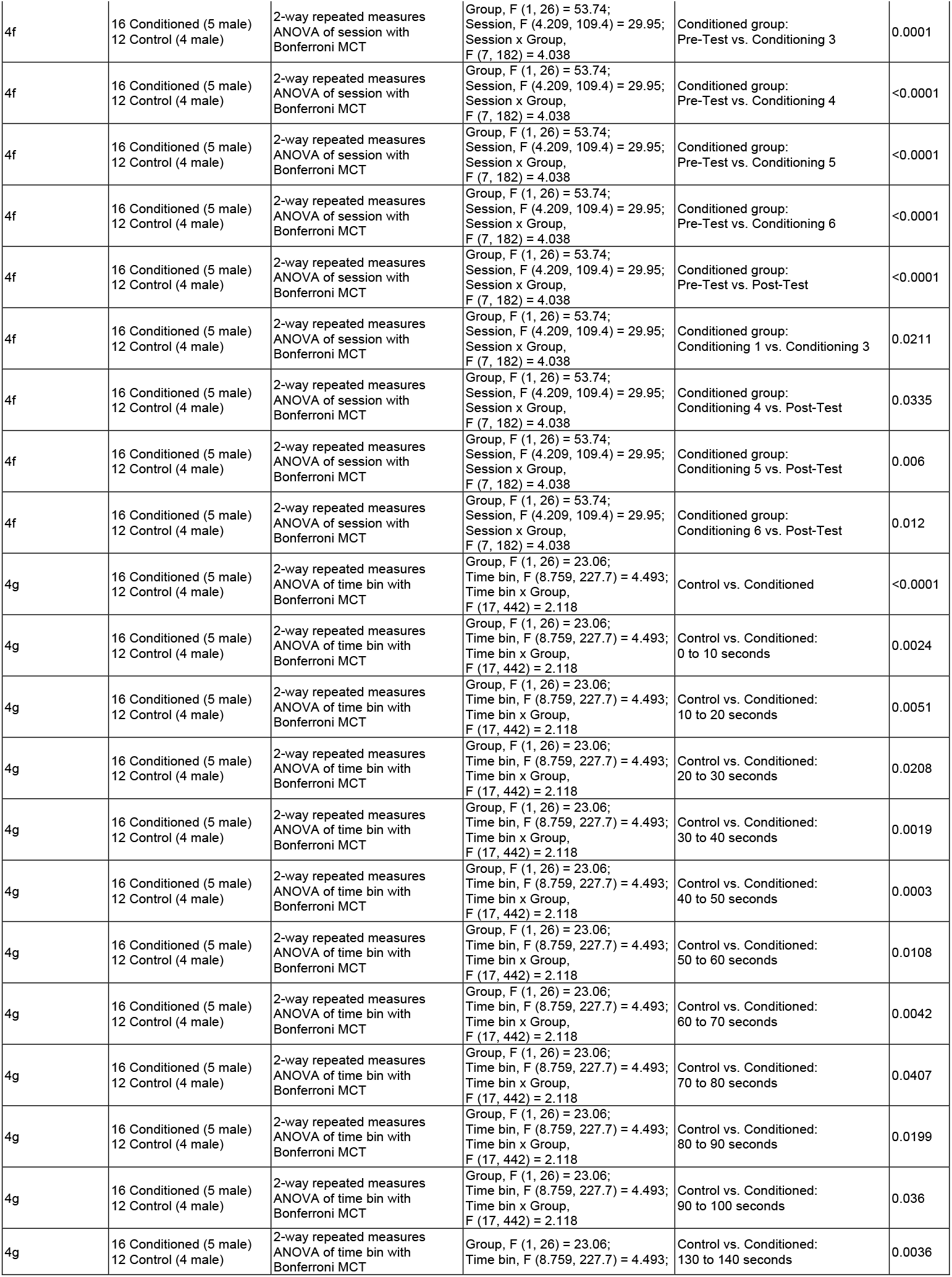
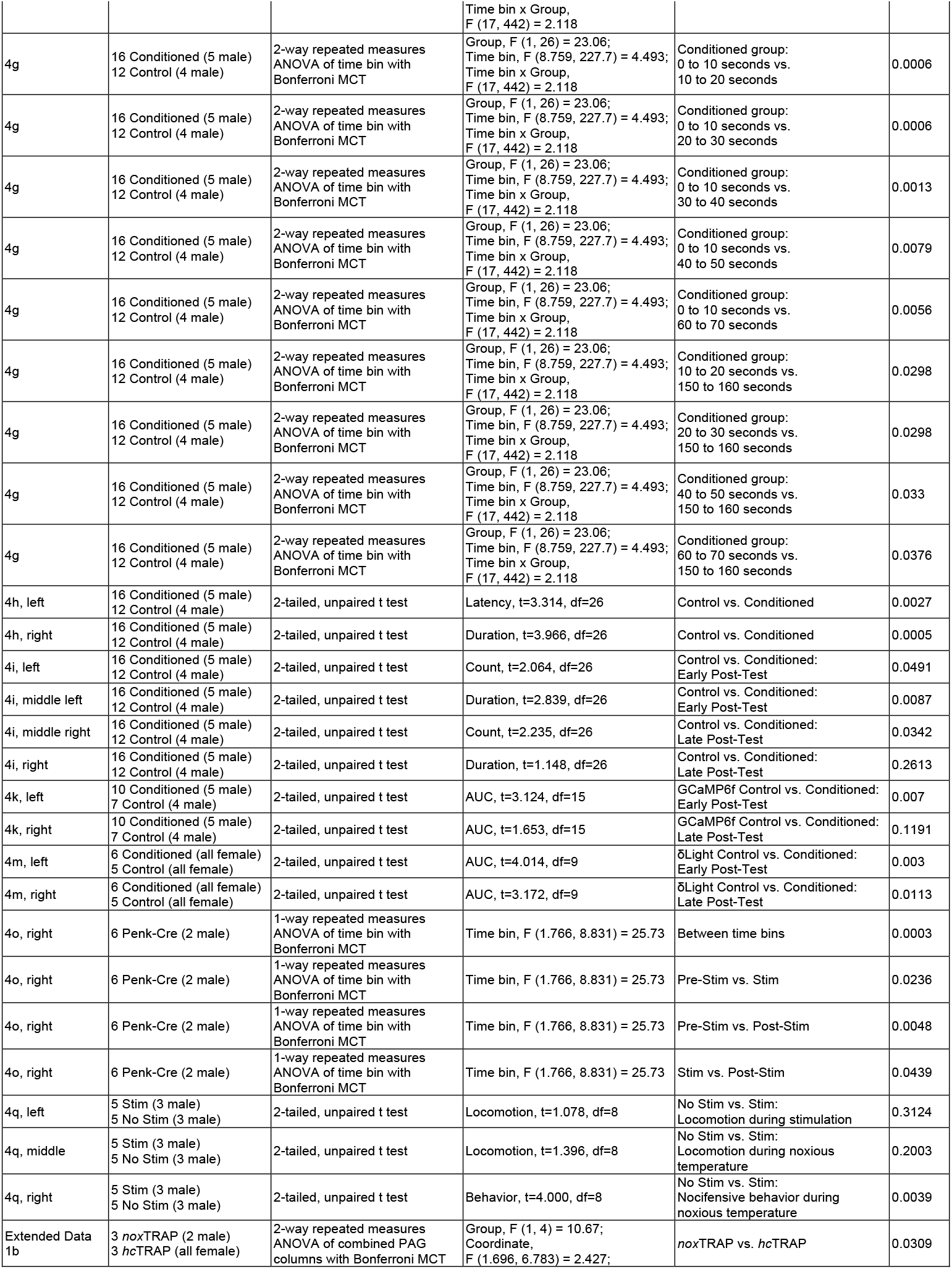
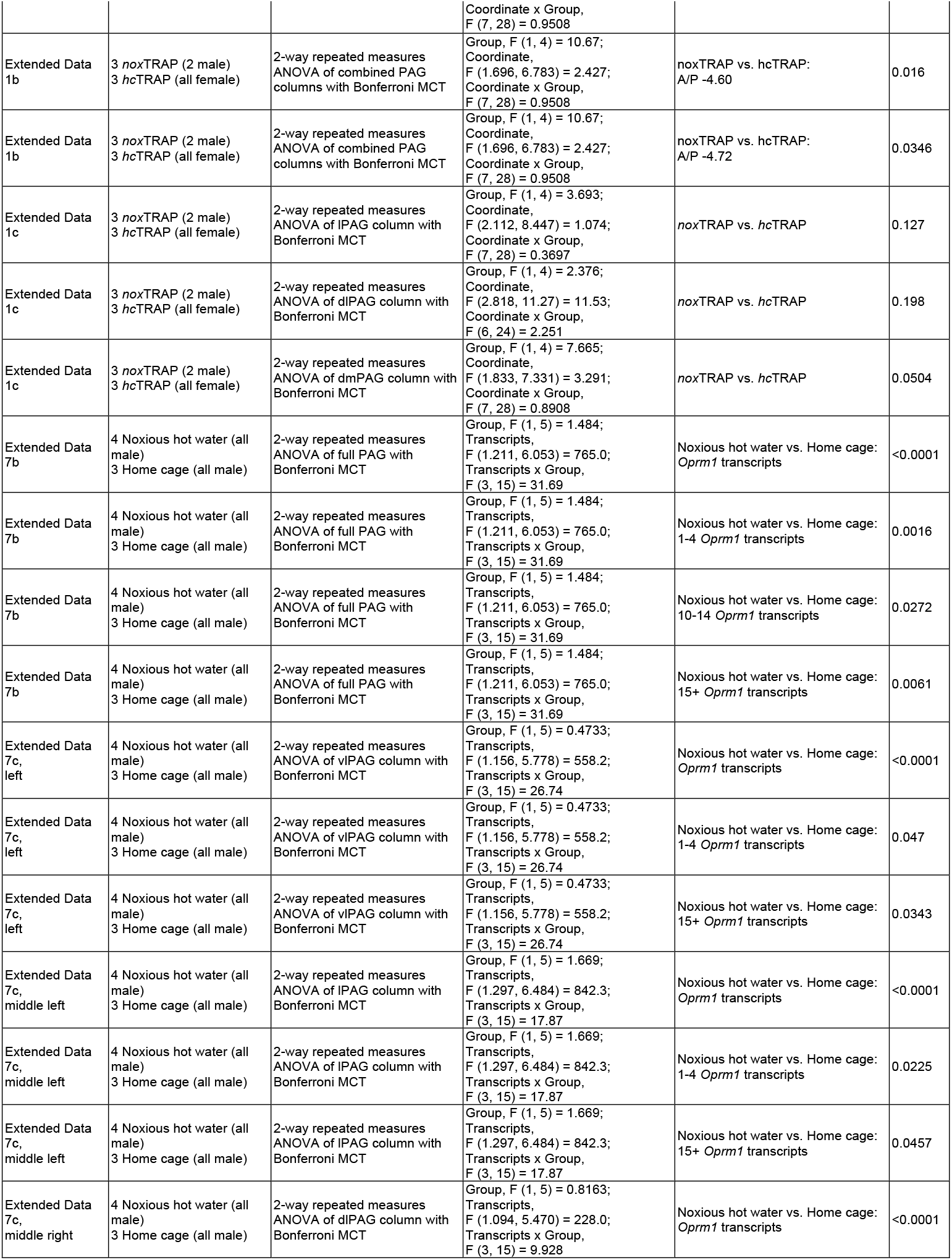
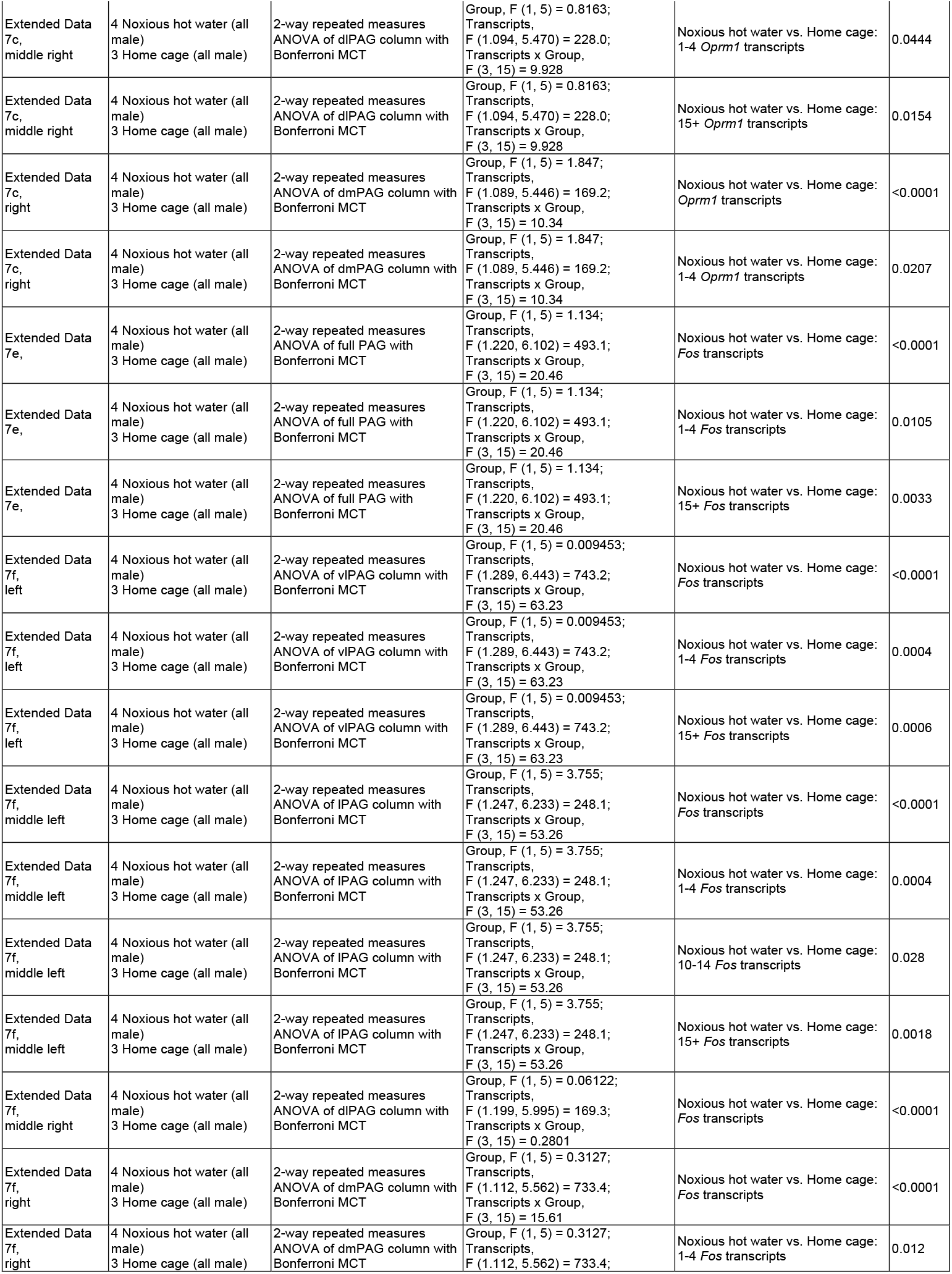
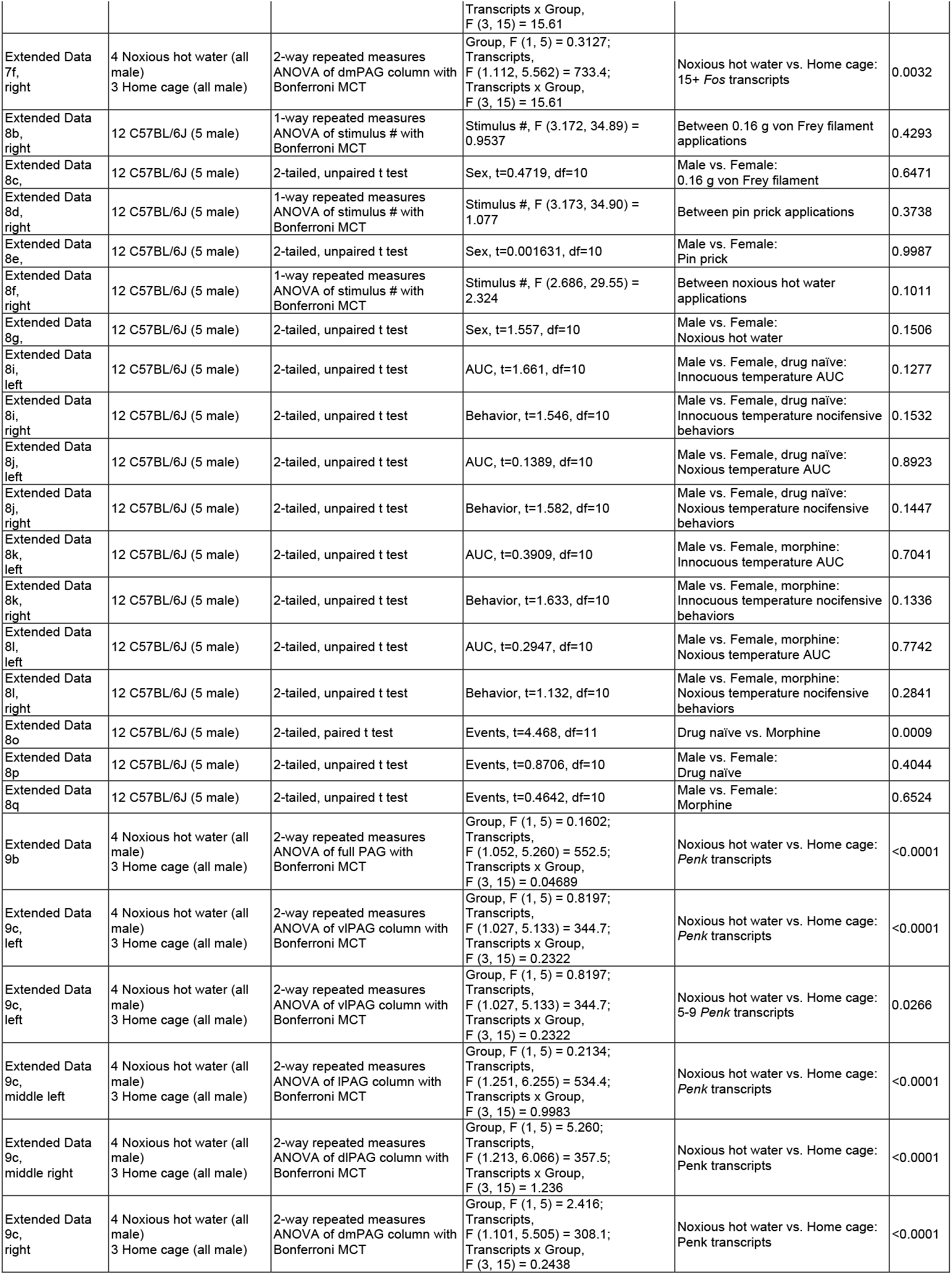
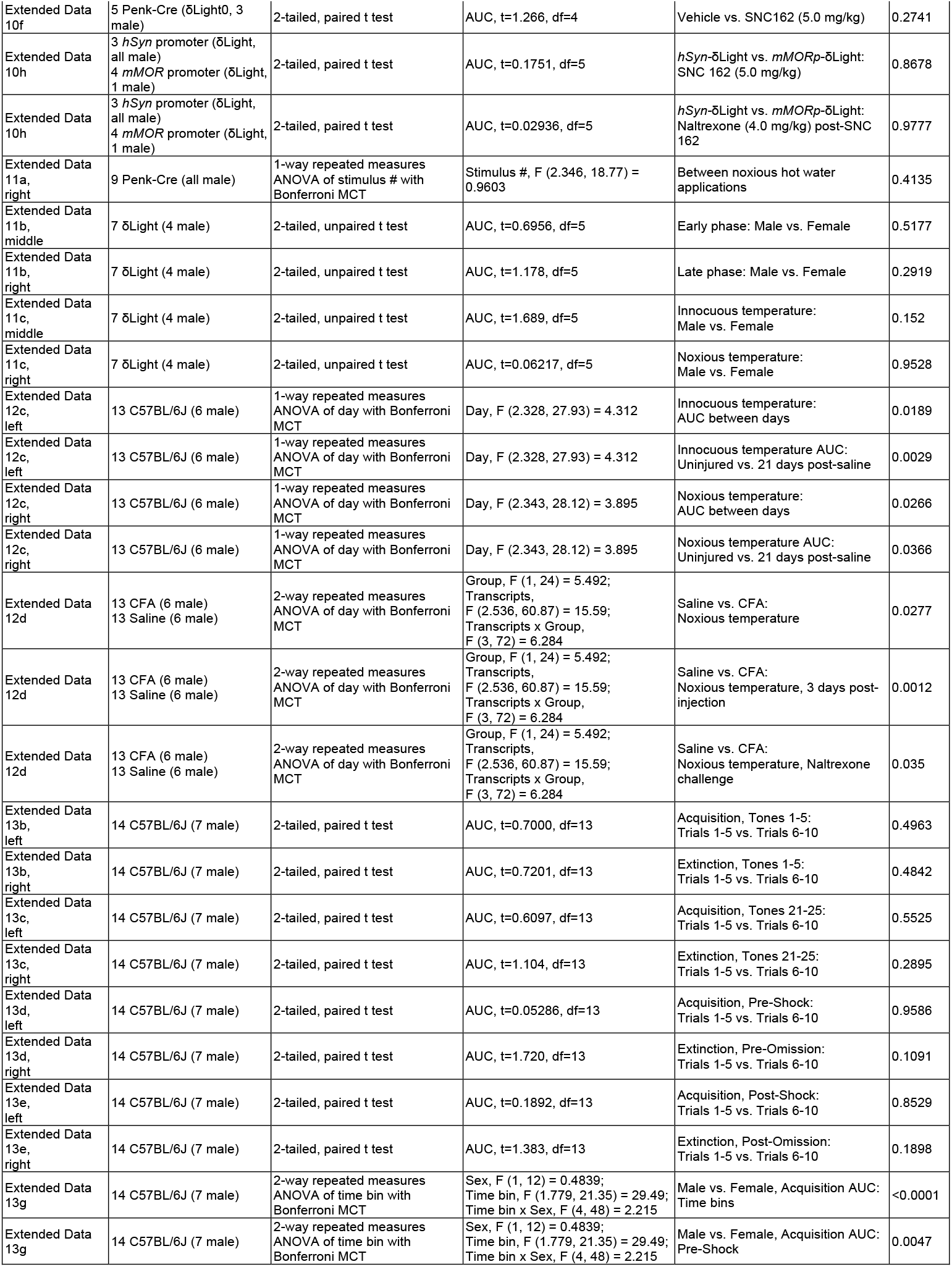
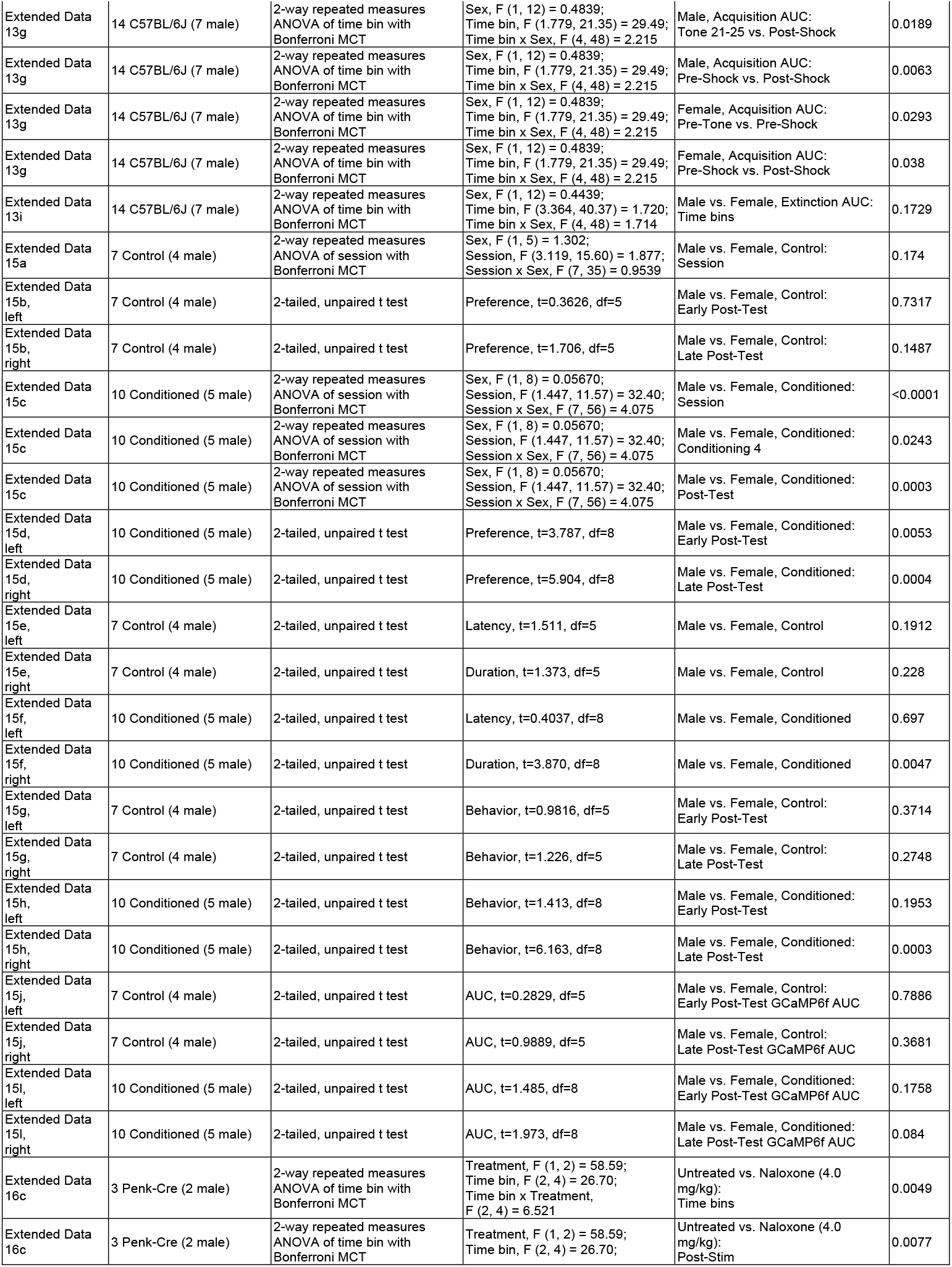
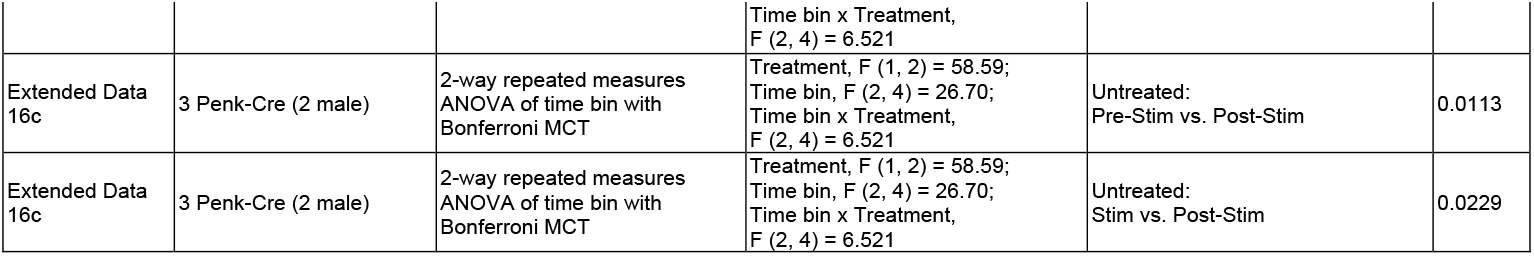
Statistical test details.

